# The Acquisition of Resistance to Carbapenem and Macrolide-mediated Quorum Sensing Inhibition by *Pseudomonas aeruginosa* via a Novel Integrative and Conjugative Element ICE_Tn4371_6385

**DOI:** 10.1101/161497

**Authors:** Yichen Ding, Jeanette Teo, Daniela I. Drautz-Moses, Stephan Christoph Schuster, Michael Givskov, Liang Yang

## Abstract

*Pseudomonas aeruginosa* can cause persistant and life-threatening infections in immunocompromised patients. Carbapenems are the first-line agents to treat *P. aeruginosa* infections; therefore, the emergence of carbapenem-resistant *P. aeruginosa* strains has greatly challenged effective antibiotic therapy. In this study, we characterised the full-length genomes of two carbapenem resistant *P. aeruginosa* clinical isolates that produce the carbapebemase New Delhi metallo-β-lactamase-1 (NDM-1). We found that the *bla*_*NDM-1*_ gene is encoded by a novel intergrative and conjugative element (ICE) ICE_Tn4371_6385, which also carries the macrolide resistance gene *msr(E)* and the florfenicol resistance gene *floR*. The *msr(E)* gene has rarely been described in *P. aeruginosa* genomes. To investigate the functional roles of *msr(E)* in *P. aeruginosa*, we exogeneously expressed this gene in *P. aeruginosa* laboratory strains and found that the acquisition of *msr(E)* could abolish the azithromycin-mediated quorum sensing inhibition *in vitro* and the anti-Pseudomonas effect of azithromycin *in vivo*. In addition, the expression of *msr(E)* almost completely restored the azithromycin-affected *P. aeruginosa* transcriptome, as shown by our RNA sequencing analysis. We present the first evidence of *bla*_*NDM-1*_ to be carried by intergrative and conjugative elements, and the first evidence of co-transfer of carbapenem resistance and the resistance to macrolide-mediated quorum sensing inhibition into *P. aeruginosa* genomes.

**Importance:** Carbapenem resistant *P. aeruginosa* has recently been listed as the top three most dangerous superbugs by World Health Organisation. The transmission of *bla*_*NDM-1*_ gene into *P. aeruginosa* can cause extreme resistance to carbapenems and fourth generation cephalosporins, which greatly compromises the effectiveness of these antibiotics against Pseudomonas infections. However, the lack of complete genome sequence of NDM-1-producing *P. aeruginosa* has limited our understanding of the transmisibility of *bla*_*NDM-1*_ in this organism. Here we showed the co-transfer of *bla*_*NDM-1*_ and *msr(E)* into *P. aeruginosa* genome by a novel integrative and conjugative element (ICE). The acquisition of these two genes confers *P. aeruginosa* with resistance to carbapenem and macrolide-mediated quorum sensing inhibition, both of which are important treatment stretagies for *P. aeruginosa* infections. Our findings highlight the potential of ICEs in transmitting carbapenem resistance, and that the anti-virulence treatment of *P. aeruginosa* infections by macrolides can be challenged by horizontal gene transfer.

## Introduction

*Pseudomonas aeruginosa* is an opportunistic pathogen which can cause life-threatening infections in immunocompromised patients (1). It is also responsible for nosocomial infections such as chronic wound infections and ventilator-associated pneumonia, and chronic airway infections in cystic fibrosis and chronic obstructive pulmonary disease patients (2-5). These infections are usually very difficult to eradicate and associated with high mortality rates (3, 6). Worse still, *P. aeruginosa* is also notorious for its ability to develop multidrug resistance, which leaves only a handful of antibiotics remain effective to treat its infections in clinical practice (7).

Carbapenems such as imipenem and meropenem are the first-line agents for treating *P. aeruginosa* infections and the last-ressort drugs in severe infections caused by Gram-negative bacteria (8). However, the clinical efficacy of carbapenems has been greatly compromised by the spreading of the carbapebemase New Delhi metallo-β-lactamase-1 (NDM-1), which is usually encoded and transmitted by broad-host self-conjugative plasmids in Enterobacteriacea spp. and *Acinetobacter baumannii* (9). The *bla*_*NDM-1*_ gene was first described in *P. aeruginosa* in 2011 and has ever since been identified in *P. aeruginosa* isolated from North America, Europe and Asia (10-16). The NDM-1-producing *P. aeruginosa* strains usually have extremely high resistance to carbapenems and many other classes of antibiotics, which makes infections caused by these superbugs even more difficult to treat (10-16). Therefore, it is important to develop novel treatment strategies for *P. aeruginosa* infections.

One alternative strategy for treating *P. aeruginosa* infections is to inhibit the production of virulence factors, which are essential for the pathogenesis of this bacterium. For instance, the type III secretion system and the secreted products such as elastase and exotoxin A have been shown to play important roles during the colonization of *P. aeruginosa* in human airways (17, 18). In particular the secreted detergent rhamnolipid causes rapid necrosis of the host immune cells and protects bacterial cells from immune attack and facilitate the establishment of infections (19). In *P. aeruginosa*, the expression of many virulence factors is under tight control of its quorum sensing (QS) systems, which can be potential targets for the design of anti-virulence drugs (20, 21). However, although several anti-QS compounds have been identified in the past, none of them has so far entered clinical trial (21-23). On the other hand, the macrolide antibiotics such as erythromycin, azithromycin (AZM) and clarithromycin were shown to have a promising anti-QS activity, which makes them ideal anti-virulence drugs for treating *P. aeruginosa* infections (21). It was suggested that macrolides repress the synthesis of QS signaling molecules by interfering with the signaling pathways of RsmZ and RsmY through yet-to-be identified targets, resulting in the downregulation of the QS-regulated virulence products such as rhamnolipids and elastase in *P. aeruginosa* (24). The use of macrolides as QS inhibitor may therefore expand our antimicrobial arsenal against *P. aeruginosa* infections until new and more efficient anti-virulence drugs become clinically available.

Although NDM-1-producing *P. aeruginosa* strains have been prevalent worldwide, the complete genome information and transmission mechanisms is still lacking (10-16). To improve the understanding of the transmission of *bla*_*NDM-1*_ among *P. aeruginosa* strains, we sequenced the complete genomes of two local clinical NDM-producing *P. aeruginosa* isolates (14). Comparative genomic analysis showed that the *bla*_*NDM-1*_ gene is encoded by a novel Tn4371 family integrative and conjugative element (ICE), which is a class of mobile genetic element present in the genomes of a broad range of β-and γ-proteobacteria (25). In addition to *bla*_*NDM-1*_, this element also carries the macrolide resistance gene *msr(E)* and the florfenicol resistance gene *floR*. We found that the acquisition of *msr(E)* by *P. aeruginosa* abolished the AZM-mediated QS inhibition *in vitro* and the anti-Pseudomonas effect of AZM *in vivo*. To our knowledge, this is the first description of *bla*_*NDM-1*_ encoded by ICEs and the first evidence on the co-acquisition of carbapenem resistance and the resistance to AZM-mediated QS inhibition by *P. aeruginosa*.

## Results

### Identification of a novel NDM-1-producing *P. aeruginosa* group

A nosocomial outbreak of NDM-1-producing *P. aeruginosa* was previously reported in a local Singapore hospital and was the first case report of NDM-1-producing *P. aeruginosa* in Southeast Asia (14). The strains isolated during this outbreak exhibited multidrug resistance to carbapenems, cephalosporins, aminoglycosides, and fluoroquinolones, whereas remained sensitive to polymyxin B (Table S1). To identify the origin of these NDM-1-producing *P. aeruginosa* isolates, we sequenced the draft genomes of 11 isolated strains on an Illumina MiSeq platform and found that all these isolates belong to multilocus sequence type ST308 and have harbored the same sets of antibiotic resistance genes (Fig. S1). In addition, the 12 genomes are highly similar to each other as shown by multiple genome alignment using Progressive Mauve (26) (Fig. S2). These results suggested that the 12 NDM-1-producing *P. aeruginosa* strains isolated in this outbreak belong to the same phylogenetic group. Therefore, we named this closely related *P. aeruginosa* group PASGNDM and selected two representative isolates, PASGNDM345 and PASGNDM699 for further investigation.

To better understand the detailed features of the PASGNDM genomes and the transmission of *bla*_*NDM-1*_ into *P. aeruginosa* genome, we further sequenced PASGNDM345 and PASGNDM699 genomes on a Pacific Biosciences RSII platform. The PacBio sequencing reads have achieved 163-and 161-fold coverage for PASGNDM345 and PASGNDM699 genomes, respectively, and were successfully assembled into two full-length genomes. Construction of a phylogenetic tree using PASGNDM345 and PASGNDM699 genomes together with other 21 *P. aeruginosa* full-length genomes showed that the two PASGNDM strains formed a monophyletic group (Fig. 1). Interestingly, the closest genome to the PASGNDM group in the phylogenetic tree is PA_D1 (NZ_CP012585.1), which is an endemic *P. aeruginosa* strain causing ventilator-associated pneumonia in China as identified in previous work by us (manuscript submitted).

**Figure 1:**
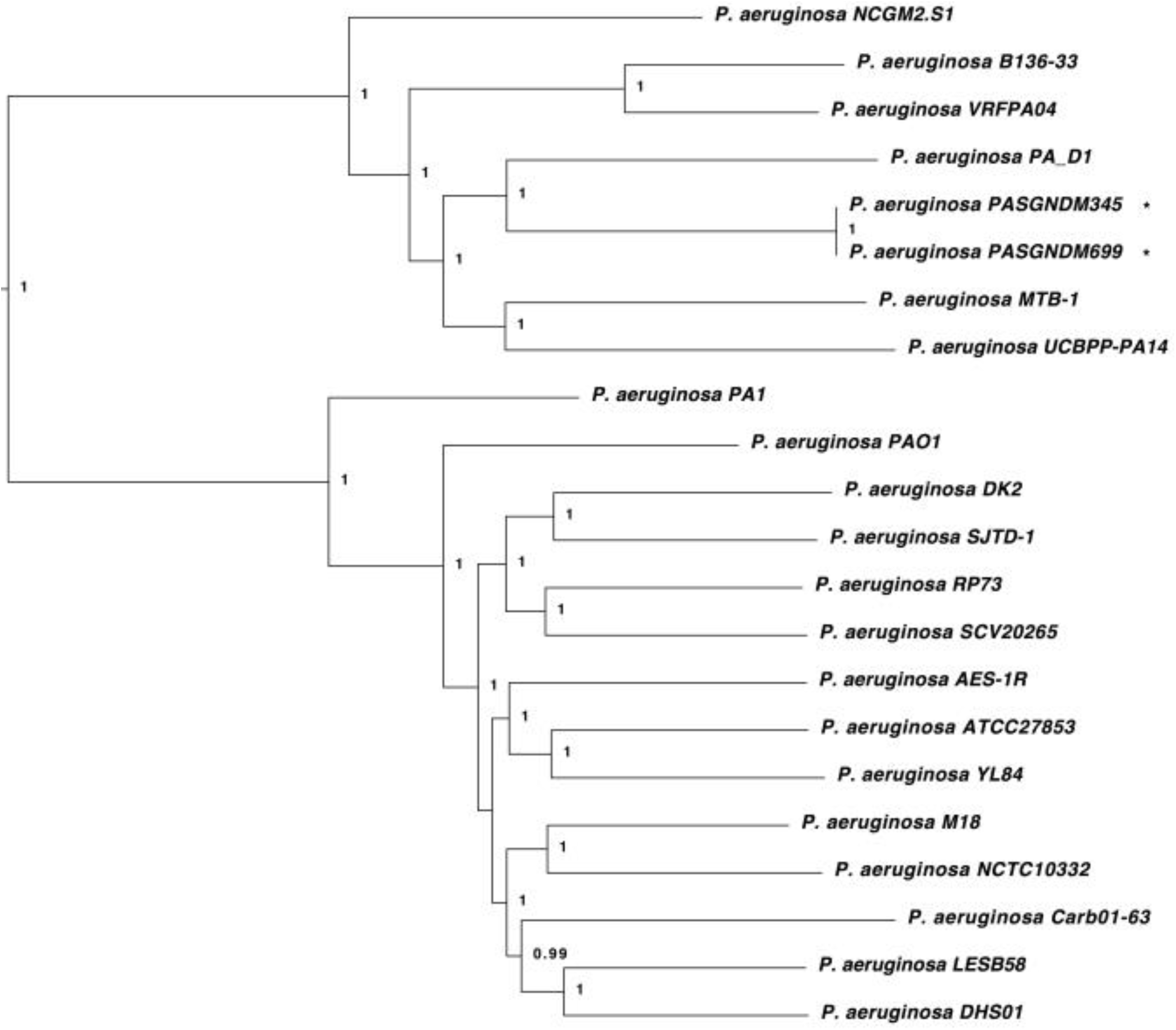
Phylogenetic tree of PASGNDM345 and PASGNDM699 with 21 *P. aeruginosa* genomes. PASGNDM345 and PASGNDM699 genomes sequenced in this study (indicated with *) was compared with 21 other *P. aeruginosa* genomes. Phylogenetic tree was constructed based on 64,240 variant sites using the approximate maximum likelihood algorithm, with clade confidence estimated with SH-like support values. Accession numbers of strains in the phylogenetic tree are listed in Table S7.

### Characterisation of PASGNDM345 and PASGNDM699 genomes

The genome of PASGNDM345 consists of a circular chromosome of 6,893,164 bp with an average GC content of 66.1%, whereas the PASGNDM699 genome is 6,985,102 bp with an average GC content of 66.0%. In total, 6,503 and 6,589 genes were predicted from PASGNDM345 and PASGNDM699 genomes, respectively. General features of these predicted genes can be found in Table S2.

To identify the strain specific genomic regions in the PASGNDM genomes, the complete genomes of PASGNDM699 and PASGNDM345 were compared with six other strains clustered in the same clade in the phylogenetic tree (Fig. 1). Genome alignment result showed that the two PASGNDM genomes possess several regions with low sequence identity to the other six strains (Fig. 2). Furthermore, we also predicted the genomic islands (GIs) in the two PASGNDM genomes using Island Viewer 3 server (27). A total of 41 and 47 GIs were predicted from the PASGNDM345 and PASGNDM699 genomes, respectively, which correlate well with the strain specific regions in the PASGNDM genome (Fig. 2 and Table S3, S4). The genes located in the GIs are mostly enriched in generating transposons, efflux pumps and multidrug resistance (Table S3, S4), which may be important for the survival and nosocomial spread of the PASGNDM strains. In addition, eight acquired antibiotic resistance genes including *bla*_*NDM-1*_ are embedded in the GIs of PASGNDM699 and PASGNDM345 genomes (Fig. 2), suggesting the importance of mobile genetic elements in the acquisition of antibiotic resistance by the PASGNDM isolates.

**Figure 2:**
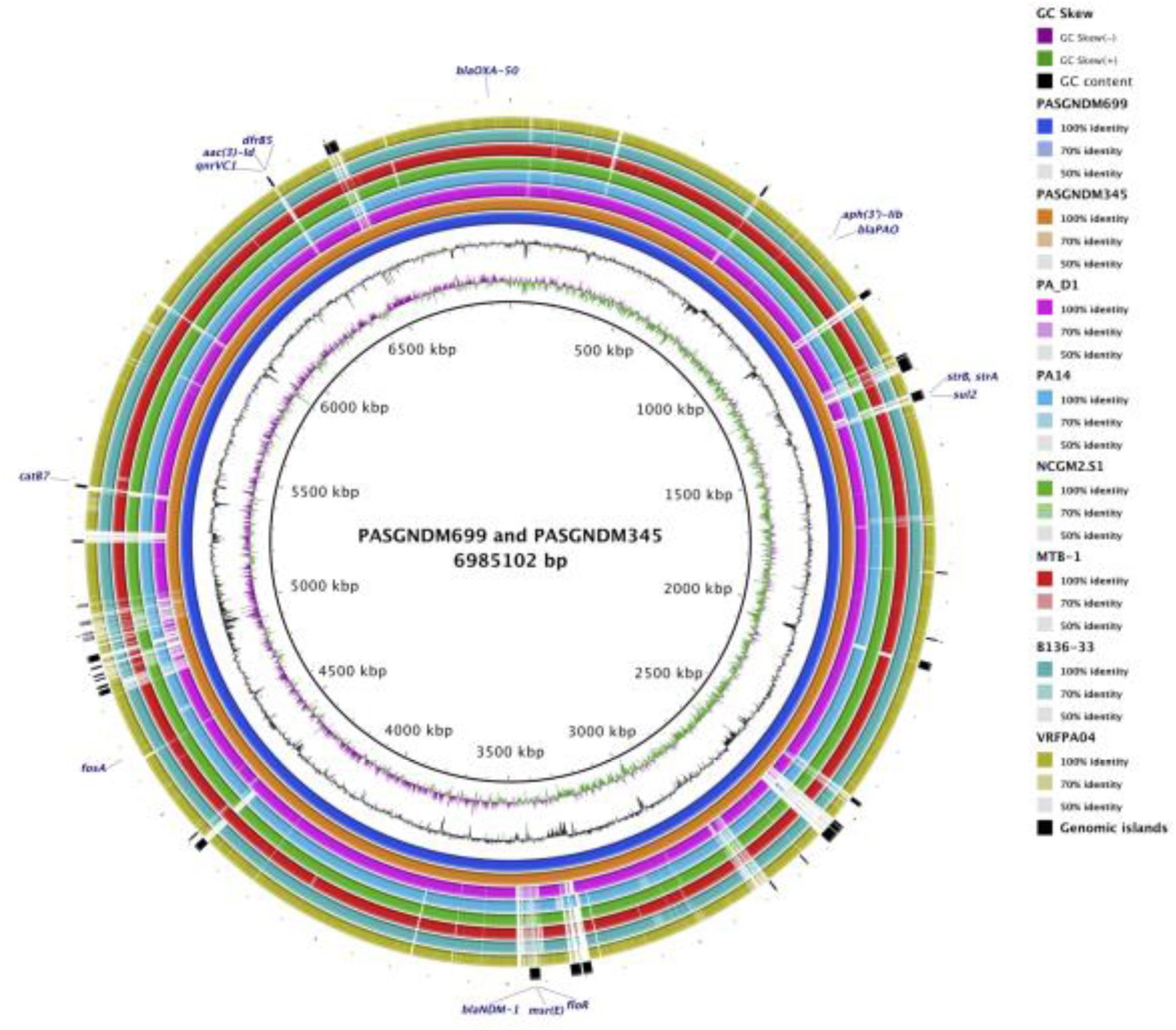
Sequence conservation between PA_D1 and 5 other *P. aeruginosa* genomes. From the innermost to outermost: Circle 1, PASGNDM699; Circle 2, PASGNDM345; Circle 3, PA_D1; Circle 4, PA14; Circle 5, DK2, NCGM2.S1; Circle 6, MTB-1; Circle 7, B136-33; Circle 8, VRFPA04. Circle 9, GIs present in PASGNDM699 and PASGNDM345 predicted by the Island Viewer3 server (27). Blocks in black indicate GIs present in both PASGNDM699 and PASGNDM345, whereas blocks in grey indicate GIs present only in PASGNDM699. Circle 10, predicted antibiotic resistance genes present in PASGNDM699 and PASGNDM345 genomes with the ResFinder 2.1 server (55). Accession numbers of the strains are listed in Table S5, and GIs are listed in Table S3 and S4.

### Identification of a novel ICE_Tn4371_6385 encoding *bla*_*NDM-1*_, *msr(E)* and *floR*

It was noted that the *bla*_*NDM-1*_ is clustered together with *msr(E)* and *floR* in a GI region, suggesting the three antibiotic resistance genes might be co-transferred into the PASGNDM genomes (Fig. 2). Further sequence analysis revealed that the three genes are embedded in a 74.2 kb ICE-like element located between the *exoY* (PA2191) and *hcnA* (PA2193) genes in the PASGNDM345 and PASGNDM699 genomes (Fig. 3). Both ends of this element are flanked by a 5’-TTTTTTGT-3’ sequence, which resembles the conserved *attB* site of almost all Tn4371 family ICEs (25). It also contains the core genes conserved among the Tn4371 family ICEs, including a *int* integrase gene, the *parB, repA* and *parA* genes of the ICE stabilisation system, and homologues to the DNA conjugative transfer machineries such as *traI* and *traG* (25) (Fig. 3). In addition, the integrase encoded by the *int* gene shared 71% identity with Int_Tn4371_ (AJ536756), which further proved that the ICE identified here should be considered as a member of the Tn4371 family, as suggested by a previous study (25). We therefore named this element ICE_Tn4371_6385 following the nomenclature system proposed by Roberts *et al.* (28).

**Figure 3:**
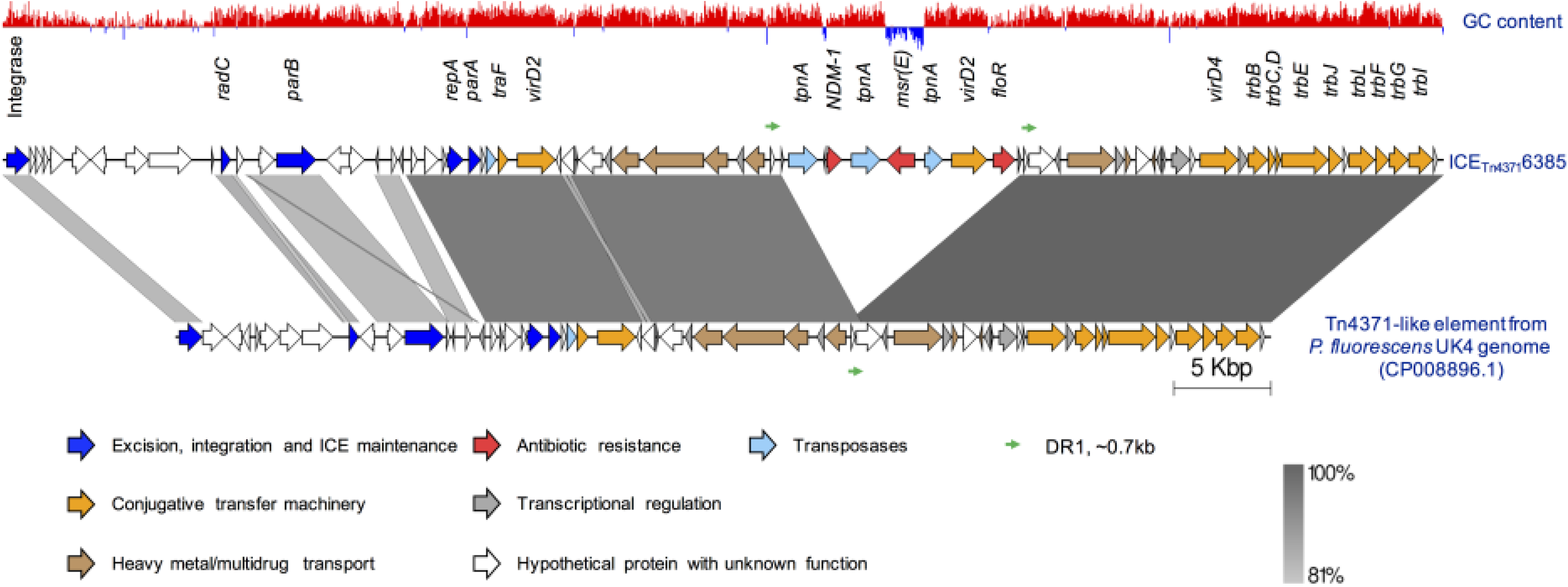
Comparison between ICE_Tn4371_6385 and the ICE-like element from *P. fluorescens* UK4 genome (CP008896.1). The 74 genes encoded by ICE_Tn4371_6385 are represented by arrows with different colors representing their functional classes. The length of each arrow is to the scale of the gene size, whereas the arrow direction indicates the transcriptional direction of the gene. The 13.7 kb segments containing three antibiotic resistance genes are between two DRs (thin green arrows) in ICE_Tn4371_6385, possibly due to a homologous recombination event. The shades between the two elements indicate the sequence identity of the linked regions.

To track the origin of ICE_Tn4371_6385, we searched its entire sequence against Genbank and found that ICE_Tn4371_6385 is similar to a 56.4 kb ICE-like element present in *Pseudomonas fluorescens* UK4 genome (CP008896.1). Comparative sequence analysis between the two elements showed that they shared a common Tn4371 ICE core gene scaffolds (25), whereas their major differences are the unique accessory gene cluster downstream of the *int* gene, and a 13.7 kb segment encoding the three antibiotic resistance genes harbored by ICE_Tn4371_6385 (Fig. 3). This 13.7 kb segment is immediately flanked by 695 bp direct repeats (termed DR), which share 99.87% identity (694/695) (Fig. 3). Interestingly, only a single copy of this DR sequence was present in the ICE-like element of the *P. fluorescens* UK4 genome (Fig. 3). It is possible that the duplicated DR sequences in ICE_Tn4371_6385 were the result of homologous recombination, which might lead to the acquisition of the 13.7 kb segment by ICE_Tn4371_6385. Therefore, we present a novel ICE_Tn4371_6385 element identified from PASGNDM genomes, which have acquired three antibiotic resistance genes namely *bla*_*NDM-1*_, *msr(E)* and *floR*. The acquisition of the *bla*_*NDM-1*_ gene by PASGNDM699 is probably responsible for its extreme resistance to carbapenems (Table S1), whereas *msr(E)* and *floR* are associated with macrolides and florfenicol resistance, respectively (29, 30).

### Acquisition of *msr(E)* protects *P. aeruginosa* from AZM-mediated QS inhibition

Previous studies have reported that sub-MIC concentrations of AZM could supress the expression of several QS-regulated virulence factors such as elastase and rhamnolipids and the swarming motility of *P. aeruginosa*, probably by inhibiting its QS systems (24, 31, 32). We therefore hypothesized that the acquisition of *msr(E)* by *P. aeruginosa* could counteract the QS-inhibition effect of AZM. However, the investigation on the functions of *msr(E)* gene in the PASGNDM strains by a reductionist approach is difficult due to their multidrug resistance; hence, an alternative approach by exogenously expressing *msr(E)* gene in laboratory strains of *P. aeruginosa* was adopted for this purpose. We first amplified the *msr(E)* gene together with its putative promoter from PASGNDM699 genome and have the entire fragment inserted into a *pUCP18* vector to construct *pUCP18*::*msr(E)* plasmid, in which the expression of *msr(E)* gene is sorely controlled under its putative promoter. Transformation of *pUCP18*::*msr(E)* into the *P. aeruginosa* strains PAO1 and PA14 increased their MICs to the AZM by more than 8 fold (from 256 μg/ml to >2048 μg/ml), indicating that the acquired *msr(E)* in PASGNDM genomes can be expressed from its own promoter to enhance resistance to AZM in *P. aeruginosa*. Both the PASGNDM699 and PASGNDM345 have comparable AZM resistance levels to the two transformed strains, suggesting that the encoded *msr(E)* is expressed in the PASGNDM strains.

We then performed elastase and rhamnolipid quantification and swarming motility assays with PAO1/*pUCP18*::*msr(E)* and PA14*/pUCP18*::*msr(E)* strains. The results showed that 8 μg/ml of AZM (1/32 of the MICs of PAO1 and PAO1/*pUCP18*) have reduced elastase production in both wildtype PAO1 and vector-carrying strain PAO1/*pUCP18* by at-least 40%, whereas the elastase produced by PAO1/*pUCP18*::*msr(E)* was not significantly affected upon AZM treatment (Fig. 4a). We also found that AZM have reduced the amount of rhamnolipid in the supernatant of both PAO1 and PAO1/*pUCP18* overnight culture by at-least 60%, whereas PAO1/*pUCP18*::*msr(E)* produced similar levels of rhamnolipid in the presence and absence of AZM (Fig. 4b). In addition, the swarming motilities of PAO1 and PAO1/*pUCP18* were significantly inhibited by 8 μg/ml of AZM, under which PAO1/*pUCP18*::*msr(E)* exhibited a normal swarming phenotype (Fig. 5). Similar results were also obtained using the PA14 strains (Fig. 4c and 4d, Fig. 5), suggesting that the anti-AZM-mediated QS-inhibition effect of Msr(E) is not strain-specific. Taken together, these results clearly showed that the acquisition of *msr(E)* could protect *P. aeruginosa* from AZM-mediated QS inhibition.

**Figure 4:**
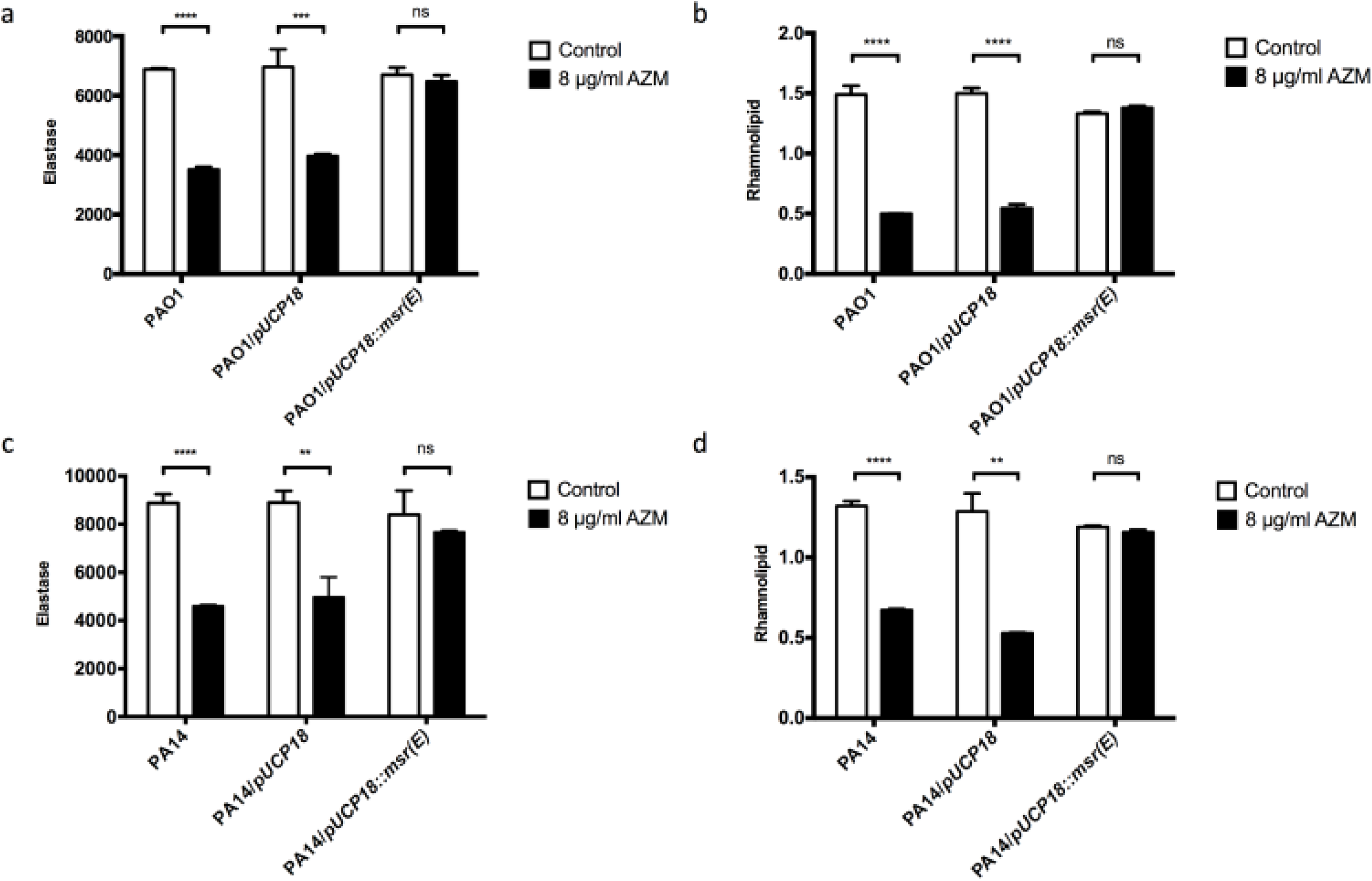
Impact of exogenous expression of Msr(E) on elastase and rhamnolipid production by PAO1 and PA14 under 8 μg/ml of AZM. The production of elastase and rhamnolipid was inhibited in wildtype strains and vector-carrying strains of PAO1 (a), (b) and PA14 (c), (d) under the treatment of 8 μg/ml AZM, whereas no inhibition was observed in the strains expressing Msr(E).

**Figure 5:**
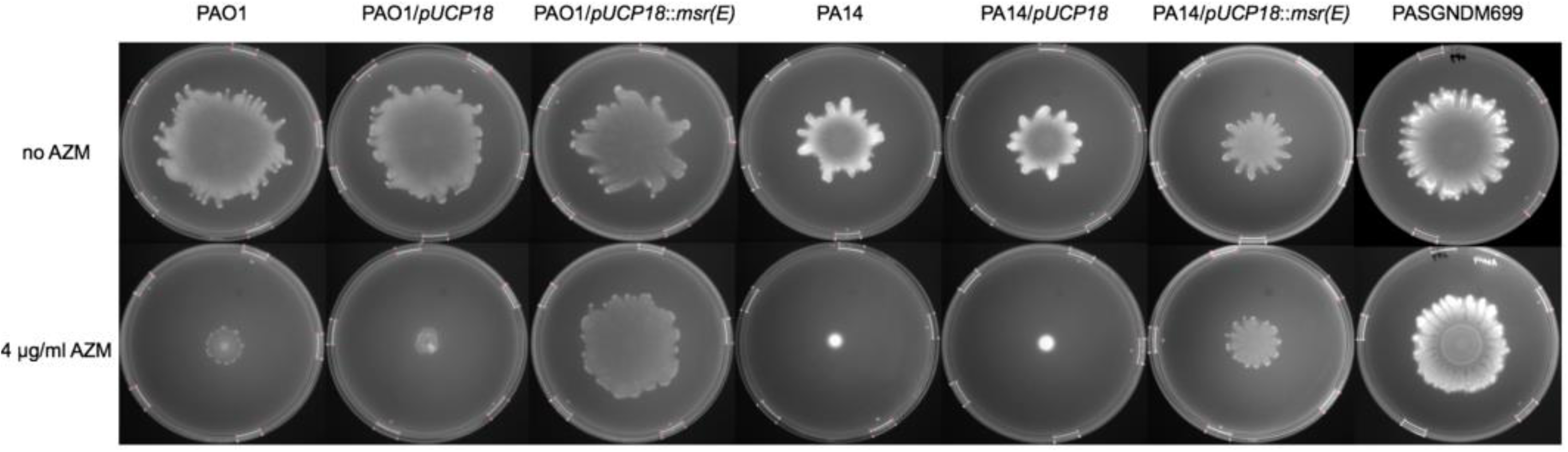
Impact of exogenous expression of Msr(E) on the swarming motilities of PAO1 and PA14 under 8 μg/ml of AZM. The swarming motility of wildtype strains and vector-carrying strains of PAO1 and PA14 were inhibited by the addition of 4 μg/ml of AZM, whereas the exogenous expression of Msr(E) in both strains restored the AZM-mediated inhibition on swarming motilities. No swarming inhibition by AZM was observed in PASGNDM699 strain with *msr(E)* encoded in the genome.

### Expression of *msr(E)* in *P. aeruginosa* restores AZM-affected transcriptome

AZM was previously reported to affect the transcriptome of *P. aeruginosa* in a microarray analysis (32). To better illustrate the effect of Msr(E) on the AZM-affected transcriptome of *P. aeruginosa*, we compared the gene expression profiles of PAO1/*pUCP18*::*msr(E)* in the presence and absence of 8 μg/ml AZM by total RNA sequencing. In PAO1, a total of 550 genes (∼10% of PAO1 genome) were differentially expressed upon AZM treatment, of which 305 were upregulated and 245 were downregulated (Table S4, Fig. 6). We found that the upregulated genes are enriched in genes encoding, the type III secretion pathway apparatus, and other possible defense mechanisms against AZM (*infA* and *efp*) (Table S5). Whereas the downregulated genes include *rsmY* and *rsmZ*, the non-coding RNA products of which positively regulates QS by sequestering the translational inhibitor RsmA, *pslE-I* of the Psl synthesis operon, *lasI* of the LasI/LasR QS system, and the *flK* gene the expression of which is essential for swarming motility (Table S5). These results are consistent with findings in previous studies (21, 24, 31-33) and could explain the QS-inhibition effect of AZM on PAO1 as shown in this study (Fig. 4, Fig.5, and Fig.6). Surprisingly, the transcriptome of PAO1/*pUCP18*::*msr(E)* was almost not affected by AZM, for which only 6 genes were differentially expressed upon AZM treatment (Table S6, Fig. 6). Therefore, Msr(E) probably protected *P. aeruginosa* from AZM-mediated QS inhibition by restoring AZM-affected transcriptome.

**Figure 6:**
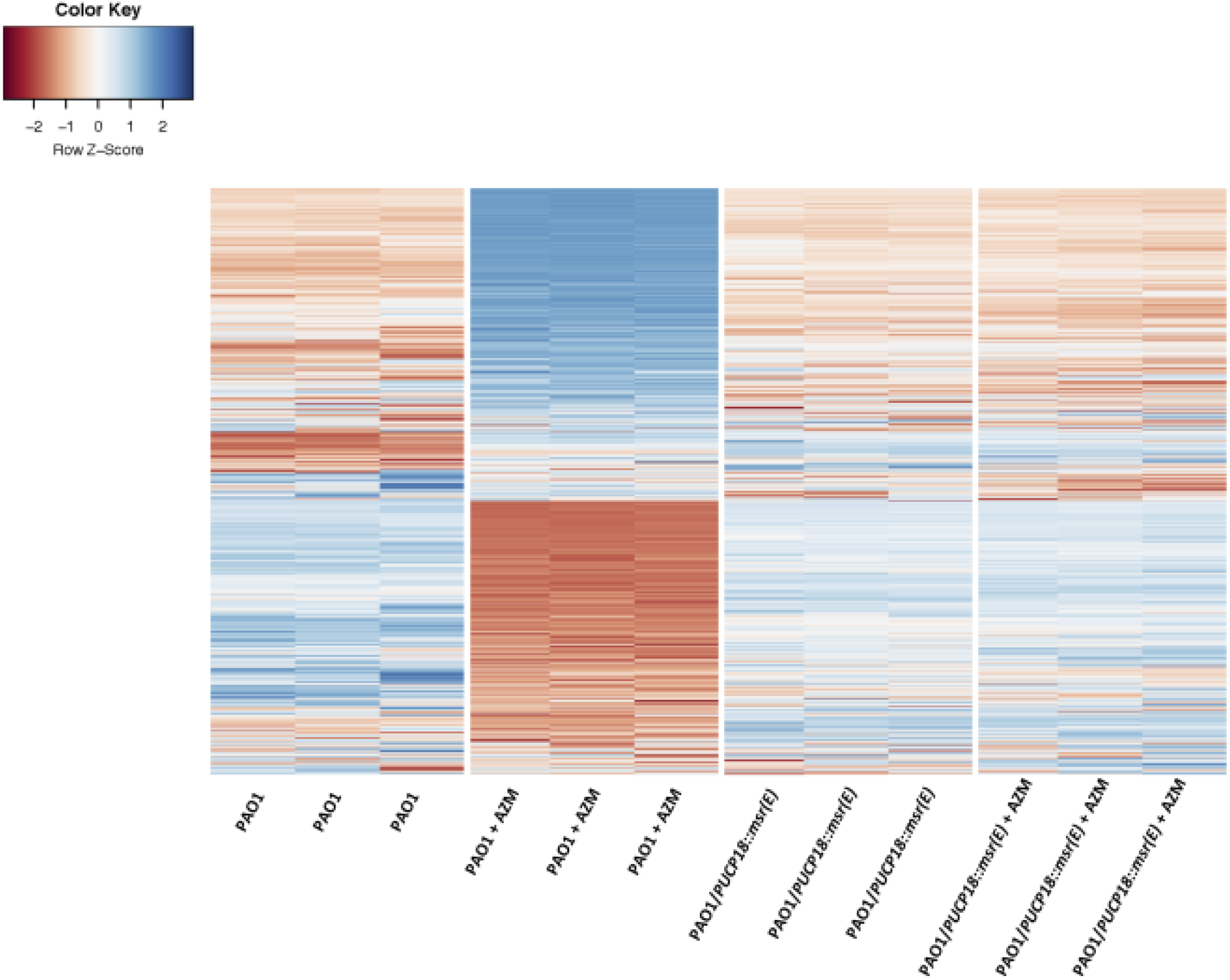
Heat map of 550 genes with significant changes between the AZM-treated and non-treated PAO1, including their behaviors in the AZM-treated and non-treated PAO1/*pUCP18::msr(E)*. The differentially expressed genes (fold-change > 4, P value < 0.05) between the AZM-treated and non-treated PAO1 cells were identified by performing a negative binomial test using the DESeq 2 package of R/Bioconductor. Whereas similar change in the gene expression profile was not observed in PAO1/*pUCP18::msr(E)* cells when treated and not treated with AZM. The full lists of genes that were differentially expressed between AZM-treated and non-treated PAO1 and between AZM-treated and non-treated PAO1/*pUCP18::msr(E)* can be found in Table S4 and Table S5.

### Msr(E) abolished anti-Pseudomonas effect of AZM *in vivo*

Previous studies showed that macrolide antibiotics such as erythromycin and clarithromycin could enhance the clearance of *P. aeruginosa* at the infection sites in a murine implant infection model and a murine lung infection model (34, 35). To investigate if the acquisition of *msr(E)* by *P. aeruginosa* can affect the anti-Pseudomonas activity of AZM *in vivo*, we used the murine implant model to compare the effectiveness of AZM treatment on PAO1 and PAO1/*pUCP18*::*msr(E)* infections. Briefly, silicone implants pre-colonized with PAO1 or PAO1/*pUCP18*::*msr(E)* strains were inserted into the peritoneal cavity of mice. After 12 hours of incubation, mice were treated with AZM by injecting AZM solution into the peritoneal cavity, whereas the control mice were injected with the same amount of saline. The silicone implants and mice spleens were harvested at 24-hour post infection to enumerate colony forming units (CFU) of *P. aeruginosa*, which were used to indicate the clearance of bacterial at infection site and the spreading of infection to other organs according to previous studies (35, 36). We found that AZM treatment could reduce the CFU of PAO1 residing in the silicone implants and the mice spleens by 2.0-log and 3.8-log, respectively (Fig. 7). By contrast, AZM treatment did not significantly affect the CFU of PAO1/*pUCP18*::*msr(E)* recovered from either the silicone implants or the mice spleens, as compared to the saline-treated control group (Fig. 7). These results indicated that Msr(E) could confer *P. aeruginosa* resistance against the anti-Pseudomonas activity of AZM *in vivo*.

**Figure 7:**
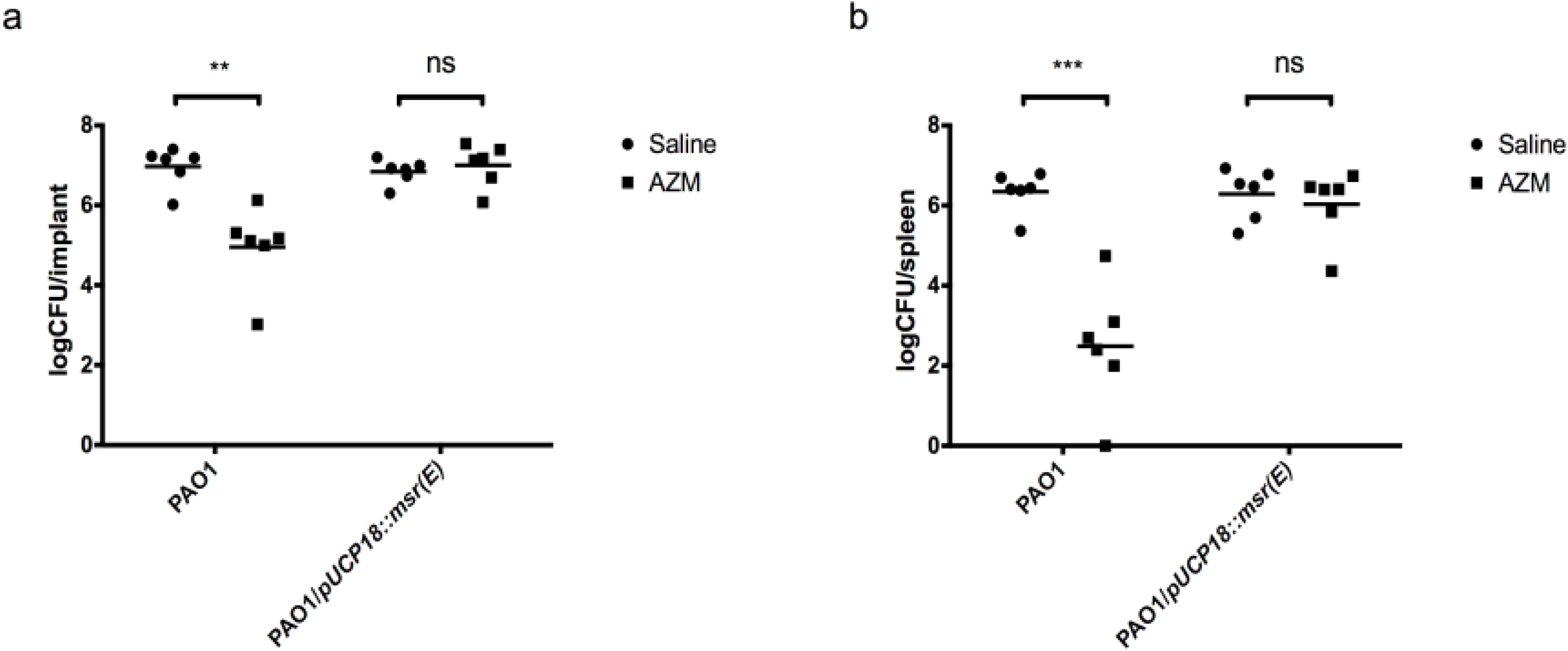
The effect of AZM treatment on PAO1 and PAO1/*pUCP18::msr(E)* in a murine silicone implant model. The silicone implants attached by PAO1 and PAO1/*pUCP18::msr(E)* were inserted into the peritoneal cavity of mice by surgery. AZM treatment at 10 mg per mouse significantly reduced the CFU of PAO1 recovered from (a) the silicone implants by 2.0-log and (b) the mice spleens by 3.8-log, whereas it had no effect on the CFU of PAO1/*pUCP18::msr(E)* recovered from either the silicone implants (a) or the mice spleens (b). Results are shown as log CFU per silicone implant and log CFU per mouse spleen, respectively (n = 6, ns P > 0.05, ** P < 0.01, **** P < 0.0001, Student’s t-test)

## Discussion

Tn4371 family ICEs have been previously described in a broad range of β-and γ-proteobacteria isolated from both environmental and clinical settings, which could confer their hosts adaptive functions such as multidrug/heavy metal resistance and the ability to metabolize specific carbon sources (25). Furthermore, a Tn4371-like element from *P. aeruginosa* genomes has recently been characterized to carry *bla*_*SPM-1*_ and *bcr1*, suggesting that the Tn4371 family ICEs is an important source for the acquisition of carbapenem resistance by *P. aeruginosa* (37). In the current work, we identified a novel ICE_Tn4371_6385 from PASGNDM699 isolated from clinical sources. This element carries three antibiotic resistance genes including *bla*_*NDM-1*_, *msr(E)* and *floR*, and retains the intact machinery for conjugative transfer (Fig. 3), suggesting that it is responsible for the transmission of these resistance genes into the PASGNDM genomes (25, 38). However, the conjugative transfer of ICE_Tn4371_6385 was not observed in mating experiments between PASGNDM699 and several recipient *P. aeruginosa* strains under laboratory conditions (data not shown). It is possible that the conjugation has extremely low efficiencies that is below our detection limit or requires specific inductive conditions (38, 39). Our findings provide the first evidence that *bla*_*NDM-1*_ is encoded and potentially disseminated by ICEs, together with *msr(E)* and *floR*. The macrolide resistance gene *msr(E)* and its variants were mainly identified from Gram-positive bacteria and several Gram-negative bacterial species such as Enterobacteriaceae spp., *Pasteurella multocida* and *A. baumannii*, but has been rarely described in *P. aeruginosa* (29, 40-42). Hence, the transmission of *msr(E)* by ICE_Tn4371_6385 into the clinical isolates characterized in this study has led us to investigate its possible functional roles in *P. aeruginosa*. Our *in vitro* assays clearly showed that the exogenous expression of *msr(E)* in *P. aeruginosa* could abolish the QS inhibition activity of AZM *in vitro*, which in turn rescued the inhibition of elastase production, rhamnolipid production and swarming motility by AZM treatment (Fig. 4, 5). In addition, the acquisition of *msr(E)* by PAO1 almost completely restored the AZM-affected transcriptome and abolished the anti-Pseudomonas activity of AZM in a murine infection model (Fig. 6, Fig. 7, Table S5, Table S6). These results demonstrate that Msr(E) could confer *P. aeruginosa* with resistance to AZM-mediated QS inhibition and that the transmission of *msr(E)* into this organism will greatly challenge the use of AZM in treating infections caused by *P. aeruginosa*.

The third antibiotic resistance gene carried by ICE_Tn4371_6385 is *floR*, which encodes an efflux pump and is the determinant for florfenicol resistance (43). This gene was first identified in *Salmonella typhimurium* DT104 in 1999, and has ever since been detected in *E. coli and P. multocida* isolated from livestock and aquaculture settings (44-47). It is possible that the emergence and spreading of *floR* is due to the selection by florfenicol, since this antibiotic has been extensively used in veterinary medicine, especially in aquaculture settings (46, 48). Therefore, the carriage of *floR* by ICE_Tn4371_6385 may confer its hosts with advantages under environmental conditions where florfenicol may be present (48, 49), and hence facilitate the transmission and dissemination of *bla*_*NDM-1*_ and *msr(E)*.

In conclusion, we have characterized the full-length genomes of two NDM-1-producing *P. aeruginosa* clinical isolates, from which we identified a novel ICE_Tn4371_6385 element encoding three antibiotic resistance genes. Among them, the *bla*_*NDM-1*_ gene is responsible for extreme resistance to anti-Pseudomonal carbapenems, whereas *msr(E)* can quench AZM-mediated QS inhibition, and *floR* may enhance the survival of host bacteria under florfenicol exposure. We present the first evidence of *bla*_*NDM-1*_ to be carried by ICEs, and the first evidence of co-transfer of carbapenem resistance and the resistance to macrolide-mediated QS inhibition into *P. aeruginosa*. Our findings highlight the importance of ICEs in transmitting antibiotic resistance, and that anti-virulence treatment of *P. aeruginosa* infections by targeting QS can be challenged by horizontal transfer of resistance mechanisms.

## Materials and Methods

### Bacterial strains and plasmids

All the strains were routinely grown in Lysogeny broth (LB) or on LB plates with 1.5% agar at 37°C. The *msr(E)* gene with its putative promoter were amplified from PASGNDM699 genome using primers 5’ACCGGCCAAGATAGTTGACG3’ and 5’AGGAAGTTCAACCGCCCTTT3’ and ligated into the smaI site of *pUCP18* vector (50) to construct *pUCP18::msr(E)* plasmid, where the *msr(E)* gene is upstream to the *lac* promoter. Both *pUCP18* and *pUCP18::msr(E)* were transformed into PAO1 and PA14 by electroporation, and the strains carrying these two plasmids were grown in LB supplemented with 300 μg/ml of carbenicillin (MP Bio) to maintain the plasmids.

### Sequencing, assembly, and annotation

Genomic DNA was purified using Blood and Cell Culture DNA Midi Kit (Qiagen) and sequenced on an Illumina MiSeq platform or a PacBio RS II system. The full-length genomes of PASGNDM345 and PASGNDM699 were assembled from long reads obtained from the PacBio RS II system using HGAP2 pipeline assisted with manual curation. The ICE_Tn4371_6385 sequence and ICE-like element from *P. fluorescens* UK4 were uploaded to the Rapid Annotations using Subsystem Technology (RAST) server for gene prediction and annotation, assisted with manual curation (51). Comparison of the two elements was done using EasyFig v2.2.2 (52).

### Phylogenetic and comparative genomic analysis

Phylogenetic analysis was carried out using Parsnp v1.1.2 (53). The phylogenetic tree was constructed using the approximate maximum likelihood algorithm based on core genome alignment, with filtration of recombination sites. Genomic islands of PASGNDM699 and PASGNDM345 genomes were predicted by the IslandViewer 3 server (54), and the encoded antibiotic resistance genes were predicted using the ResFinder 2.1 server (55). Comparison of PASGNDM699 and PASGNDM345 genomes with six other full-length *P. aeruginosa* genomes was done by BLASTn search using BLAST Ring Image Generator v0.95 (56), with genomic islands and antibiotic resistance genes plotted in the same image.

### Quantification of elastase, rhamnolipid and swarming motility

Bacterial strains were grown in ABT minimal medium supplemented with 5 g l_–1_ glucose and 2 g l_–1_ casamino acids (ABTGC), with or without addition of 8 μg/ml AZM (Spectrum). The supernatants of overnight culture were filter sterilised before quantification of elastase and rhamnolipid production. Elastase activity was measured using EnzCheck Elastase Assay Kit (Thermo Fisher Scientific) as instructed by the manual. The quantification of rhamnolipid was performed as previously described with modifications (57). Briefly, rhamnolipid was extracted from supernatants with diethyl esther and dissolved in deionised water. The solution was added with 0.19% (w/v) orcinol dissolved in 50% H_2_SO_4_, followed by heating at 80°C for 30 min. The elastase activity and rhamnolipid were quantified by measuring emission at 530 nm and absorbance at 460 nm by using an Infinite 200 PRO system (Tecan), respectively. Both experiments were performed in at least triplicates, and the results were normalised with optical density at 600 nm (OD_600_) and shown as mean ± standard deviation in the figures. The swarming motility of each strain was measured on 0.5% agar plates containing 8 g/L nutrient broth (Oxoid) and 5 g/L glucose (58). One μl of bacterial overnight culture (adjusted to OD_600_ = 1) was inoculated to the center of the plate, followed by incubation at 37°C for 12 hours. The images were taken by a Gel Doc XR+ System (Bio-Rad).

### Transcriptomic analysis

Strains were grown in ABTGC medium at 37°C with 200 rpm shaking. Cells were harvested at mid-log phase (OD_600_ = 0.6), and the total RNA was extracted using RNeasy Mini Kit (Qiagen). RNA sequencing was performed on an Illumina HiSeq 2000 platform to generate paired-end reads of 100 nt, which were mapped to PAO1 genome (NC_002516.2) by using CLC Genomics Workbench 9.0 (Qiagen). The transcript count table was analysed by DESeq 2 package of the R/Bioconductor by performing negative binomial test (59). Hierarchical clustering analysis was performed to produce the heatmap for the differentially expressed genes with statistical significance (fold change > 4, P <0.05) using heatmap.2 package of the R/Bioconductor (60).

### Animal model

A murine silicone implant model was used to evaluate effects of AZM treatment *in vivo* on the two strains as previously described with modifications (36). Briefly, washed bacterial cells were re-suspended in 0.9% NaCl to an OD_600_ of 0.01. Bacteria cells were allowed to attach onto sterilize silicone tubes (length, 3 mm; inner diameter, 4 mm; outer diameter, 6 mm; Ole Dich) by incubation at 37°C, with 110 rpm shaking for 18 hours, after which the silicone tubes coated with bacteria were implanted into mouse peritoneal cavity by surgery. Treatment with AZM (10 mg per mouse, dissolved in 0.2 ml of 0.9% saline) or saline was performed by injection into the mouse peritoneal cavity 12 hours after surgery. Mice were sacraficed 24 after surgery and the bacterial cells residing on the silicone tube and in the mouse spleen were dispersed into the 0.9% NaCl solution by sonication using an Elmasonic P120H (Elma, Germany; power=50% and frequency=37 KHz) and homogenisation using a Bio-Gen PRO200 Homogenizer (Pro Scientific), respectively. The CFU was quantified by serial dilution and plating on LB agar plates, and the results were shown in log CFU with mean ± standard deviation.

### Ethics

The use of clinical specimen samples was approved by Department of Laboratory Medicine, National University Hospital, Singapore, registered under the reference 2016/00856. The animal model protocols were approved by the Institutional Animal Care and Use Committee (IACUC) of Nanyang Technological University, under the permit number A-0191 AZ. Animal experiments were performed in accordance to the NACLAR Guidelines of Animal and Birds (Care and Use of Animals for Scientific Purposes) Rules by Agri-Food & Authority of Singapore (AVA).

### Accession numbers

The genome sequencing data and RNA-Seq data have been deposited in the NCBI Short Read Archive database with the accession codes SRP103165 and SRP103155. The complete and draft genome sequences have been submitted to NCBI registered under BioProject PRJNA381838.

## Acknowledgments

This research was supported by the National Research Foundation and Ministry of Education Singapore under its Research Centre of Excellence Program and AcRF Tier 2 (MOE2014-T2-2-172) from Ministry of Education, Singapore. We thank Zi Jing Seng for her help in RNA extraction for *P. aeruginosa* RNA-seq analysis.

Y.D. and L.Y. designed the experiments; J.T. collected clinical samples and conducted sample identification; D.I.D. did RNA sequencing; Y.D. analyzed sequencing data, carried out laboratory experiments, animal work, interpreted results and wrote the manuscript; All authors read and commented on the manuscript.

---

**Figure S1:**
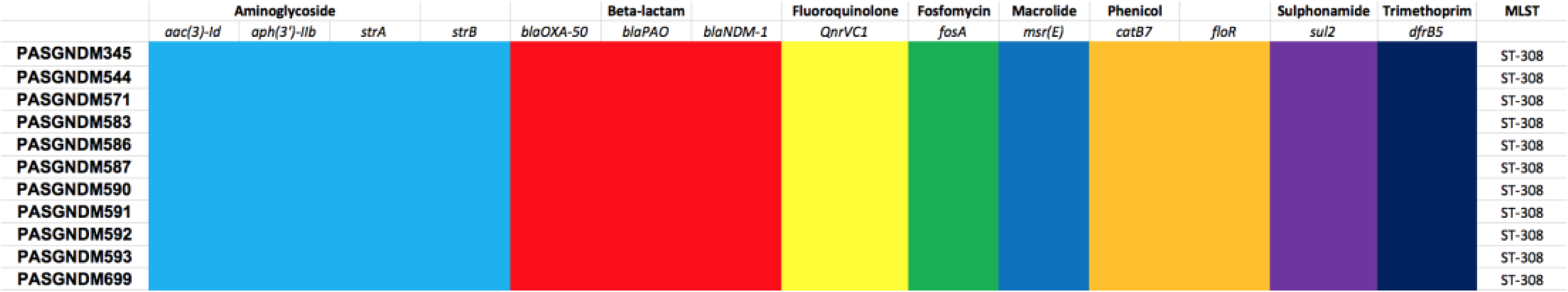
Antibiotic resistance genes present in 11 PASGNDM genomes. Presence of antibiotic resistance genes in genomes were predict by ResFinder 2.1 server (55). Multilocus sequencing type (MLST) of each strain was determined using DTU MLST server 1.8 (61).

**Figure S2:**
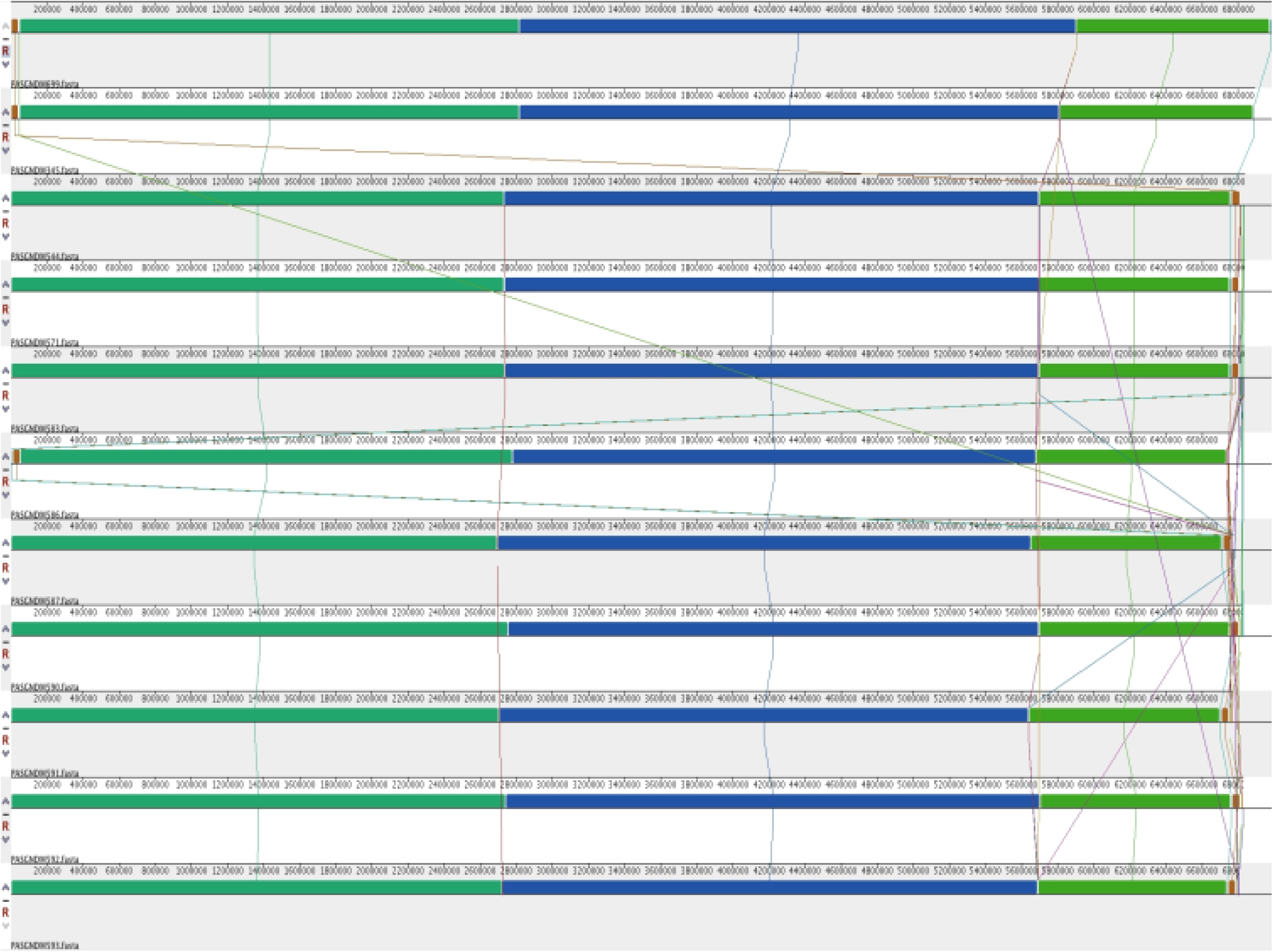
Alignment of the 11 PASGNDM genomes. Complete genomes of PASGNDM345 and PASGNDM699 were aligned with draft genomes of the other 9 PASGNDM strains using Progressive Mauve (26). Alignment results showed the 11 strains are highly similar.

**Table S1:** The MICs of PASGNDM strains to various antibiotics. The strains only remain sensitive to polymyxin B. Clinical specimens from which the strains were isolated are also shown in the table.

| Strain | Specimen | Ceftazidime | Cefepime | Imipenem | Meropenem | Gentamicin | Amikacin | Ciprofloxacin | Levofloxacin | Ertapenem | Polymyxin B |
| --- | --- | --- | --- | --- | --- | --- | --- | --- | --- | --- | --- |
| PASGNDM544 | ETT aspirate | >64 | >64 | >16 | >16 | >16 | >64 | >4 | >8 | >8 | 1.5 |
| PASGNDM699 | Sputum | >64 | >64 | >16 | >16 | >16 | >64 | >4 | >8 | >8 | 1.5 |
| PASGNDM345 | Sputum | >64 | >64 | >16 | >16 | >16 | >64 | >4 | >8 | >8 | 2 |
| PASGNDM571 | Urine | >64 | >64 | >16 | >16 | >16 | >64 | >4 | >8 | >8 | 1 |
| PASGNDM583 | Urine | >64 | >64 | >16 | >16 | >16 | >64 | >4 | >8 | >8 | 1 |
| PASGNDM586 | Urine | >64 | >64 | >16 | >16 | >16 | >64 | >4 | >8 | >8 | 0.5 |
| PASGNDM587 | Wound swab | >64 | >64 | >16 | >16 | >16 | >64 | >4 | >8 | >8 | 0.5 |
| PASGNDM590 | Urine | >64 | >64 | >16 | >16 | >16 | >64 | >4 | >8 | >8 | 0.75 |
| PASGNDM591 | Urine | >64 | >64 | >16 | >16 | >16 | >64 | >4 | >8 | >8 | 2 |
| PASGNDM592 | Urine | >64 | >64 | >16 | >16 | >16 | >64 | >4 | >8 | >8 | 2 |
| PASGNDM593 | Urine | >64 | >64 | >16 | >16 | >16 | >64 | >4 | >8 | >8 | 0.75 |

**Table S2:** Genome features of PASGNDM699 and PASGNDM345.

|  | **Length**<br>**(bp)** | **G+C**<br>**content** | **Genes** | **Protein-coding**<br>**genes** | **tRNA** | **rRNA** |
| --- | --- | --- | --- | --- | --- | --- |
| PASGNDM345 | 6,893,164 | 66.1% | 6503 | 6427 | 64 | 12 |
| PASGNDM699 | 6,985,102 | 66.0% | 6589 | 6513 | 64 | 12 |

**Table S3:** Predicted genomic islands from the full-length genome of PASGNDM699. Fonts in bold indicates the GI only present in PASGNDM699 but not PASGNDM345 genome.

| **Island start** | **Island end** | **Length** | **Method** | **Gene start** | **Gene end** | **Product** |
| --- | --- | --- | --- | --- | --- | --- |
| 692910 | 699692 | 6782 | Predicted by at least one method | 692780 | 693475 | Phage minor tail protein |
| 692910 | 699692 | 6782 | Predicted by at least one method | 693478 | 694248 | Phage tail assembly protein |
| 692910 | 699692 | 6782 | Predicted by at least one method | 694303 | 694905 | Phage tail assembly protein I |
| 692910 | 699692 | 6782 | Predicted by at least one method | 694964 | 698617 | Phage tail fiber protein |
| 692910 | 699692 | 6782 | Predicted by at least one method | 699603 | 700703 | FIG00966085: hypothetical protein |
| 1060767 | 1077432 | 16665 | Predicted by at least one method | 1060868 | 1061077 | FIG00954142: hypothetical protein |
| 1060767 | 1077432 | 16665 | Predicted by at least one method | 1061279 | 1061758 | Transcriptional regulator, AsnC family |
| 1060767 | 1077432 | 16665 | Predicted by at least one method | 1062236 | 1062919 | FIG00954854: hypothetical protein |
| 1060767 | 1077432 | 16665 | Predicted by at least one method | 1062912 | 1063808 | Cobalt-zinc-cadmium resistance protein |
| 1060767 | 1077432 | 16665 | Predicted by at least one method | 1063897 | 1064313 | Polyribonucleotide nucleotidyltransferase (polynucleotide phosphorylase) |
| 1060767 | 1077432 | 16665 | Predicted by at least one method | 1064455 | 1066968 | ATP-dependent helicase HrpB |
| 1060767 | 1077432 | 16665 | Predicted by at least one method | 1067024 | 1068490 | site-specific recombinase, phage integrase family |
| 1060767 | 1077432 | 16665 | Predicted by at least one method | 1068487 | 1070415 | hypothetical protein |
| 1060767 | 1077432 | 16665 | Predicted by at least one method | 1070412 | 1072460 | Phage Integrase |
| 1060767 | 1077432 | 16665 | Predicted by at least one method | 1073024 | 1073206 | Transcriptional regulator, XRE family |
| 1060767 | 1077432 | 16665 | Predicted by at least one method | 1073925 | 1075325 | FOG: GGDEF domain |
| 1060767 | 1077432 | 16665 | Predicted by at least one method | 1076625 | 1076921 | Mobile element protein |
| 1060767 | 1077432 | 16665 | Predicted by at least one method | 1076944 | 1077324 | Mobile element protein |
| 1250067 | 1256766 | 6699 | Predicted by at least one method | 1250412 | 1251239 | Integrase |
| 1250067 | 1256766 | 6699 | Predicted by at least one method | 1251864 | 1252256 | Rossmann fold nucleotide-binding protein Smf possibly involved in DNA uptake |
| 1250067 | 1256766 | 6699 | Predicted by at least one method | 1252789 | 1252905 | hypothetical protein |
| 1250067 | 1256766 | 6699 | Predicted by at least one method | 1252954 | 1253256 | hypothetical protein |
| 1250067 | 1256766 | 6699 | Predicted by at least one method | 1253253 | 1253594 | hypothetical protein |
| 1250067 | 1256766 | 6699 | Predicted by at least one method | 1253599 | 1254288 | Phage protein |
| 1250067 | 1256766 | 6699 | Predicted by at least one method | 1254461 | 1254652 | Phage protein |
| 1250067 | 1256766 | 6699 | Predicted by at least one method | 1254652 | 1254768 | Phage protein |
| 1250067 | 1256766 | 6699 | Predicted by at least one method | 1254765 | 1255427 | Phage protein |
| 1250067 | 1256766 | 6699 | Predicted by at least one method | 1255424 | 1255759 | Phage protein |
| 1250067 | 1256766 | 6699 | Predicted by at least one method | 1255762 | 1255992 | Phage protein |
| 1250067 | 1256766 | 6699 | Predicted by at least one method | 1256090 | 1256323 | hypothetical protein |
| 1250067 | 1256766 | 6699 | Predicted by at least one method | 1256326 | 1256712 | Transcriptional regulator, LuxR family |
| 1259039 | 1294047 | 35008 | Predicted by at least one method | 1260441 | 1260728 | hypothetical protein |
| 1259039 | 1294047 | 35008 | Predicted by at least one method | 1260725 | 1261033 | hypothetical protein |
| 1259039 | 1294047 | 35008 | Predicted by at least one method | 1261301 | 1261534 | hypothetical protein |
| 1259039 | 1294047 | 35008 | Predicted by at least one method | 1261780 | 1262301 | hypothetical protein |
| 1259039 | 1294047 | 35008 | Predicted by at least one method | 1262298 | 1262528 | FIG00964051: hypothetical protein |
| 1259039 | 1294047 | 35008 | Predicted by at least one method | 1262697 | 1263698 | Primosomal protein I |
| 1259039 | 1294047 | 35008 | Predicted by at least one method | 1263695 | 1264483 | DNA replication protein DnaC |
| 1259039 | 1294047 | 35008 | Predicted by at least one method | 1264480 | 1265910 | Replicative DNA helicase |
| 1259039 | 1294047 | 35008 | Predicted by at least one method | 1265943 | 1266191 | FIG00956875: hypothetical protein |
| 1259039 | 1294047 | 35008 | Predicted by at least one method | 1268528 | 1268860 | phage holin, lambda family |
| 1259039 | 1294047 | 35008 | Predicted by at least one method | 1268857 | 1269474 | Lytic enzyme |
| 1259039 | 1294047 | 35008 | Predicted by at least one method | 1269549 | 1269713 | hypothetical protein |
| 1259039 | 1294047 | 35008 | Predicted by at least one method | 1269665 | 1269907 | hypothetical protein |
| 1259039 | 1294047 | 35008 | Predicted by at least one method | 1271731 | 1273506 | Phage terminase, large subunit |
| 1259039 | 1294047 | 35008 | Predicted by at least one method | 1273506 | 1273709 | hypothetical protein |
| 1259039 | 1294047 | 35008 | Predicted by at least one method | 1273711 | 1275237 | Phage portal protein |
| 1259039 | 1294047 | 35008 | Predicted by at least one method | 1275245 | 1277200 | ATP-dependent Clp protease proteolytic subunit |
| 1259039 | 1294047 | 35008 | Predicted by at least one method | 1277272 | 1277634 | hypothetical protein |
| 1259039 | 1294047 | 35008 | Predicted by at least one method | 1277953 | 1278426 | Phage protein |
| 1259039 | 1294047 | 35008 | Predicted by at least one method | 1278430 | 1278624 | FIG00964642: hypothetical protein |
| 1259039 | 1294047 | 35008 | Predicted by at least one method | 1278626 | 1279378 | hypothetical protein |
| 1259039 | 1294047 | 35008 | Predicted by at least one method | 1280513 | 1282996 | FIG00954405: hypothetical protein |
| 1259039 | 1294047 | 35008 | Predicted by at least one method | 1283420 | 1283557 | PE-PGRS virulence associated protein |
| 1259039 | 1294047 | 35008 | Predicted by at least one method | 1283557 | 1285254 | FIG00956273: hypothetical protein |
| 1259039 | 1294047 | 35008 | Predicted by at least one method | 1285257 | 1285667 | FIG00964183: hypothetical protein |
| 1259039 | 1294047 | 35008 | Predicted by at least one method | 1286223 | 1286627 | FIG00963969: hypothetical protein |
| 1259039 | 1294047 | 35008 | Predicted by at least one method | 1286624 | 1288282 | COG1156: Archaeal/vacuolar-type H<sup>+</sup>-ATPase subunit B |
| 1259039 | 1294047 | 35008 | Predicted by at least one method | 1288312 | 1288899 | FIG00967075: hypothetical protein |
| 1259039 | 1294047 | 35008 | Predicted by at least one method | 1289055 | 1289648 | FIG00965686: hypothetical protein |
| 1259039 | 1294047 | 35008 | Predicted by at least one method | 1289654 | 1289929 | FIG00956286: hypothetical protein |
| 1259039 | 1294047 | 35008 | Predicted by at least one method | 1290524 | 1290826 | FIG00958463: hypothetical protein |
| 1259039 | 1294047 | 35008 | Predicted by at least one method | 1290823 | 1291053 | Phage protein |
| 1259039 | 1294047 | 35008 | Predicted by at least one method | 1291219 | 1291356 | Phage protein |
| 1259039 | 1294047 | 35008 | Predicted by at least one method | 1291714 | 1291863 | hypothetical protein |
| 1259039 | 1294047 | 35008 | Predicted by at least one method | 1293131 | 1293331 | hypothetical protein |
| 1351730 | 1356817 | 5087 | Predicted by at least one method | 1351730 | 1351993 | Transcriptional regulator in PFGI-1-like cluster |
| 1351730 | 1356817 | 5087 | Predicted by at least one method | 1352073 | 1352834 | FIG00642059: hypothetical protein |
| 1351730 | 1356817 | 5087 | Predicted by at least one method | 1353145 | 1353378 | FIG01214592: hypothetical protein |
| 1351730 | 1356817 | 5087 | Predicted by at least one method | 1353712 | 1354257 | RecA-family ATPase |
| 1351730 | 1356817 | 5087 | Predicted by at least one method | 1354304 | 1355116 | Phage replication protein |
| 1351730 | 1356817 | 5087 | Predicted by at least one method | 1355848 | 1356126 | IncP-type oriT binding protein TraK |
| 1351730 | 1356817 | 5087 | Predicted by at least one method | 1356446 | 1356817 | IncP-type oriT binding protein TraJ |
| 1356446 | 1376081 | 19635 | Predicted by at least one method | 1356446 | 1356817 | IncP-type oriT binding protein TraJ |
| 1356446 | 1376081 | 19635 | Predicted by at least one method | 1357083 | 1357868 | Conjugative transfer protein TrbJ |
| 1356446 | 1376081 | 19635 | Predicted by at least one method | 1357884 | 1358153 | Conjugative transfer entry exclusion protein TrbK |
| 1356446 | 1376081 | 19635 | Predicted by at least one method | 1358150 | 1359739 | Conjugative transfer protein TrbL |
| 1356446 | 1376081 | 19635 | Predicted by at least one method | 1360077 | 1360307 | FIG01213713: hypothetical protein |
| 1356446 | 1376081 | 19635 | Predicted by at least one method | 1360552 | 1361100 | hypothetical protein |
| 1356446 | 1376081 | 19635 | Predicted by at least one method | 1361271 | 1361489 | Mercuric transport protein, MerE |
| 1356446 | 1376081 | 19635 | Predicted by at least one method | 1361486 | 1361851 | Mercuric resistance operon coregulator |
| 1356446 | 1376081 | 19635 | Predicted by at least one method | 1362616 | 1364304 | Mercuric ion reductase |
| 1356446 | 1376081 | 19635 | Predicted by at least one method | 1364315 | 1364593 | Periplasmic mercury(+2) binding protein |
| 1356446 | 1376081 | 19635 | Predicted by at least one method | 1364607 | 1364957 | Mercuric transport protein, MerT |
| 1356446 | 1376081 | 19635 | Predicted by at least one method | 1365029 | 1365187 | Mercuric resistance operon regulatory protein |
| 1356446 | 1376081 | 19635 | Predicted by at least one method | 1365257 | 1366093 | Aminoglycoside 3'-phosphotransferase 2 @ Streptomycin 3'-kinase StrB |
| 1356446 | 1376081 | 19635 | Predicted by at least one method | 1367702 | 1370587 | Mobile element protein |
| 1356446 | 1376081 | 19635 | Predicted by at least one method | 1370666 | 1371232 | Mobile element protein |
| 1356446 | 1376081 | 19635 | Predicted by at least one method | 1371512 | 1372147 | Type IV secretory pathway, VirD2 components (relaxase) |
| 1356446 | 1376081 | 19635 | Predicted by at least one method | 1374023 | 1375516 | Mobile element protein |
| 1370666 | 1375516 | 4850 | Predicted by at least one method | 1370666 | 1371232 | Mobile element protein |
| 1370666 | 1375516 | 4850 | Predicted by at least one method | 1371512 | 1372147 | Type IV secretory pathway, VirD2 components (relaxase) |
| 1370666 | 1375516 | 4850 | Predicted by at least one method | 1374023 | 1375516 | Mobile element protein |
| 1824170 | 1828537 | 4367 | Predicted by at least one method | 1824170 | 1824571 | hypothetical protein |
| 1824170 | 1828537 | 4367 | Predicted by at least one method | 1824761 | 1825951 | NADH oxidase |
| 1824170 | 1828537 | 4367 | Predicted by at least one method | 1825932 | 1826537 | hypothetical protein |
| 1824170 | 1828537 | 4367 | Predicted by at least one method | 1827041 | 1828537 | Type III restriction-modification system methylation subunit |
| 1996654 | 2001263 | 4609 | Predicted by at least one method | 1996654 | 1997271 | FIG01219979: hypothetical protein |
| 1996654 | 2001263 | 4609 | Predicted by at least one method | 1997481 | 1997618 | Glycine cleavage system H protein |
| 1996654 | 2001263 | 4609 | Predicted by at least one method | 1997978 | 1998154 | hypothetical protein |
| 1996654 | 2001263 | 4609 | Predicted by at least one method | 1998205 | 1998816 | FIG00984683: hypothetical protein |
| 1996654 | 2001263 | 4609 | Predicted by at least one method | 1998961 | 1999578 | Glutathione S-transferase |
| 1996654 | 2001263 | 4609 | Predicted by at least one method | 1999654 | 2000208 | Multimeric flavodoxin WrbA |
| 1996654 | 2001263 | 4609 | Predicted by at least one method | 2000310 | 2001263 | putative LysR-family transcriptional regulator |
| 2057249 | 2078394 | 21145 | Predicted by at least one method | 2057249 | 2058346 | Chorismate mutase I / Prephenate dehydratase |
| 2057249 | 2078394 | 21145 | Predicted by at least one method | 2058415 | 2059524 | Biosynthetic Aromatic amino acid aminotransferase beta |
| 2057249 | 2078394 | 21145 | Predicted by at least one method | 2059517 | 2061757 | Cyclohexadienyl dehydrogenase(EC 1.3.1.43) / 5-Enolpyruvylshikimate-3-phosphate synthase |
| 2057249 | 2078394 | 21145 | Predicted by at least one method | 2061757 | 2062446 | Cytidylate kinase |
| 2057249 | 2078394 | 21145 | Predicted by at least one method | 2062714 | 2064393 | SSU ribosomal protein S1p |
| 2057249 | 2078394 | 21145 | Predicted by at least one method | 2064530 | 2064814 | Integration host factor beta subunit |
| 2057249 | 2078394 | 21145 | Predicted by at least one method | 2070388 | 2070660 | UDP-N-acetylglucosamine 4,6-dehydratase |
| 2057249 | 2078394 | 21145 | Predicted by at least one method | 2070675 | 2070827 | UDP-N-acetylglucosamine 4,6-dehydratase |
| 2057249 | 2078394 | 21145 | Predicted by at least one method | 2070831 | 2071949 | WbjC |
| 2057249 | 2078394 | 21145 | Predicted by at least one method | 2072270 | 2073097 | UDP-N-acetylglucosamine 2-epimerase |
| 2057249 | 2078394 | 21145 | Predicted by at least one method | 2073409 | 2074317 | Putative glycosyltransferase |
| 2057249 | 2078394 | 21145 | Predicted by at least one method | 2074575 | 2075258 | UDP-glucose 4-epimerase |
| 2057249 | 2078394 | 21145 | Predicted by at least one method | 2076532 | 2078394 | nucleotide sugar epimerase/dehydratase WbpM |
| 2457987 | 2469377 | 11390 | Predicted by at least one method | 2457819 | 2458094 | hypothetical protein |
| 2457987 | 2469377 | 11390 | Predicted by at least one method | 2458094 | 2458213 | hypothetical protein |
| 2457987 | 2469377 | 11390 | Predicted by at least one method | 2458198 | 2458410 | hypothetical protein |
| 2457987 | 2469377 | 11390 | Predicted by at least one method | 2458407 | 2460287 | C-5 cytosine-specific DNA methylase family protein |
| 2457987 | 2469377 | 11390 | Predicted by at least one method | 2460284 | 2460466 | hypothetical protein |
| 2457987 | 2469377 | 11390 | Predicted by at least one method | 2460773 | 2461012 | hypothetical protein |
| 2457987 | 2469377 | 11390 | Predicted by at least one method | 2461009 | 2461635 | hypothetical protein |
| 2457987 | 2469377 | 11390 | Predicted by at least one method | 2461685 | 2462002 | transcriptional regulator, putative |
| 2457987 | 2469377 | 11390 | Predicted by at least one method | 2461999 | 2462229 | hypothetical protein |
| 2457987 | 2469377 | 11390 | Predicted by at least one method | 2462380 | 2462664 | hypothetical protein |
| 2457987 | 2469377 | 11390 | Predicted by at least one method | 2462683 | 2462994 | hypothetical protein |
| 2457987 | 2469377 | 11390 | Predicted by at least one method | 2464587 | 2464835 | hypothetical protein |
| 2457987 | 2469377 | 11390 | Predicted by at least one method | 2465325 | 2465609 | hypothetical protein |
| 2457987 | 2469377 | 11390 | Predicted by at least one method | 2465613 | 2465843 | hypothetical protein |
| 2457987 | 2469377 | 11390 | Predicted by at least one method | 2466057 | 2467178 | DNA primase, phage associated |
| 2533316 | 2539611 | 6295 | Predicted by at least one method | 2533619 | 2534038 | Hemoglobin-like protein HbO |
| 2533316 | 2539611 | 6295 | Predicted by at least one method | 2534625 | 2534738 | hypothetical protein |
| 2533316 | 2539611 | 6295 | Predicted by at least one method | 2539299 | 2541452 | hypothetical protein |
| 2541257 | 2548821 | 7564 | Predicted by at least one method | 2539299 | 2541452 | hypothetical protein |
| 2541257 | 2548821 | 7564 | Predicted by at least one method | 2541581 | 2544784 | hypothetical protein |
| 2541257 | 2548821 | 7564 | Predicted by at least one method | 2544797 | 2546125 | hypothetical protein |
| 2541257 | 2548821 | 7564 | Predicted by at least one method | 2546255 | 2546419 | hypothetical protein |
| 2541257 | 2548821 | 7564 | Predicted by at least one method | 2546504 | 2546710 | hypothetical protein |
| 2541257 | 2548821 | 7564 | Predicted by at least one method | 2547459 | 2547578 | hypothetical protein |
| 2541257 | 2548821 | 7564 | Predicted by at least one method | 2548819 | 2551380 | autotransporter, putative |
| 2549193 | 2554952 | 5759 | Predicted by at least one method | 2548819 | 2551380 | autotransporter, putative |
| 2549193 | 2554952 | 5759 | Predicted by at least one method | 2551578 | 2552024 | hypothetical protein |
| 2549193 | 2554952 | 5759 | Predicted by at least one method | 2552163 | 2552510 | Transcriptional regulator, XRE family |
| 2549193 | 2554952 | 5759 | Predicted by at least one method | 2552540 | 2552842 | hypothetical protein |
| 2549193 | 2554952 | 5759 | Predicted by at least one method | 2552961 | 2553143 | hypothetical protein |
| 2549193 | 2554952 | 5759 | Predicted by at least one method | 2553190 | 2554110 | FIG00965493: hypothetical protein |
| 2549193 | 2554952 | 5759 | Predicted by at least one method | 2554190 | 2555194 | Phage-related protein |
| 2557074 | 2571538 | 14464 | Predicted by at least one method | 2556939 | 2557268 | FIG00957676: hypothetical protein |
| 2557074 | 2571538 | 14464 | Predicted by at least one method | 2557410 | 2558978 | Phage T7 exclusion protein |
| 2557074 | 2571538 | 14464 | Predicted by at least one method | 2558993 | 2559355 | Phage T7 exclusion protein |
| 2557074 | 2571538 | 14464 | Predicted by at least one method | 2559720 | 2560250 | Phage T7 exclusion protein associated hypothetical protein |
| 2557074 | 2571538 | 14464 | Predicted by at least one method | 2560247 | 2561617 | FIG014574: hypothetical protein |
| 2557074 | 2571538 | 14464 | Predicted by at least one method | 2562084 | 2562386 | Putative deoxyribonuclease similar to YcfH, type 4 |
| 2557074 | 2571538 | 14464 | Predicted by at least one method | 2563836 | 2564162 | Phage integrase |
| 2557074 | 2571538 | 14464 | Predicted by at least one method | 2564501 | 2569630 | ATP-dependent DNA helicase RecQ |
| 2557074 | 2571538 | 14464 | Predicted by at least one method | 2569729 | 2569944 | VrII protein |
| 2557074 | 2571538 | 14464 | Predicted by at least one method | 2569973 | 2571979 | Type I restriction-modification system, DNA-methyltransferase subunit M |
| 2660496 | 2664751 | 4255 | Predicted by at least one method | 2660496 | 2660921 | General secretion pathway protein I |
| 2660496 | 2664751 | 4255 | Predicted by at least one method | 2660918 | 2661508 | Type II secretory pathway, pseudopilin PulG |
| 2660496 | 2664751 | 4255 | Predicted by at least one method | 2661564 | 2662577 | Type II secretory pathway, component PulK |
| 2660496 | 2664751 | 4255 | Predicted by at least one method | 2663224 | 2663607 | hypothetical protein |
| 2660496 | 2664751 | 4255 | Predicted by at least one method | 2663607 | 2664191 | FIG00959371: hypothetical protein |
| 2660496 | 2664751 | 4255 | Predicted by at least one method | 2664434 | 2664751 | FIG00956760: hypothetical protein |
| 2764602 | 2769537 | 4935 | Predicted by at least one method | 2764681 | 2765685 | Integrase |
| 2764602 | 2769537 | 4935 | Predicted by at least one method | 2766142 | 2766264 | hypothetical protein |
| 2764602 | 2769537 | 4935 | Predicted by at least one method | 2766257 | 2766736 | Phage protein |
| 2764602 | 2769537 | 4935 | Predicted by at least one method | 2766736 | 2768016 | Phage protein |
| 2764602 | 2769537 | 4935 | Predicted by at least one method | 2768114 | 2768620 | hypothetical protein |
| 2764602 | 2769537 | 4935 | Predicted by at least one method | 2768979 | 2769539 | Phage protein |
| 2764602 | 2769537 | 4935 | Predicted by at least one method | 2769536 | 2770162 | Phage-related protein |
| 3282382 | 3292424 | 10042 | Predicted by at least one method | 3281822 | 3282541 | Uncharacterized protein conserved in bacteria |
| 3282382 | 3292424 | 10042 | Predicted by at least one method | 3289853 | 3290875 | elements of external origin; transposon-related functions |
| 3282382 | 3292424 | 10042 | Predicted by at least one method | 3291562 | 3291912 | hypothetical protein |
| 3282541 | 3299331 | 16790 | Predicted by at least one method | 3289853 | 3290875 | elements of external origin; transposon-related functions |
| 3282541 | 3299331 | 16790 | Predicted by at least one method | 3291562 | 3291912 | hypothetical protein |
| 3282541 | 3299331 | 16790 | Predicted by at least one method | 3292986 | 3293207 | hypothetical protein |
| 3282541 | 3299331 | 16790 | Predicted by at least one method | 3293246 | 3294247 | Cointegrate resolution protein T |
| 3282541 | 3299331 | 16790 | Predicted by at least one method | 3294428 | 3295399 | Tn4652, cointegrate resolution protein S |
| 3282541 | 3299331 | 16790 | Predicted by at least one method | 3295429 | 3296172 | Bacteriophage protein gp37 |
| 3282541 | 3299331 | 16790 | Predicted by at least one method | 3296194 | 3297510 | hypothetical protein |
| 3282541 | 3299331 | 16790 | Predicted by at least one method | 3297922 | 3299331 | Homospermidine synthase |
| 3296194 | 3305099 | 8905 | Predicted by at least one method | 3296194 | 3297510 | hypothetical protein |
| 3296194 | 3305099 | 8905 | Predicted by at least one method | 3297922 | 3299331 | Homospermidine synthase |
| 3296194 | 3305099 | 8905 | Predicted by at least one method | 3301978 | 3303249 | Mobile element protein |
| 3296194 | 3305099 | 8905 | Predicted by at least one method | 3303843 | 3305099 | Heavy metal RND efflux outer membrane protein, CzcC family |
| 3296194 | 3305099 | 8905 | Predicted by at least one method | 3305096 | 3306577 | Cobalt/zinc/cadmium efflux RND transporter, membrane fusion protein, CzcB family |
| 3309732 | 3328937 | 19205 | Predicted by at least one method | 3306574 | 3309735 | Cobalt-zinc-cadmium resistance protein CzcA; Cation efflux system protein CusA |
| 3309732 | 3328937 | 19205 | Predicted by at least one method | 3309732 | 3310088 | hypothetical protein |
| 3309732 | 3328937 | 19205 | Predicted by at least one method | 3310514 | 3311200 | BNR repeat domain protein |
| 3309732 | 3328937 | 19205 | Predicted by at least one method | 3311295 | 3311957 | Putative protein-S-isoprenylcysteine methyltransferase |
| 3309732 | 3328937 | 19205 | Predicted by at least one method | 3312114 | 3312278 | hypothetical protein |
| 3309732 | 3328937 | 19205 | Predicted by at least one method | 3312382 | 3312504 | hypothetical protein |
| 3309732 | 3328937 | 19205 | Predicted by at least one method | 3312858 | 3312971 | hypothetical protein |
| 3309732 | 3328937 | 19205 | Predicted by at least one method | 3314027 | 3314245 | FIG00954364: hypothetical protein |
| 3309732 | 3328937 | 19205 | Predicted by at least one method | 3314435 | 3315754 | hypothetical protein |
| 3309732 | 3328937 | 19205 | Predicted by at least one method | 3315777 | 3316520 | Bacteriophage protein gp37 |
| 3309732 | 3328937 | 19205 | Predicted by at least one method | 3316550 | 3317521 | Tn4652, cointegrate resolution protein S |
| 3309732 | 3328937 | 19205 | Predicted by at least one method | 3317701 | 3318699 | Cointegrate resolution protein T |
| 3309732 | 3328937 | 19205 | Predicted by at least one method | 3318786 | 3319382 | hypothetical protein |
| 3309732 | 3328937 | 19205 | Predicted by at least one method | 3319471 | 3319842 | FIG00960741: hypothetical protein |
| 3309732 | 3328937 | 19205 | Predicted by at least one method | 3320774 | 3321208 | Mercuric resistance operon regulatory protein |
| 3309732 | 3328937 | 19205 | Predicted by at least one method | 3321280 | 3321630 | Mercuric transport protein, MerT |
| 3309732 | 3328937 | 19205 | Predicted by at least one method | 3321643 | 3321918 | Periplasmic mercury(+2) binding protein |
| 3309732 | 3328937 | 19205 | Predicted by at least one method | 3321990 | 3323675 | Mercuric ion reductase |
| 3309732 | 3328937 | 19205 | Predicted by at least one method | 3323693 | 3324058 | Mercuric resistance operon coregulator |
| 3309732 | 3328937 | 19205 | Predicted by at least one method | 3324055 | 3324291 | Mercuric transport protein, MerE |
| 3309732 | 3328937 | 19205 | Predicted by at least one method | 3324288 | 3325277 | Tn21 protein of unknown function Urf2 |
| 3309732 | 3328937 | 19205 | Predicted by at least one method | 3325408 | 3325968 | Transposon Tn21 resolvase |
| 3309732 | 3328937 | 19205 | Predicted by at least one method | 3325971 | 3328937 | Mobile element protein |
| 3309979 | 3316639 | 6660 | Predicted by at least one method | 3309732 | 3310088 | hypothetical protein |
| 3309979 | 3316639 | 6660 | Predicted by at least one method | 3310514 | 3311200 | BNR repeat domain protein |
| 3309979 | 3316639 | 6660 | Predicted by at least one method | 3311295 | 3311957 | Putative protein-S-isoprenylcysteine methyltransferase |
| 3309979 | 3316639 | 6660 | Predicted by at least one method | 3312114 | 3312278 | hypothetical protein |
| 3309979 | 3316639 | 6660 | Predicted by at least one method | 3312382 | 3312504 | hypothetical protein |
| 3309979 | 3316639 | 6660 | Predicted by at least one method | 3312858 | 3312971 | hypothetical protein |
| 3309979 | 3316639 | 6660 | Predicted by at least one method | 3314027 | 3314245 | FIG00954364: hypothetical protein |
| 3309979 | 3316639 | 6660 | Predicted by at least one method | 3314435 | 3315754 | hypothetical protein |
| 3309979 | 3316639 | 6660 | Predicted by at least one method | 3315777 | 3316520 | Bacteriophage protein gp37 |
| 3309979 | 3316639 | 6660 | Predicted by at least one method | 3316550 | 3317521 | Tn4652, cointegrate resolution protein S |
| 3317489 | 3336291 | 18802 | Predicted by at least one method | 3316550 | 3317521 | Tn4652, cointegrate resolution protein S |
| 3317489 | 3336291 | 18802 | Predicted by at least one method | 3317701 | 3318699 | Cointegrate resolution protein T |
| 3317489 | 3336291 | 18802 | Predicted by at least one method | 3318786 | 3319382 | hypothetical protein |
| 3317489 | 3336291 | 18802 | Predicted by at least one method | 3319471 | 3319842 | FIG00960741: hypothetical protein |
| 3317489 | 3336291 | 18802 | Predicted by at least one method | 3320774 | 3321208 | Mercuric resistance operon regulatory protein |
| 3317489 | 3336291 | 18802 | Predicted by at least one method | 3321280 | 3321630 | Mercuric transport protein, MerT |
| 3317489 | 3336291 | 18802 | Predicted by at least one method | 3321643 | 3321918 | Periplasmic mercury(+2) binding protein |
| 3317489 | 3336291 | 18802 | Predicted by at least one method | 3321990 | 3323675 | Mercuric ion reductase |
| 3317489 | 3336291 | 18802 | Predicted by at least one method | 3323693 | 3324058 | Mercuric resistance operon coregulator |
| 3317489 | 3336291 | 18802 | Predicted by at least one method | 3324055 | 3324291 | Mercuric transport protein, MerE |
| 3317489 | 3336291 | 18802 | Predicted by at least one method | 3324288 | 3325277 | Tn21 protein of unknown function Urf2 |
| 3317489 | 3336291 | 18802 | Predicted by at least one method | 3325408 | 3325968 | Transposon Tn21 resolvase |
| 3317489 | 3336291 | 18802 | Predicted by at least one method | 3325971 | 3328937 | Mobile element protein |
| 3317489 | 3336291 | 18802 | Predicted by at least one method | 3330769 | 3332055 | Phage replication initiation protein |
| 3317489 | 3336291 | 18802 | Predicted by at least one method | 3332403 | 3332753 | hypothetical protein |
| 3317489 | 3336291 | 18802 | Predicted by at least one method | 3332965 | 3333171 | hypothetical protein |
| 3317489 | 3336291 | 18802 | Predicted by at least one method | 3333322 | 3334776 | Probable coat protein A precursor |
| **3415786** | **3442204** | **26418** | **Predicted by at least one** **method** | **3415786** | **3415980** | **Copper chaperone** |
| **3415786** | **3442204** | **26418** | **Predicted by at least one method** | **3416055** | **3416813** | **hypothetical protein** |
| **3415786** | **3442204** | **26418** | **Predicted by at least one method** | **3417031** | **3417369** | **Ferredoxin, 2Fe-2S** |
| **3415786** | **3442204** | **26418** | **Predicted by at least one method** | **3417386** | **3417865** | **HTH-type transcriptional regulator cueR** |
| **3415786** | **3442204** | **26418** | **Predicted by at least one method** | **3417881** | **3420310** | **Lead, cadmium, zinc and mercury transporting ATPase; Copper-translocating P-type ATPase** |
| **3415786** | **3442204** | **26418** | **Predicted by at least one method** | **3420690** | **3420971** | **Transcriptional regulator, ArsR family** |
| **3415786** | **3442204** | **26418** | **Predicted by at least one method** | **3421010** | **3422320** | **Probable NreB protein** |
| **3415786** | **3442204** | **26418** | **Predicted by at least one method** | **3422330** | **3422587** | **hypothetical protein** |
| **3415786** | **3442204** | **26418** | **Predicted by at least one method** | **3422678** | **3422884** | **LysR family transcriptional regulator STM3121** |
| **3415786** | **3442204** | **26418** | **Predicted by at least one method** | **3422994** | **3424154** | **Bicyclomycin resistance protein** |
| **3415786** | **3442204** | **26418** | **Predicted by at least one method** | **3424440** | **3426302** | **Type IV secretory pathway, VirD2 components (relaxase)** |
| **3415786** | **3442204** | **26418** | **Predicted by at least one method** | **3426725** | **3427684** | **transposase InsA** |
| **3415786** | **3442204** | **26418** | **Predicted by at least one method** | **3428169** | **3429644** | **COG0488: ATPase components of ABC transporters with duplicated ATPase domains** |
| **3415786** | **3442204** | **26418** | **Predicted by at least one method** | **3429926** | **3431455** | **transposase InsA** |
| **3415786** | **3442204** | **26418** | **Predicted by at least one method** | **3431913** | **3432725** | **Beta-lactamase** |
| **3415786** | **3442204** | **26418** | **Predicted by at least one** **method** | **3433164** | **3434693** | **transposase InsA** |
| **3415786** | **3442204** | **26418** | **Predicted by at least one method** | **3434920** | **3435066** | **Heat shock protein 60 family chaperone GroEL** |
| **3415786** | **3442204** | **26418** | **Predicted by at least one method** | **3435368** | **3435625** | **NreA-like protein** |
| **3415786** | **3442204** | **26418** | **Predicted by at least one method** | **3435916** | **3436947** | **Cobalt-zinc-cadmium resistance protein CzcD** |
| **3415786** | **3442204** | **26418** | **Predicted by at least one method** | **3436991** | **3437368** | **Transcriptional regulator, ArsR family** |
| **3415786** | **3442204** | **26418** | **Predicted by at least one method** | **3437793** | **3439031** | **Probable Co/Zn/Cd efflux system membrane fusion protein / Multidrug efflux membrane fusion protein M** |
| **3415786** | **3442204** | **26418** | **Predicted by at least one method** | **3439028** | **3442204** | **RND multidrug efflux transporter; Acriflavin resistance protein** |
| 3753079 | 3757622 | 4543 | Predicted by at least one method | 3753482 | 3756742 | hypothetical protein |
| 4367437 | 4374562 | 7125 | Predicted by at least one method | 4367437 | 4367721 | FIG00953120: hypothetical protein |
| 4367437 | 4374562 | 7125 | Predicted by at least one method | 4368197 | 4368787 | Adenylylsulfate kinase |
| 4367437 | 4374562 | 7125 | Predicted by at least one method | 4370089 | 4370433 | probable glycosyl transferase |
| 4367437 | 4374562 | 7125 | Predicted by at least one method | 4370506 | 4371969 | hypothetical protein |
| 4367437 | 4374562 | 7125 | Predicted by at least one method | 4373078 | 4374562 | Twin-arginine translocation protein TatC |
| 4368021 | 4387227 | 19206 | Predicted by at least one method | 4368197 | 4368787 | Adenylylsulfate kinase |
| 4368021 | 4387227 | 19206 | Predicted by at least one method | 4370089 | 4370433 | probable glycosyl transferase |
| 4368021 | 4387227 | 19206 | Predicted by at least one method | 4370506 | 4371969 | hypothetical protein |
| 4368021 | 4387227 | 19206 | Predicted by at least one method | 4373078 | 4374562 | Twin-arginine translocation protein TatC |
| 4368021 | 4387227 | 19206 | Predicted by at least one method | 4374659 | 4375387 | hypothetical protein |
| 4368021 | 4387227 | 19206 | Predicted by at least one method | 4375404 | 4377095 | hypothetical protein |
| 4368021 | 4387227 | 19206 | Predicted by at least one method | 4377107 | 4378090 | SN-glycerol-3-phosphate transport ATP-binding protein UgpC (TC 3.A.1.1.3) |
| 4368021 | 4387227 | 19206 | Predicted by at least one method | 4378395 | 4379978 | Lipid carrier : UDP-N-acetylgalactosaminyltransferase / Alpha-1,3-N-acetylgalactosamine transferase |
| 4368021 | 4387227 | 19206 | Predicted by at least one method | 4380320 | 4380436 | hypothetical protein |
| 4368021 | 4387227 | 19206 | Predicted by at least one method | 4380704 | 4382338 | hypothetical protein |
| 4368021 | 4387227 | 19206 | Predicted by at least one method | 4383028 | 4385037 | General secretion pathway protein D |
| 4368021 | 4387227 | 19206 | Predicted by at least one method | 4385291 | 4385578 | hypothetical protein |
| 4368021 | 4387227 | 19206 | Predicted by at least one method | 4386143 | 4387054 | Transcriptional regulator, AraC family |
| 4368021 | 4387227 | 19206 | Predicted by at least one method | 4387201 | 4388034 | Short-chain dehydrogenase/reductase SDR |
| 4824516 | 4829497 | 4981 | Predicted by at least one method | 4825034 | 4825393 | hypothetical protein |
| 4824516 | 4829497 | 4981 | Predicted by at least one method | 4825582 | 4826148 | Phage protein |
| 4824516 | 4829497 | 4981 | Predicted by at least one method | 4826135 | 4826602 | Phage protein |
| 4824516 | 4829497 | 4981 | Predicted by at least one method | 4826604 | 4826822 | hypothetical protein |
| 4824516 | 4829497 | 4981 | Predicted by at least one method | 4826824 | 4827219 | inhibition of exonuclease digestion of Mu DNA |
| 4824516 | 4829497 | 4981 | Predicted by at least one method | 4827221 | 4827844 | hypothetical protein |
| 4824516 | 4829497 | 4981 | Predicted by at least one method | 4827837 | 4828037 | Phage protein |
| 4824516 | 4829497 | 4981 | Predicted by at least one method | 4828641 | 4828757 | hypothetical protein |
| 4824516 | 4829497 | 4981 | Predicted by at least one method | 4828854 | 4829042 | hypothetical protein |
| 4824516 | 4829497 | 4981 | Predicted by at least one method | 4829081 | 4829422 | Phage protein |
| 4824516 | 4829497 | 4981 | Predicted by at least one method | 4829424 | 4830590 | Eha |
| 4838194 | 4862792 | 24598 | Predicted by at least one method | 4837679 | 4838302 | Lipoprotein, phage-associated |
| 4838194 | 4862792 | 24598 | Predicted by at least one method | 4838302 | 4838622 | hypothetical protein |
| 4838194 | 4862792 | 24598 | Predicted by at least one method | 4838619 | 4838921 | hypothetical protein |
| 4838194 | 4862792 | 24598 | Predicted by at least one method | 4838924 | 4839472 | Phage terminase, small subunit |
| 4838194 | 4862792 | 24598 | Predicted by at least one method | 4839474 | 4841147 | Phage terminase, large subunit |
| 4838194 | 4862792 | 24598 | Predicted by at least one method | 4841150 | 4842715 | Mu-like prophage FluMu protein gp29 |
| 4838194 | 4862792 | 24598 | Predicted by at least one method | 4842705 | 4843943 | Phage (Mu-like) virion morphogenesis protein |
| 4838194 | 4862792 | 24598 | Predicted by at least one method | 4844032 | 4844520 | possible bacteriophage Mu G-like protein |
| 4838194 | 4862792 | 24598 | Predicted by at least one method | 4844731 | 4845840 | Mu-like prophage FluMu I protein |
| 4838194 | 4862792 | 24598 | Predicted by at least one method | 4845846 | 4846250 | Phage protein |
| 4838194 | 4862792 | 24598 | Predicted by at least one method | 4846265 | 4847161 | Phage major capsid protein |
| 4838194 | 4862792 | 24598 | Predicted by at least one method | 4847393 | 4847608 | Phage protein |
| 4838194 | 4862792 | 24598 | Predicted by at least one method | 4847611 | 4848126 | Phage protein |
| 4838194 | 4862792 | 24598 | Predicted by at least one method | 4848123 | 4848575 | Phage protein |
| 4838194 | 4862792 | 24598 | Predicted by at least one method | 4848572 | 4848775 | hypothetical protein |
| 4838194 | 4862792 | 24598 | Predicted by at least one method | 4848782 | 4849522 | Phage protein |
| 4838194 | 4862792 | 24598 | Predicted by at least one method | 4849624 | 4850007 | Phage protein |
| 4838194 | 4862792 | 24598 | Predicted by at least one method | 4850261 | 4853893 | Phage tail length tape-measure protein |
| 4838194 | 4862792 | 24598 | Predicted by at least one method | 4853893 | 4854852 | Phage protein |
| 4838194 | 4862792 | 24598 | Predicted by at least one method | 4855814 | 4857484 | Phage protein |
| 4838194 | 4862792 | 24598 | Predicted by at least one method | 4857738 | 4858289 | Phage FAD/FMN-containing dehydrogenase |
| 4838194 | 4862792 | 24598 | Predicted by at least one method | 4858299 | 4858529 | Phage protein |
| 4838194 | 4862792 | 24598 | Predicted by at least one method | 4858830 | 4860953 | Phage protein |
| 4838194 | 4862792 | 24598 | Predicted by at least one method | 4860950 | 4862098 | Phage protein |
| 4838194 | 4862792 | 24598 | Predicted by at least one method | 4862095 | 4862382 | Phage protein |
| 4838194 | 4862792 | 24598 | Predicted by at least one method | 4862543 | 4862680 | Phage protein |
| 4877189 | 4881287 | 4098 | Predicted by at least one method | 4877189 | 4877437 | FIG00960798: hypothetical protein |
| 4877189 | 4881287 | 4098 | Predicted by at least one method | 4877440 | 4879170 | Protein with ParB-like nuclease domain in PFGI-1-like cluster |
| 4877189 | 4881287 | 4098 | Predicted by at least one method | 4879198 | 4879965 | FIG004780: hypothetical protein in PFGI-1-like cluster |
| 4877189 | 4881287 | 4098 | Predicted by at least one method | 4879962 | 4881287 | FIG141751: hypothetical protein in PFGI-1-like cluster |
| 4888002 | 4892800 | 4798 | Predicted by at least one method | 4888002 | 4888154 | Haemophilus-specific protein, uncharacterized |
| 4888002 | 4892800 | 4798 | Predicted by at least one method | 4888188 | 4888754 | Haemophilus-specific protein, uncharacterized |
| 4888002 | 4892800 | 4798 | Predicted by at least one method | 4888871 | 4889476 | Homoserine/homoserine lactone efflux protein |
| 4888002 | 4892800 | 4798 | Predicted by at least one method | 4889712 | 4890467 | Transcriptional regulator, LysR family |
| 4888002 | 4892800 | 4798 | Predicted by at least one method | 4890911 | 4892800 | hypothetical protein in PFGI-1-like cluster |
| 4888002 | 4892800 | 4798 | Predicted by at least one method | 4892797 | 4894770 | DNA/RNA helicase in PFGI-1-like cluster |
| 4926918 | 4944808 | 17890 | Predicted by at least one method | 4925894 | 4926934 | Inner membrane component of tripartite multidrug resistance system |
| 4926918 | 4944808 | 17890 | Predicted by at least one method | 4926918 | 4928954 | Membrane fusion component of tripartite multidrug resistance system |
| 4926918 | 4944808 | 17890 | Predicted by at least one method | 4929370 | 4929969 | Transcriptional regulator, LysR family |
| 4926918 | 4944808 | 17890 | Predicted by at least one method | 4929930 | 4930046 | hypothetical protein |
| 4926918 | 4944808 | 17890 | Predicted by at least one method | 4930205 | 4930657 | PrnC |
| 4926918 | 4944808 | 17890 | Predicted by at least one method | 4932274 | 4933722 | Multidrug resistance protein B |
| 4926918 | 4944808 | 17890 | Predicted by at least one method | 4933810 | 4934277 | Membrane fusion component of tripartite multidrug resistance system |
| 4926918 | 4944808 | 17890 | Predicted by at least one method | 4935008 | 4936465 | Outer membrane component of tripartite multidrug resistance system |
| 4926918 | 4944808 | 17890 | Predicted by at least one method | 4937342 | 4937803 | Pirin |
| 4926918 | 4944808 | 17890 | Predicted by at least one method | 4940332 | 4941912 | Catalase |
| 4926918 | 4944808 | 17890 | Predicted by at least one method | 4942286 | 4943233 | Mobile element protein |
| 4926918 | 4944808 | 17890 | Predicted by at least one method | 4943362 | 4944042 | Methyl-accepting chemotaxis protein |
| 4926918 | 4944808 | 17890 | Predicted by at least one method | 4944053 | 4944808 | probable exported protein STY4558 |
| 4926918 | 4944808 | 17890 | Predicted by at least one method | 4944793 | 4945374 | Soluble lytic murein transglycosylase and related regulatory proteins (some contain LysM/invasin dom |
| 4929252 | 4942176 | 12924 | Predicted by at least one method | 4929370 | 4929969 | Transcriptional regulator, LysR family |
| 4929252 | 4942176 | 12924 | Predicted by at least one method | 4929930 | 4930046 | hypothetical protein |
| 4929252 | 4942176 | 12924 | Predicted by at least one method | 4930205 | 4930657 | PrnC |
| 4929252 | 4942176 | 12924 | Predicted by at least one method | 4932274 | 4933722 | Multidrug resistance protein B |
| 4929252 | 4942176 | 12924 | Predicted by at least one method | 4933810 | 4934277 | Membrane fusion component of tripartite multidrug resistance system |
| 4929252 | 4942176 | 12924 | Predicted by at least one method | 4935008 | 4936465 | Outer membrane component of tripartite multidrug resistance system |
| 4929252 | 4942176 | 12924 | Predicted by at least one method | 4937342 | 4937803 | Pirin |
| 4929252 | 4942176 | 12924 | Predicted by at least one method | 4940332 | 4941912 | Catalase |
| 5146667 | 5151453 | 4786 | Predicted by at least one method | 5145495 | 5146751 | Phage integrase |
| 5146667 | 5151453 | 4786 | Predicted by at least one method | 5147106 | 5149706 | DEAD/DEAH box helicase domain protein |
| 5146667 | 5151453 | 4786 | Predicted by at least one method | 5149706 | 5149939 | bll2187; hypothetical protein |
| 5146667 | 5151453 | 4786 | Predicted by at least one method | 5150951 | 5151223 | FIG00957676: hypothetical protein |
| 5146667 | 5151453 | 4786 | Predicted by at least one method | 5151318 | 5152286 | Mycobacteriophage Barnyard protein gp56 |
| 5286995 | 5294366 | 7371 | Predicted by at least one method | 5286983 | 5287303 | Phage protein |
| 5286995 | 5294366 | 7371 | Predicted by at least one method | 5288289 | 5288453 | hypothetical protein |
| 6237355 | 6243264 | 5909 | Predicted by at least one method | 6236450 | 6237358 | TniB NTP-binding protein |
| 6237355 | 6243264 | 5909 | Predicted by at least one method | 6237355 | 6237474 | TniQ |
| 6237355 | 6243264 | 5909 | Predicted by at least one method | 6242509 | 6243264 | TniQ |
| 6403449 | 6413362 | 9913 | Predicted by at least one method | 6402981 | 6404762 | Phage terminase, large subunit (P2-like 5'-extended COS ends) |
| 6403449 | 6413362 | 9913 | Predicted by at least one method | 6404897 | 6405718 | Phage capsid scaffolding protein |
| 6403449 | 6413362 | 9913 | Predicted by at least one method | 6405754 | 6406770 | Phage major capsid protein |
| 6403449 | 6413362 | 9913 | Predicted by at least one method | 6406776 | 6407477 | Phage terminase, small subunit |
| 6403449 | 6413362 | 9913 | Predicted by at least one method | 6407581 | 6408042 | Phage head completion-stabilization protein |
| 6403449 | 6413362 | 9913 | Predicted by at least one method | 6408042 | 6408254 | Phage tail X |
| 6403449 | 6413362 | 9913 | Predicted by at least one method | 6408279 | 6408632 | hypothetical protein |
| 6403449 | 6413362 | 9913 | Predicted by at least one method | 6408634 | 6408906 | Phage protein |
| 6403449 | 6413362 | 9913 | Predicted by at least one method | 6408903 | 6409709 | Putative phage-encoded peptidoglycan binding protein |
| 6403449 | 6413362 | 9913 | Predicted by at least one method | 6409706 | 6410167 | Phage endolysin |
| 6403449 | 6413362 | 9913 | Predicted by at least one method | 6410245 | 6410781 | Phage tail completion protein |
| 6403449 | 6413362 | 9913 | Predicted by at least one method | 6410774 | 6411244 | Phage tail completion protein |
| 6403449 | 6413362 | 9913 | Predicted by at least one method | 6412086 | 6412658 | Baseplate assembly protein V |
| 6403449 | 6413362 | 9913 | Predicted by at least one method | 6412655 | 6412999 | Phage baseplate assembly protein |
| 6403449 | 6413362 | 9913 | Predicted by at least one method | 6412996 | 6413913 | Baseplate assembly protein J |
| 6414976 | 6431666 | 16690 | Predicted by at least one method | 6414524 | 6416341 | Phage tail fiber protein |
| 6414976 | 6431666 | 16690 | Predicted by at least one method | 6416351 | 6416785 | Phage tail fiber assembly protein |
| 6414976 | 6431666 | 16690 | Predicted by at least one method | 6416886 | 6418061 | Phage major tail sheath protein |
| 6414976 | 6431666 | 16690 | Predicted by at least one method | 6418118 | 6418633 | Phage major tail tube protein |
| 6414976 | 6431666 | 16690 | Predicted by at least one method | 6418688 | 6419017 | Phage tail fibers |
| 6414976 | 6431666 | 16690 | Predicted by at least one method | 6419026 | 6419145 | hypothetical protein |
| 6414976 | 6431666 | 16690 | Predicted by at least one method | 6419135 | 6421894 | Phage tail length tape-measure protein |
| 6414976 | 6431666 | 16690 | Predicted by at least one method | 6421900 | 6422340 | Phage tail protein |
| 6414976 | 6431666 | 16690 | Predicted by at least one method | 6422337 | 6423614 | Phage protein D |
| 6414976 | 6431666 | 16690 | Predicted by at least one method | 6425085 | 6425435 | Phage protein |
| 6414976 | 6431666 | 16690 | Predicted by at least one method | 6425488 | 6425604 | Phage protein |
| 6414976 | 6431666 | 16690 | Predicted by at least one method | 6426181 | 6426393 | hypothetical protein |
| 6414976 | 6431666 | 16690 | Predicted by at least one method | 6426456 | 6426893 | Orf33 |
| 6414976 | 6431666 | 16690 | Predicted by at least one method | 6427180 | 6427530 | hypothetical protein |
| 6414976 | 6431666 | 16690 | Predicted by at least one method | 6427607 | 6427840 | Phage protein |
| 6414976 | 6431666 | 16690 | Predicted by at least one method | 6427837 | 6430557 | Phage-related protein |
| 6414976 | 6431666 | 16690 | Predicted by at least one method | 6430602 | 6430955 | hypothetical protein |
| 6414976 | 6431666 | 16690 | Predicted by at least one method | 6430967 | 6431173 | FIG00957497: hypothetical protein |
| 6414976 | 6431666 | 16690 | Predicted by at least one method | 6431477 | 6433387 | C-5 cytosine-specific DNA methylase |
| 6431894 | 6436329 | 4435 | Predicted by at least one method | 6431477 | 6433387 | C-5 cytosine-specific DNA methylase |
| 6431894 | 6436329 | 4435 | Predicted by at least one method | 6433384 | 6434616 | 3'-phosphoadenosine 5'-phosphosulfate sulfurtransferase DndC |
| 6431894 | 6436329 | 4435 | Predicted by at least one method | 6435021 | 6436169 | Mobile element protein |
| 6431894 | 6436329 | 4435 | Predicted by at least one method | 6436288 | 6436485 | hypothetical protein |

**Table S4:**
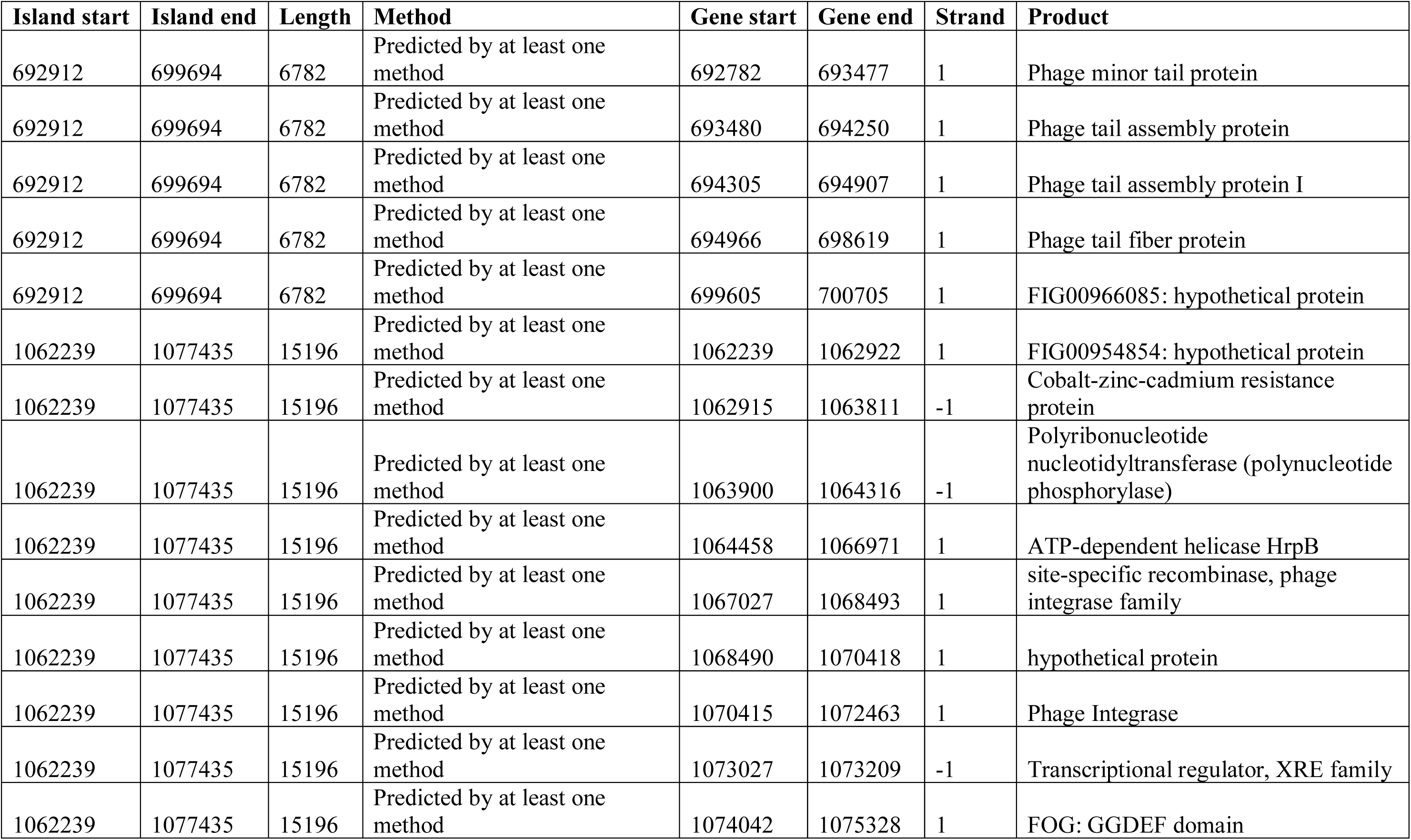
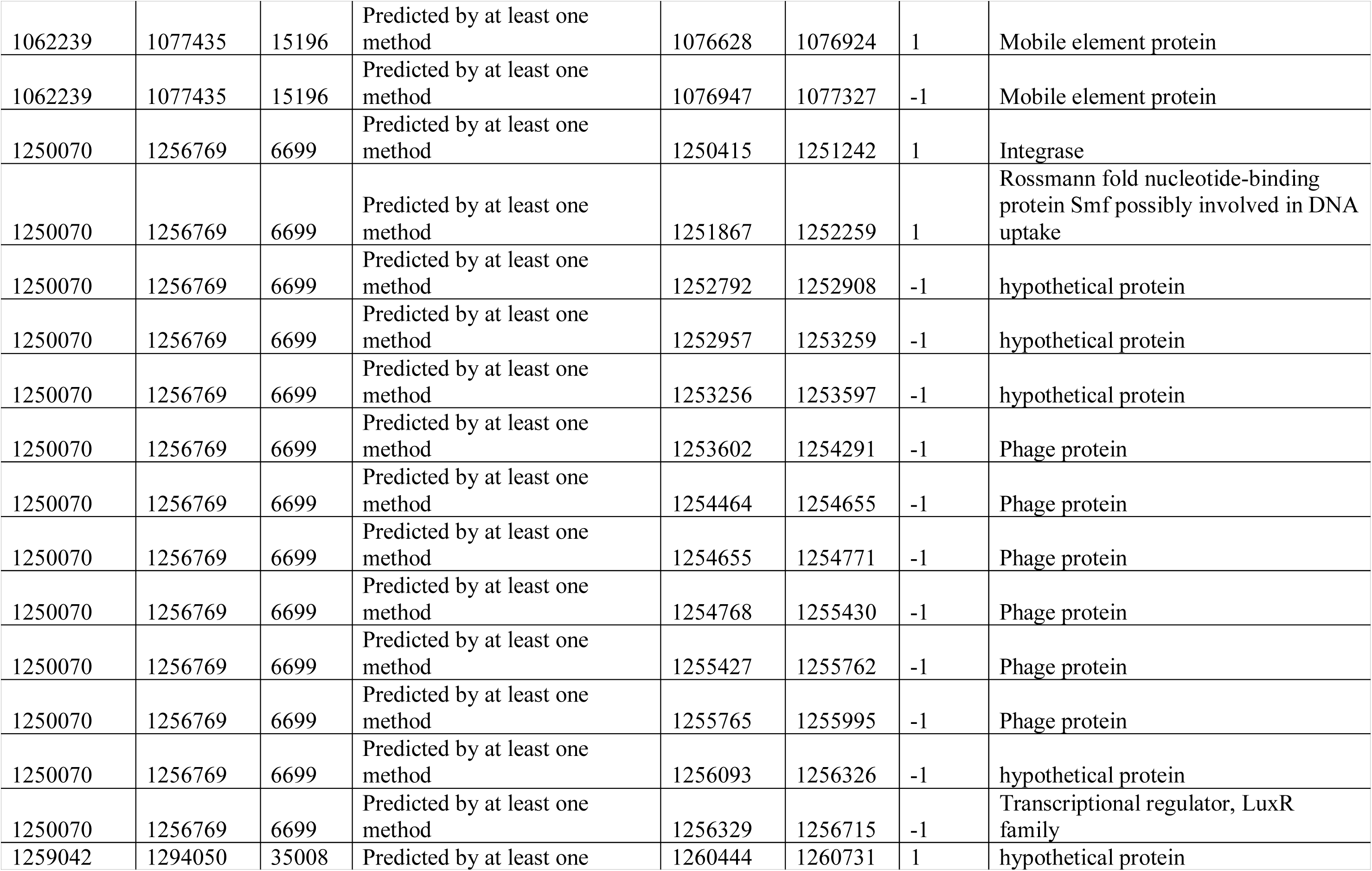
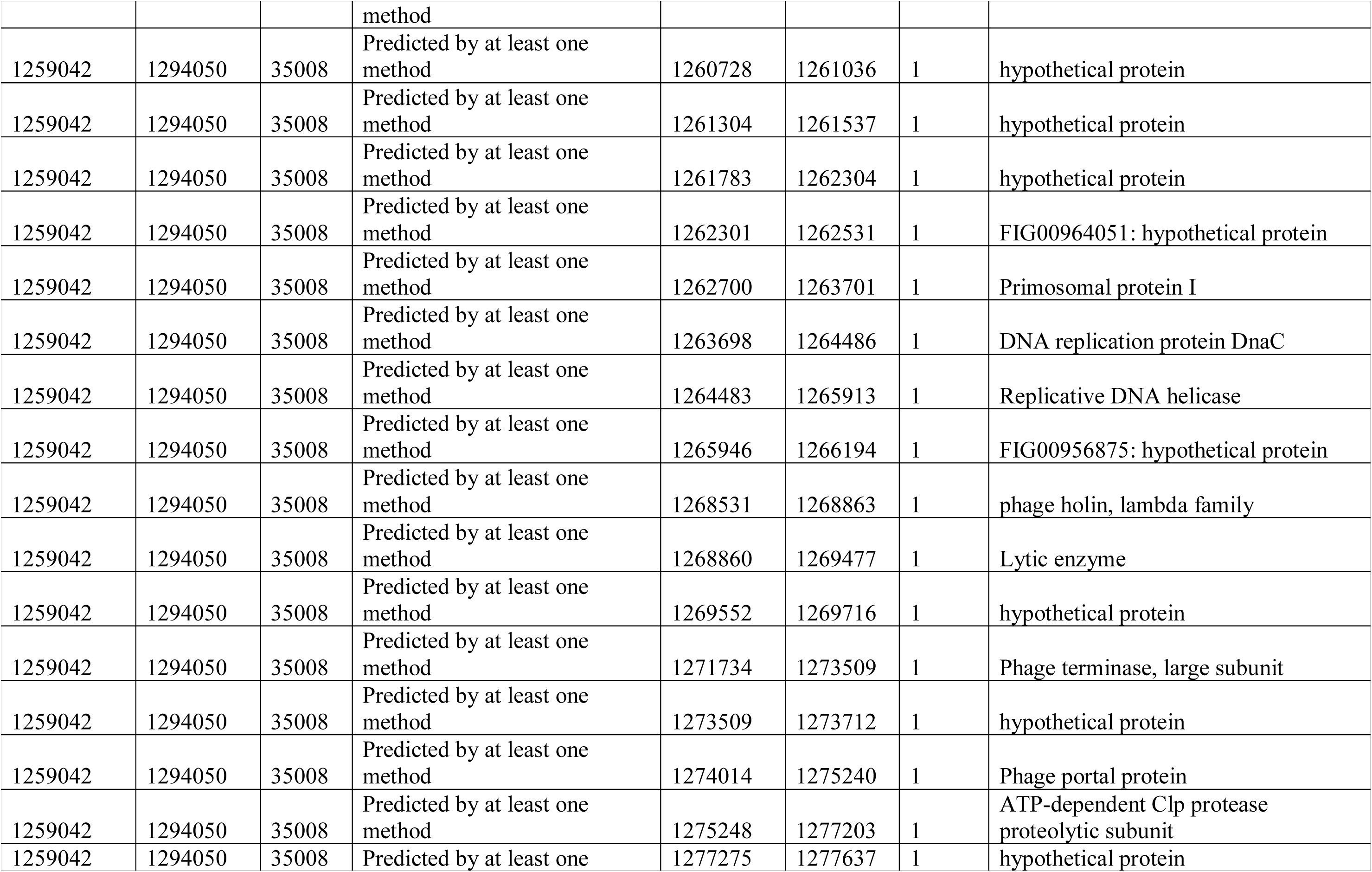
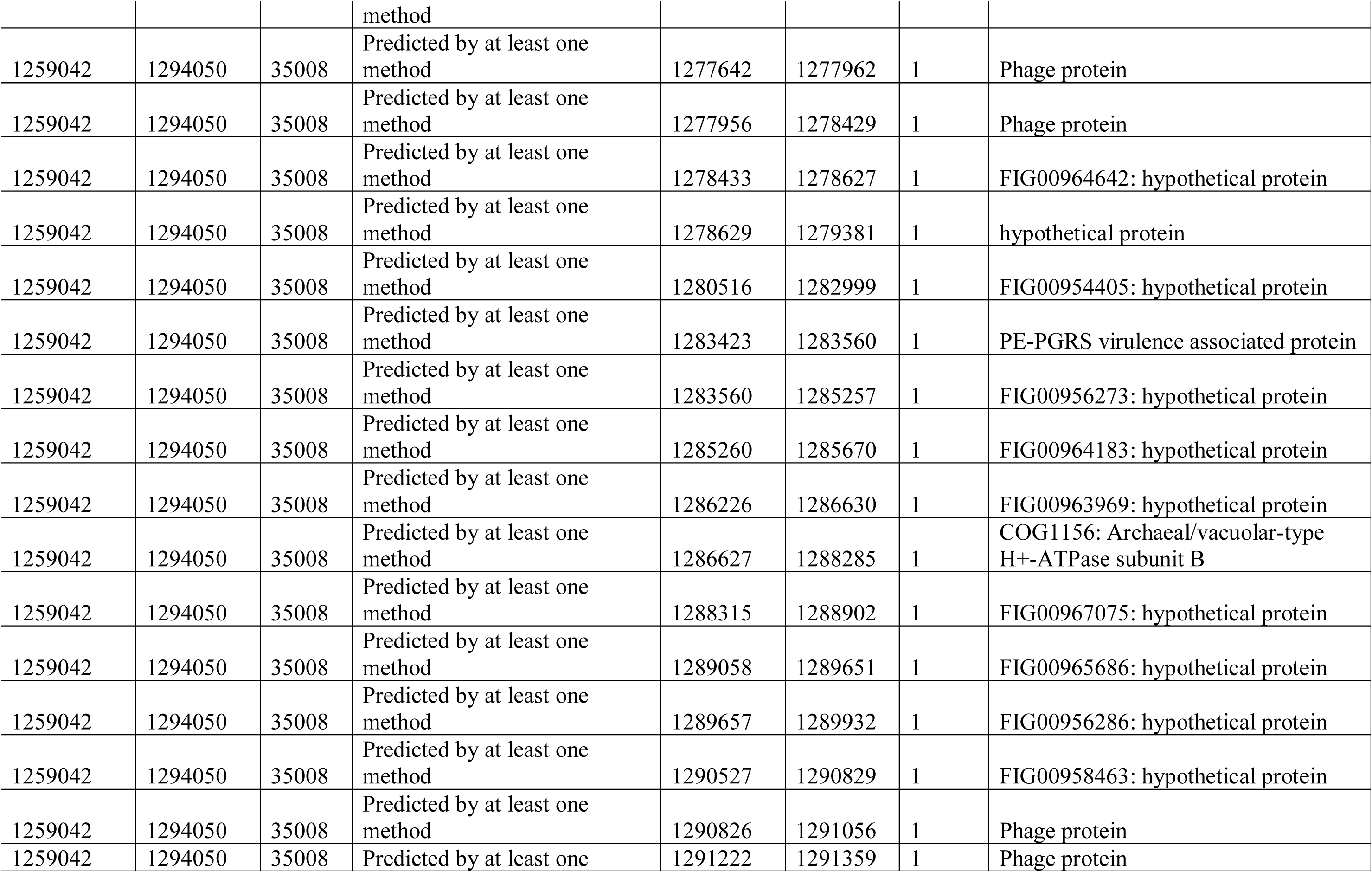
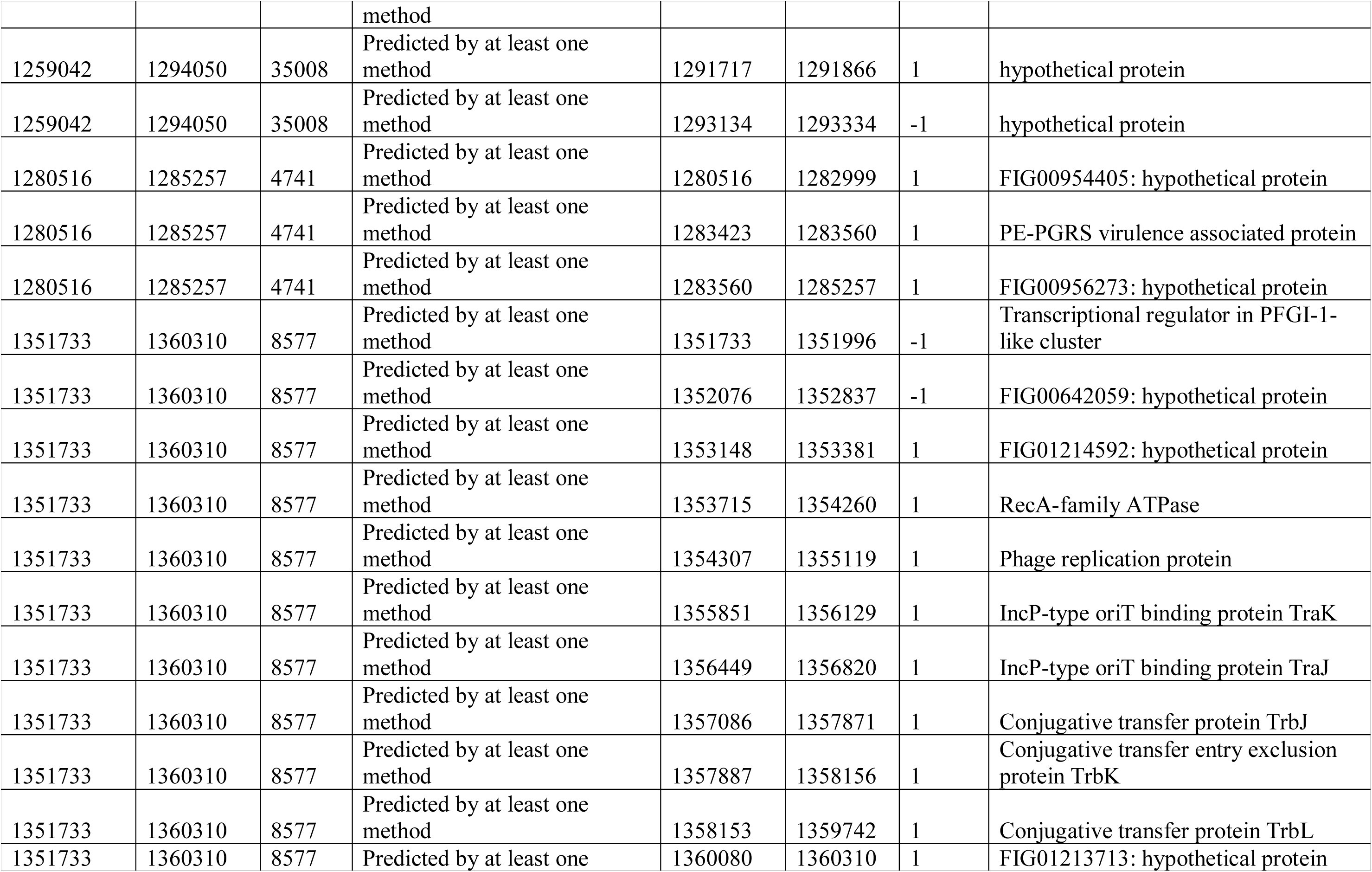
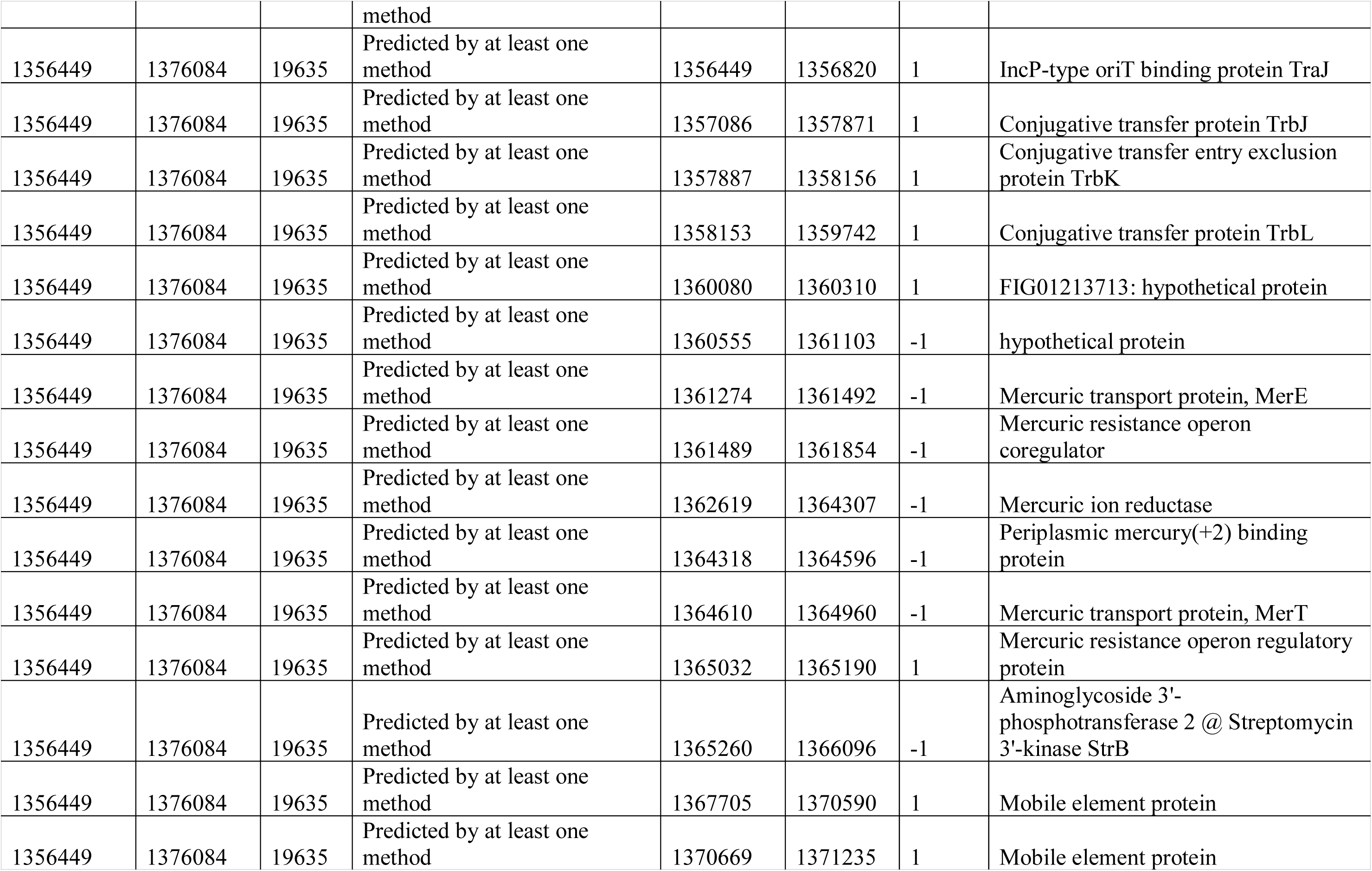
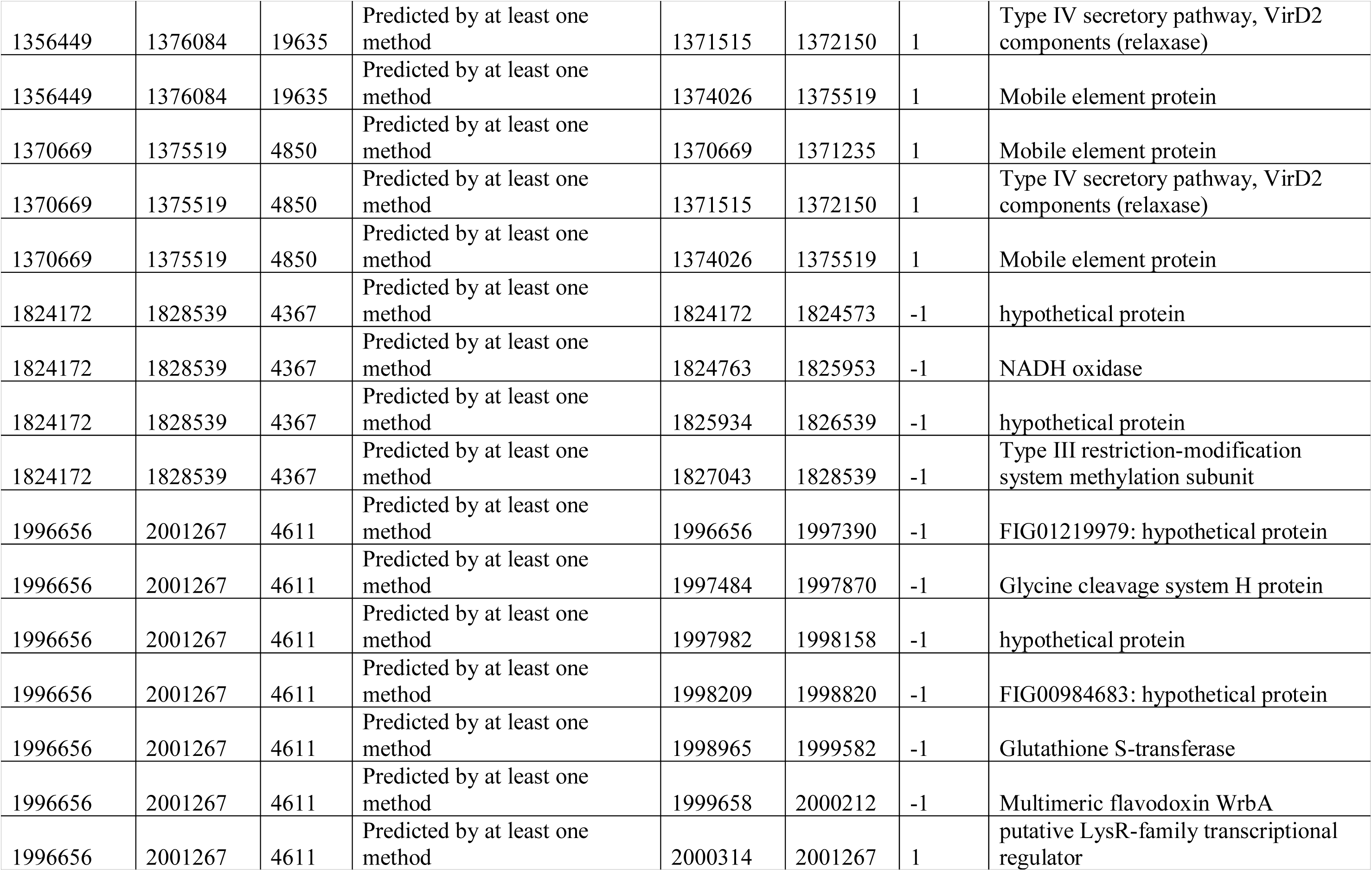
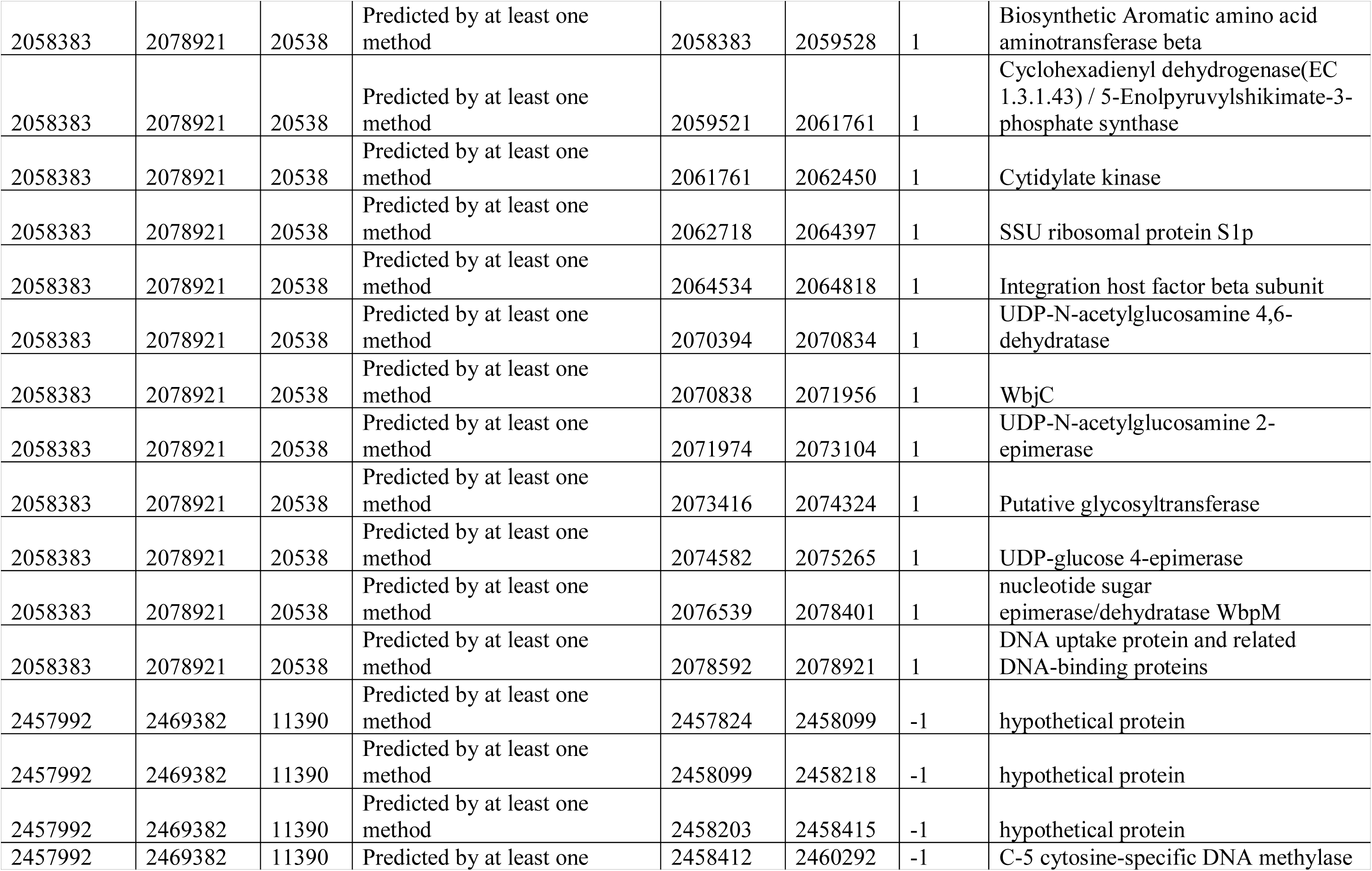
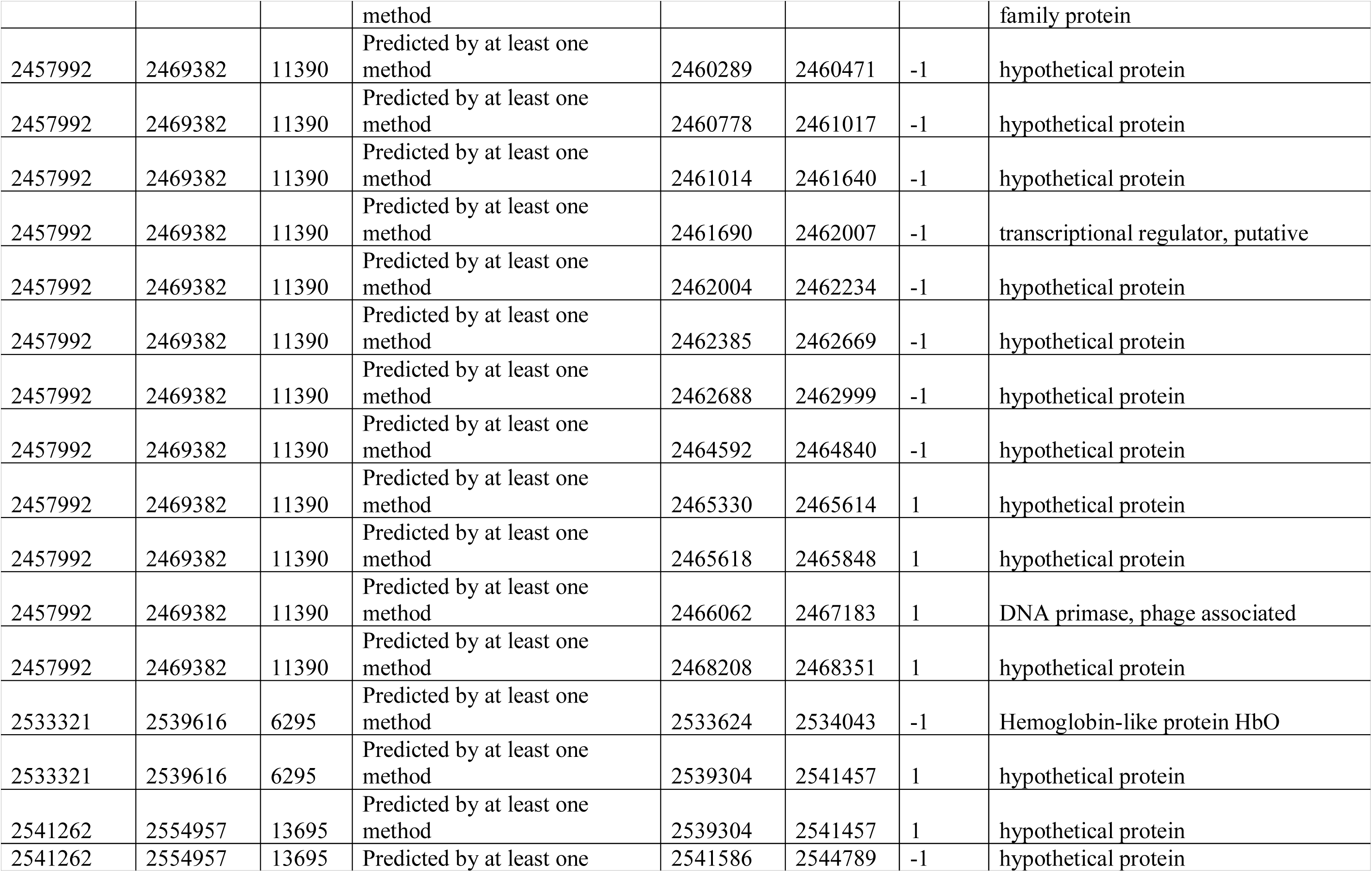
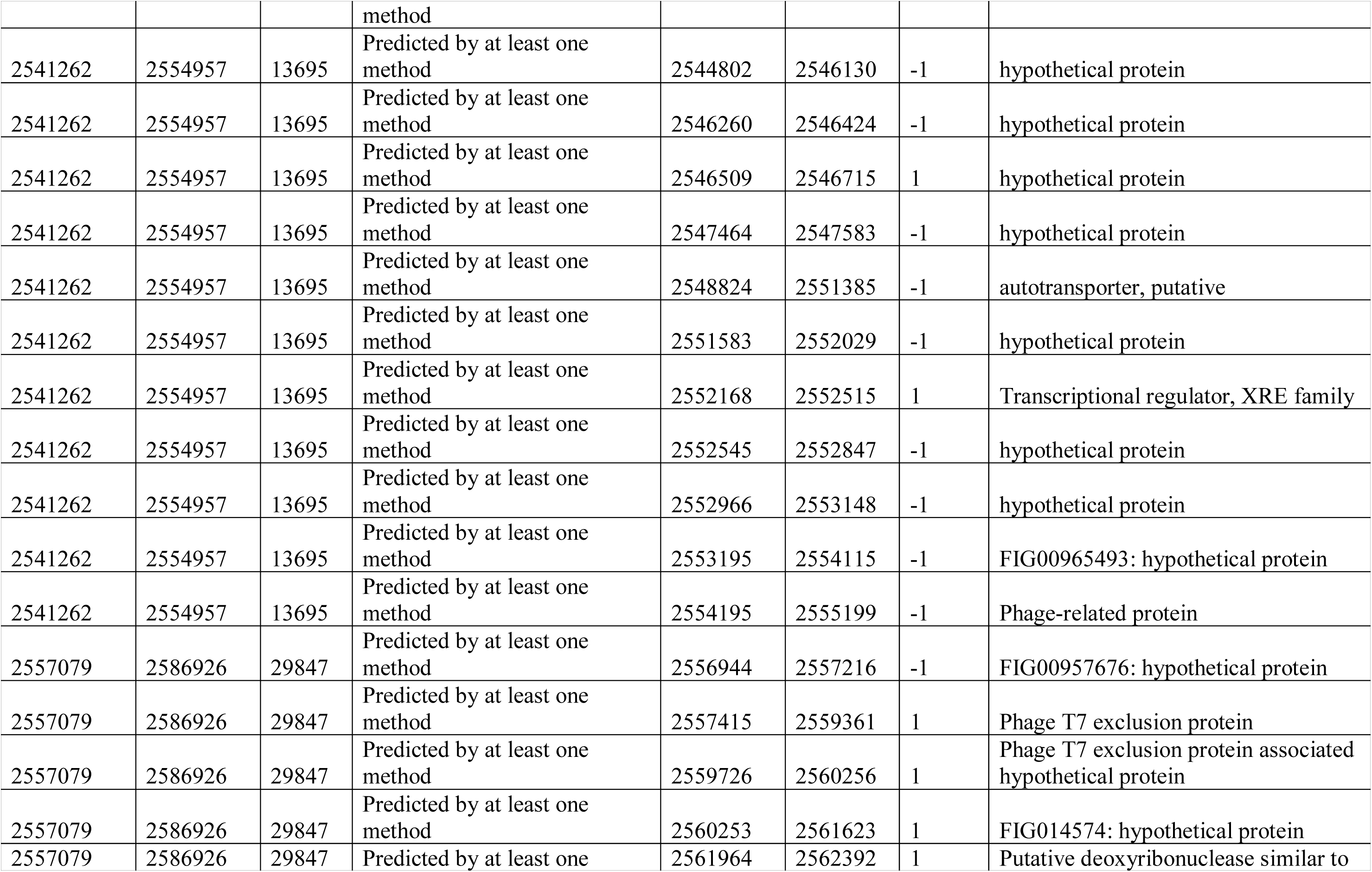
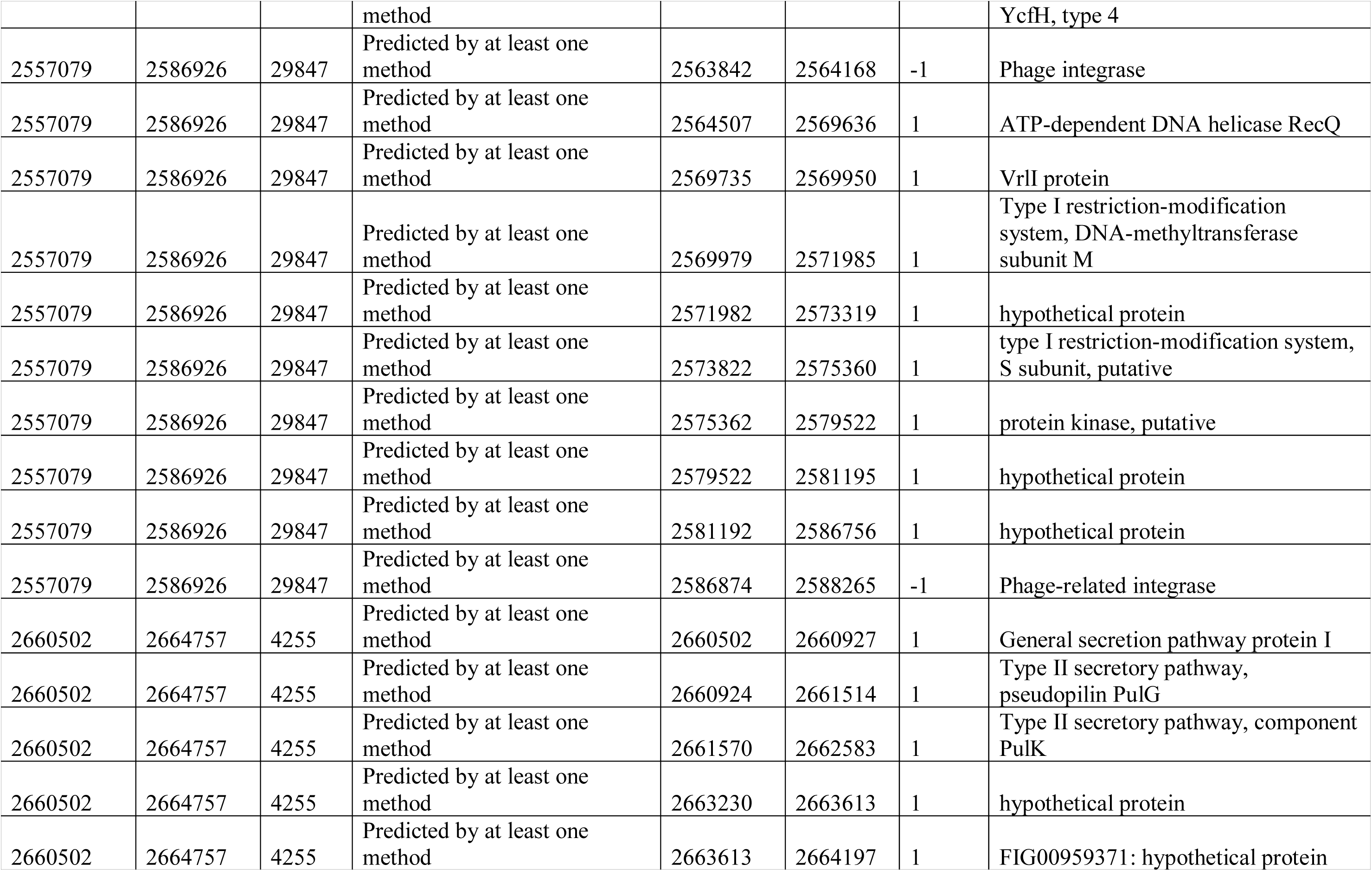
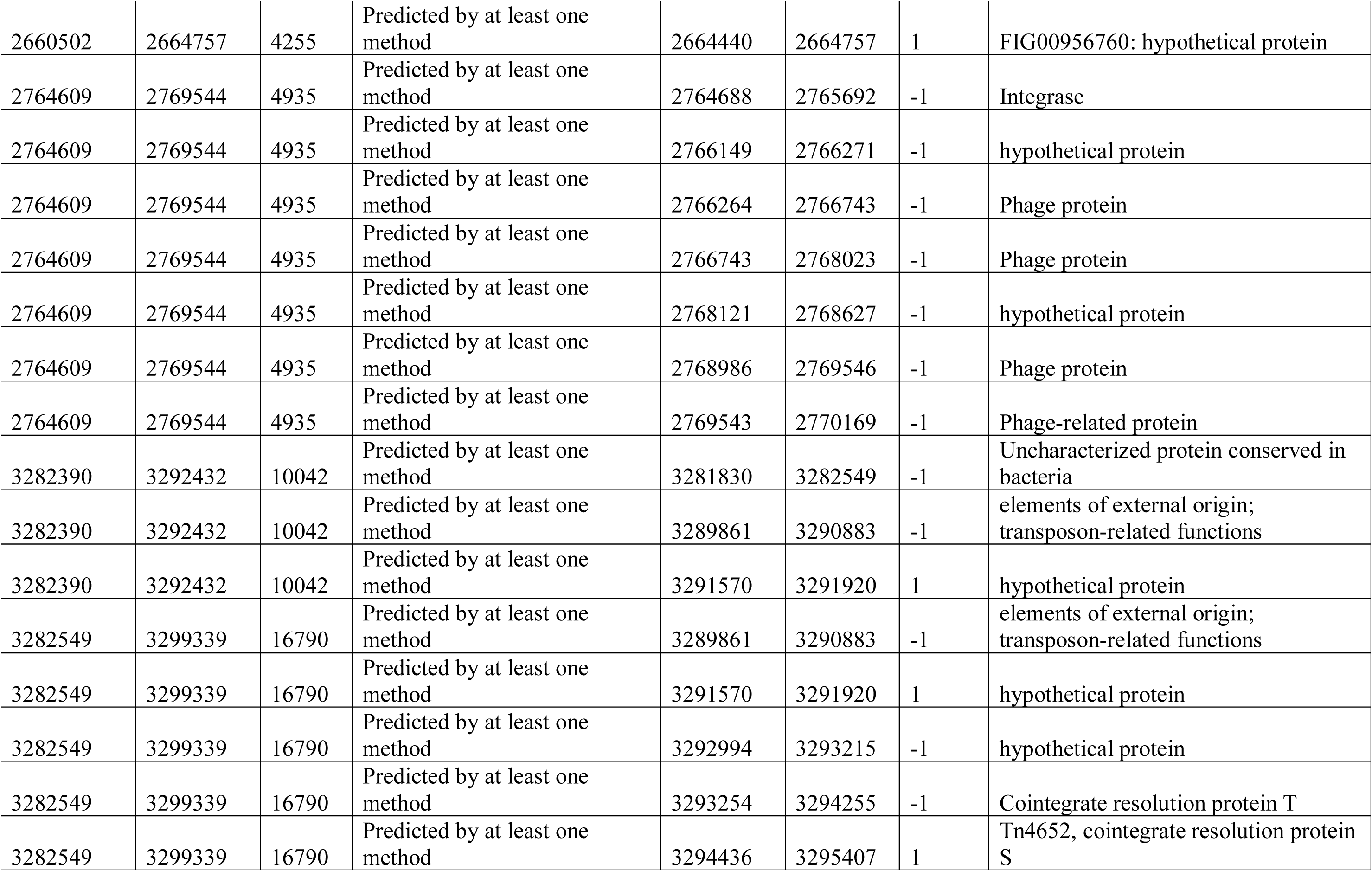
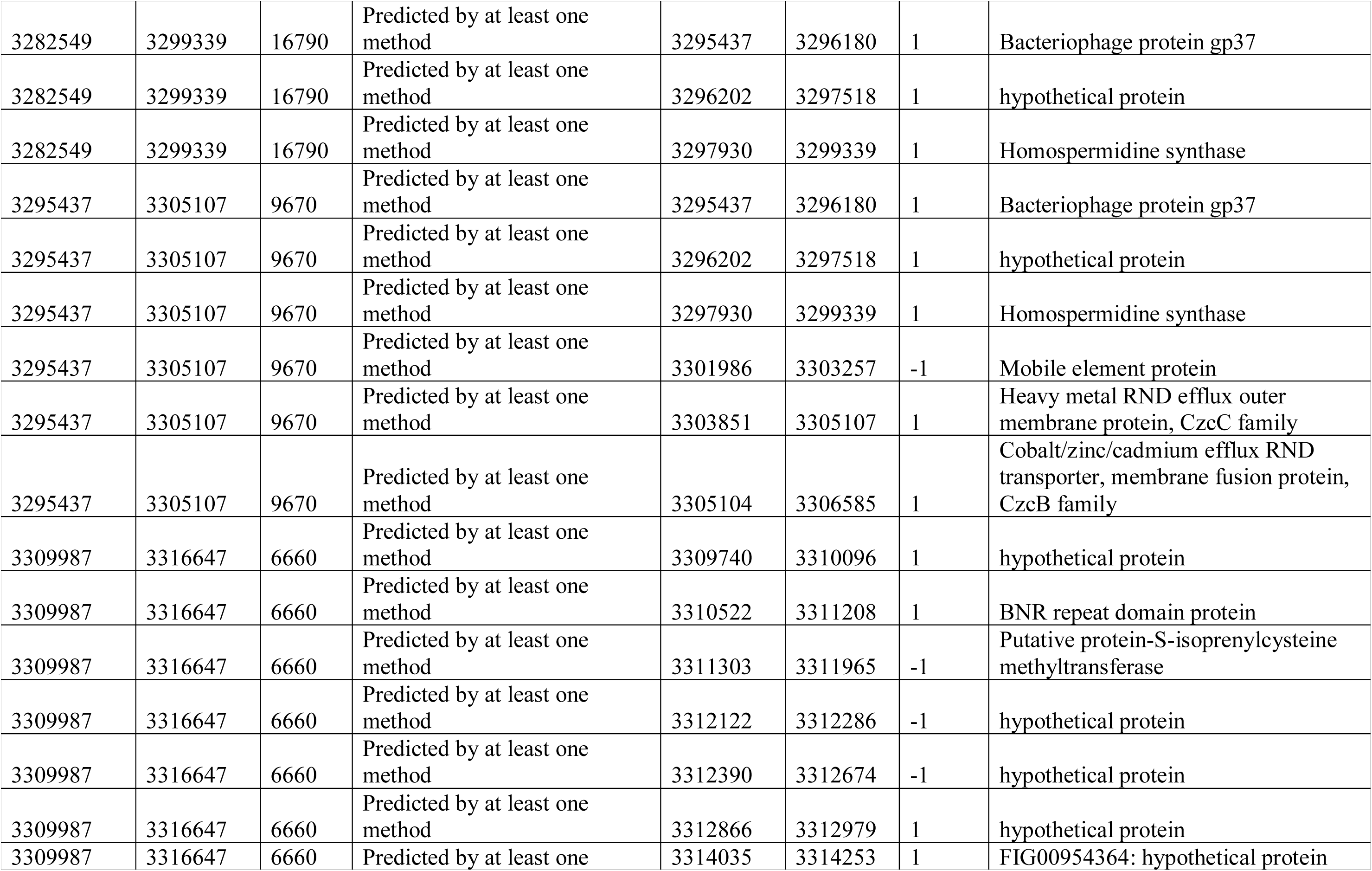
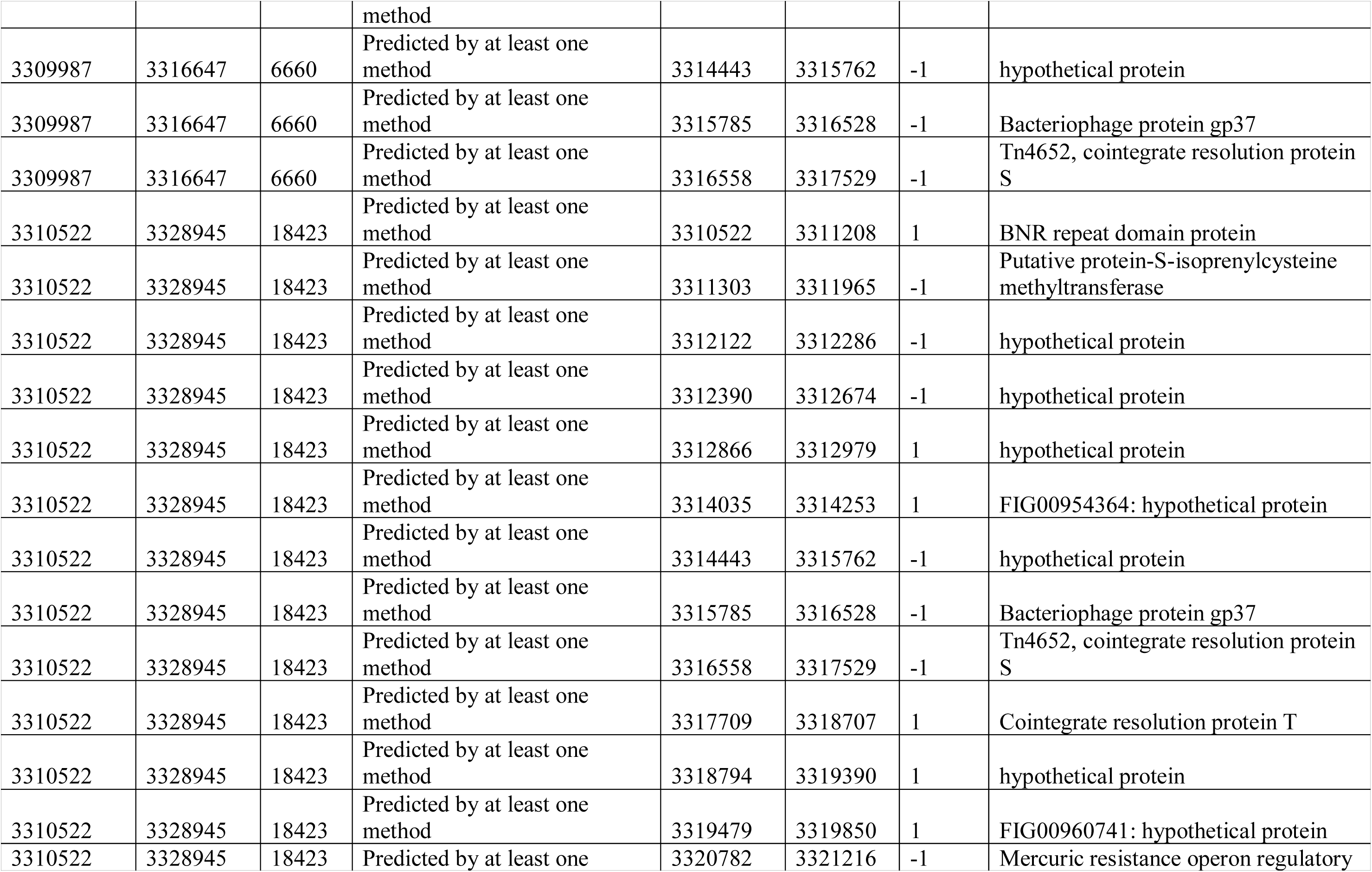
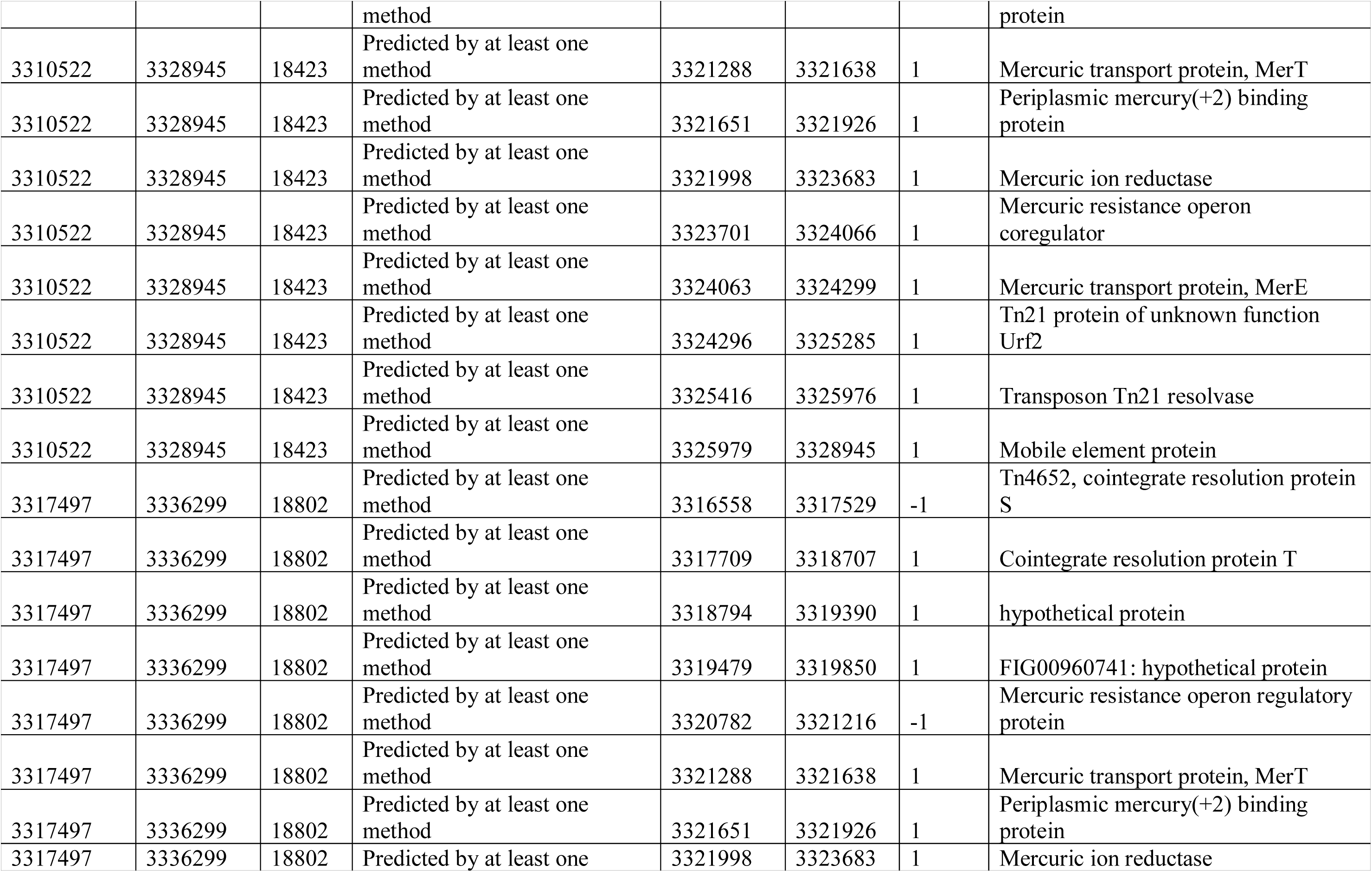
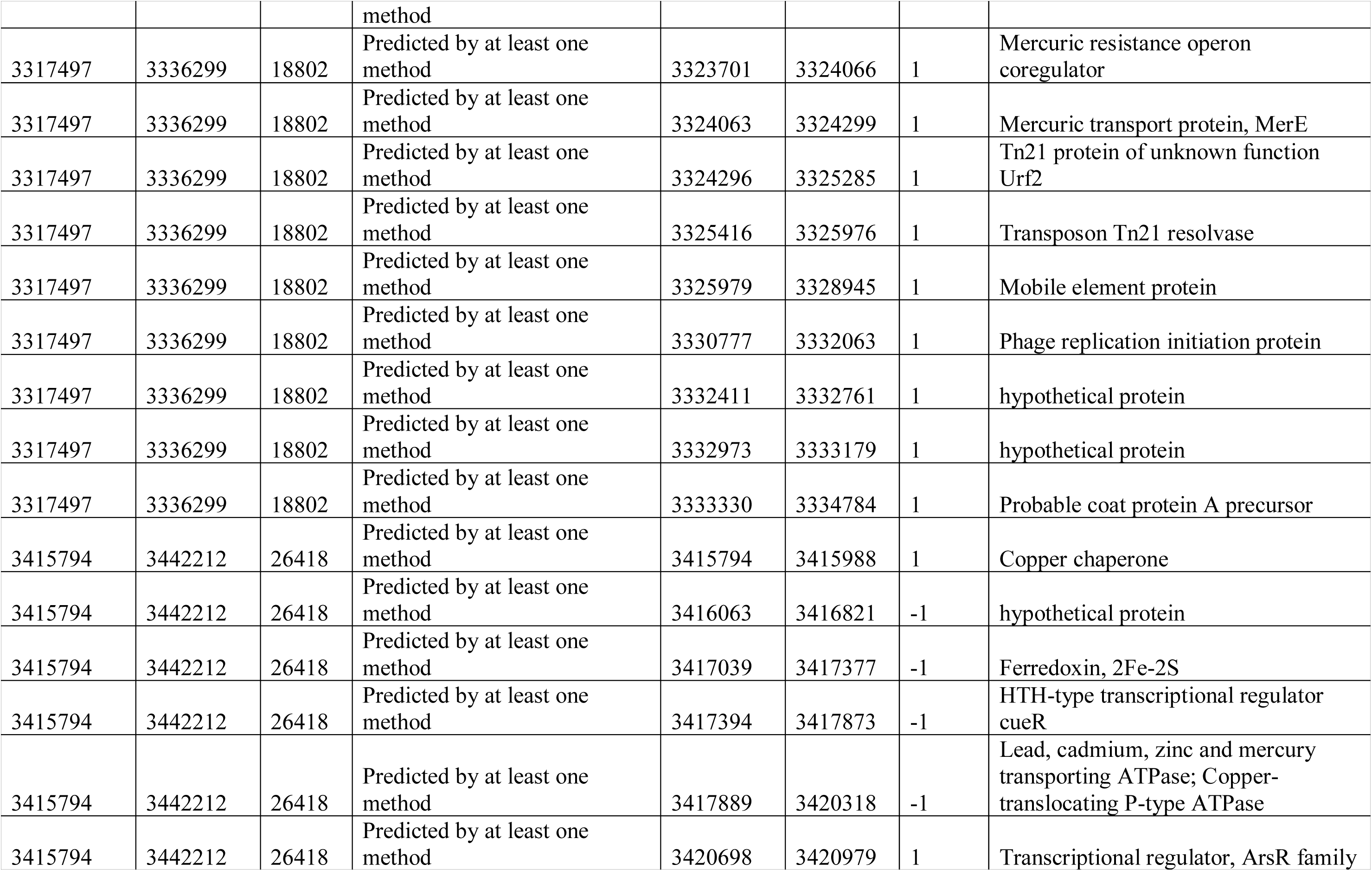
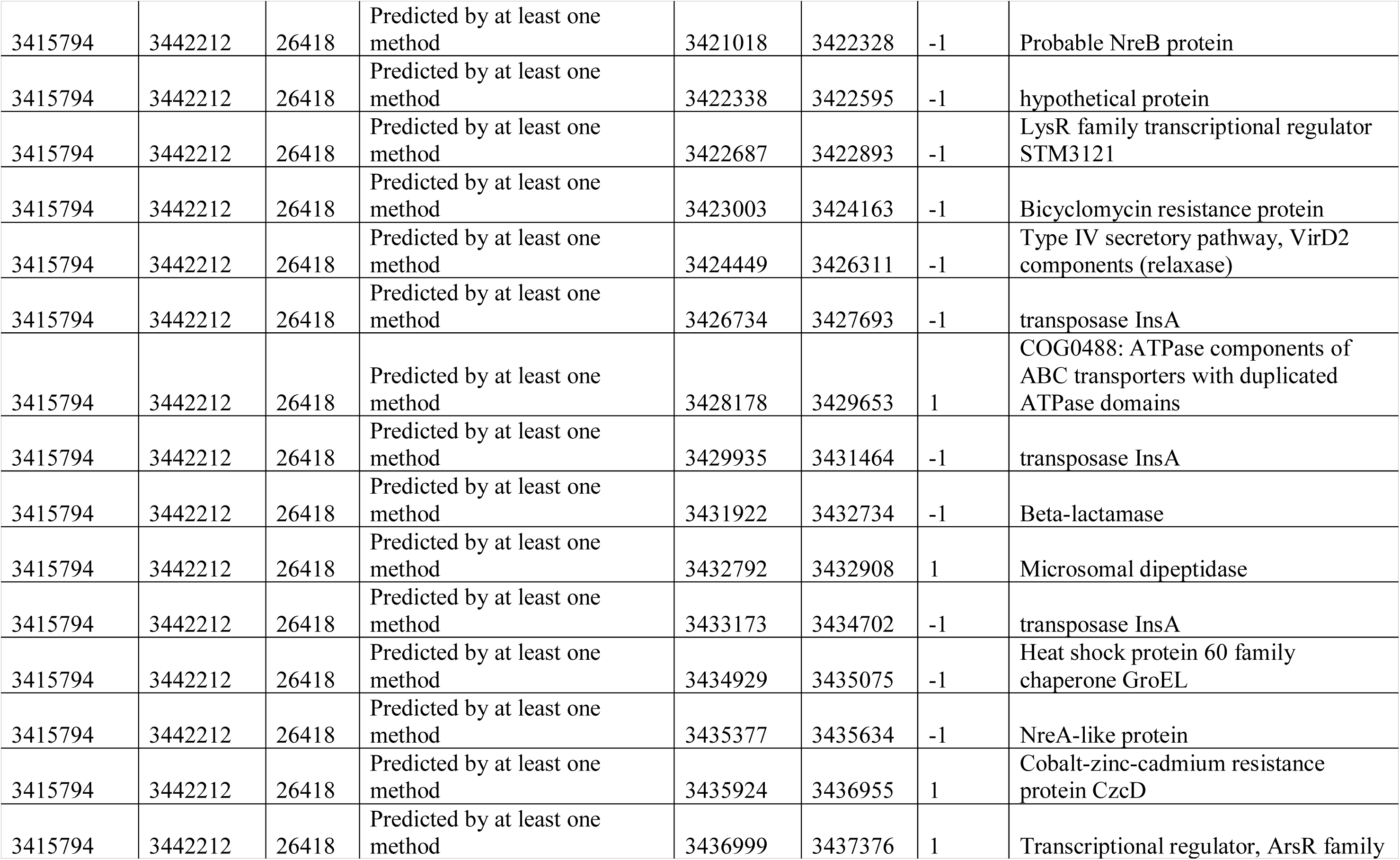
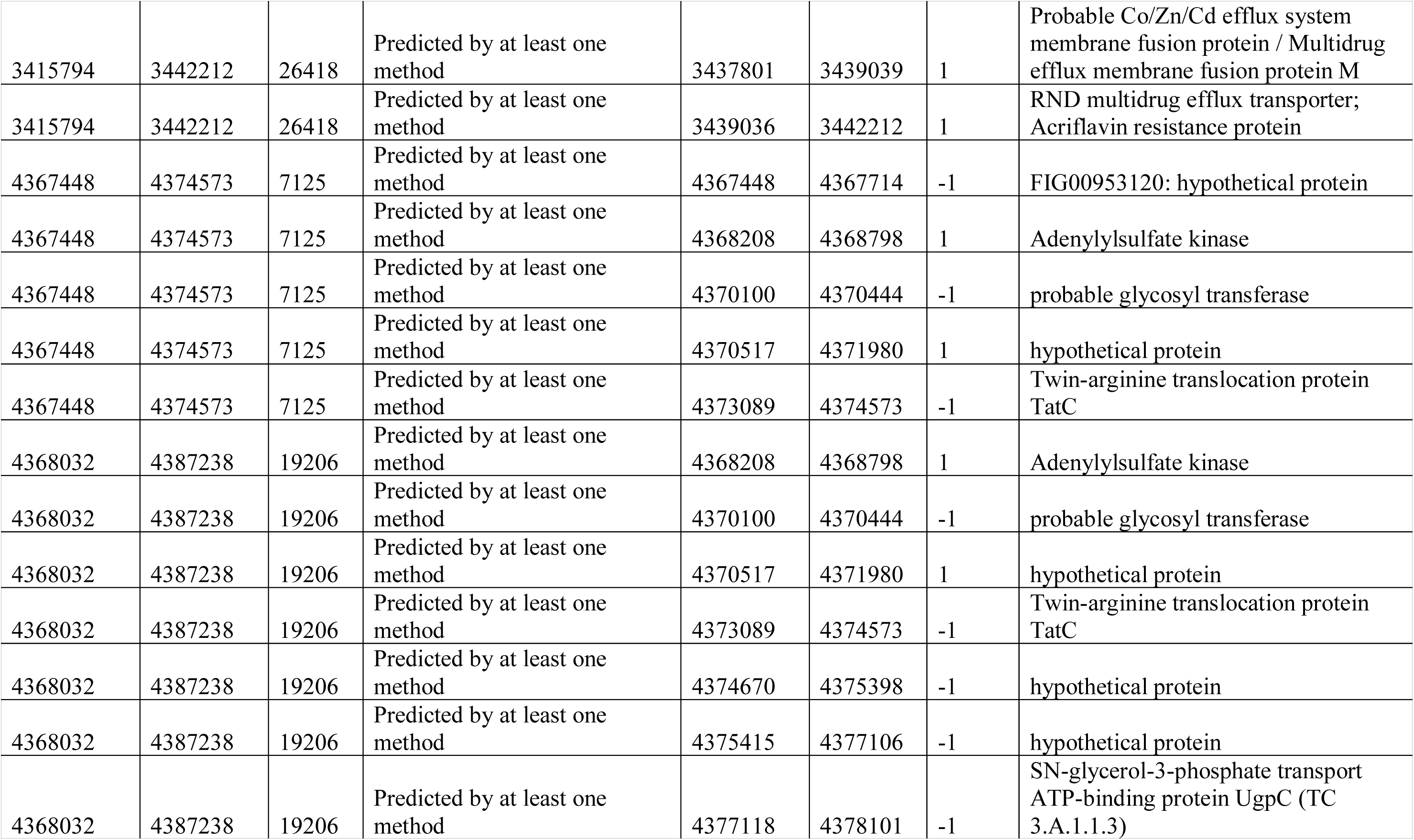
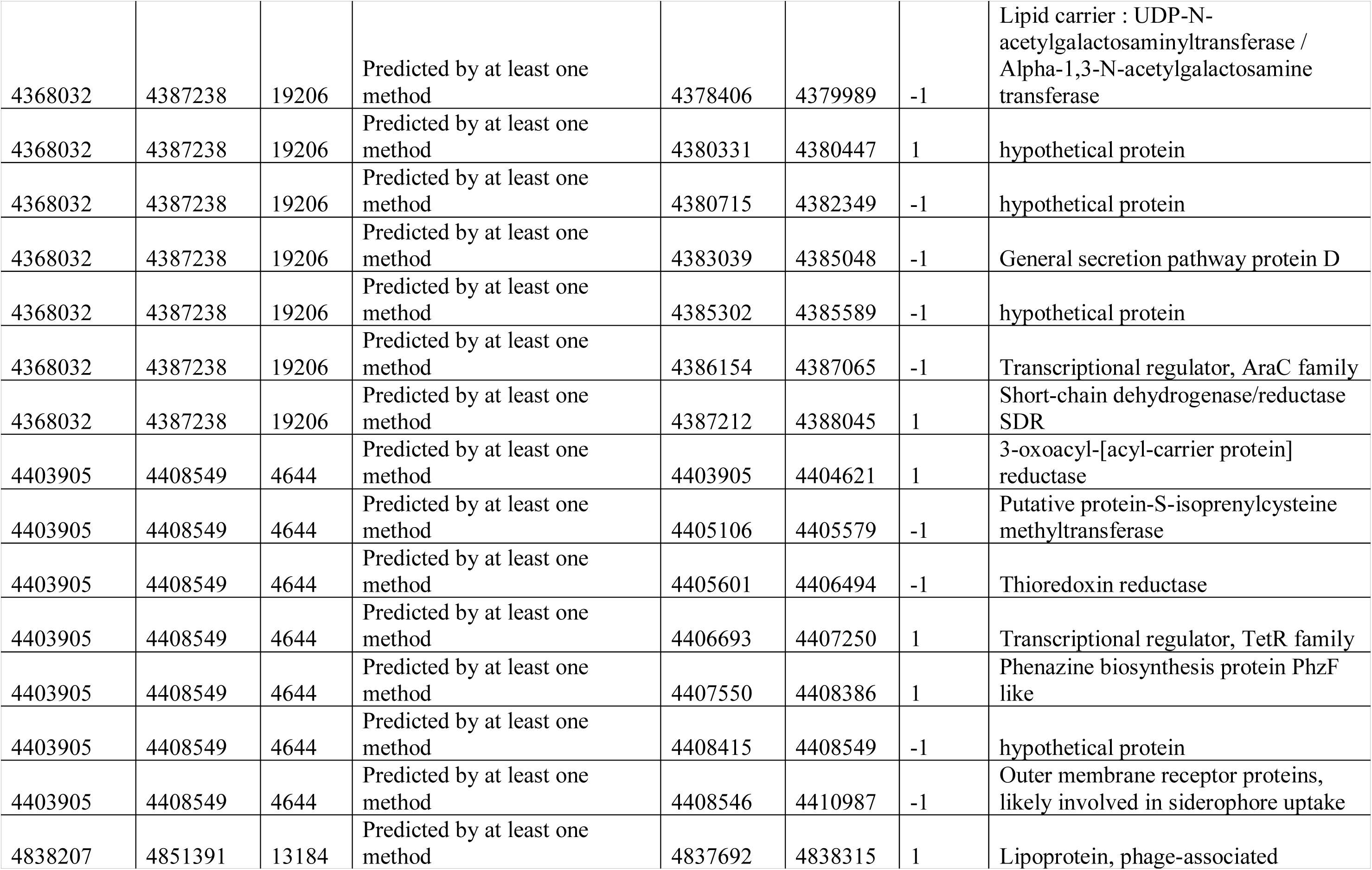
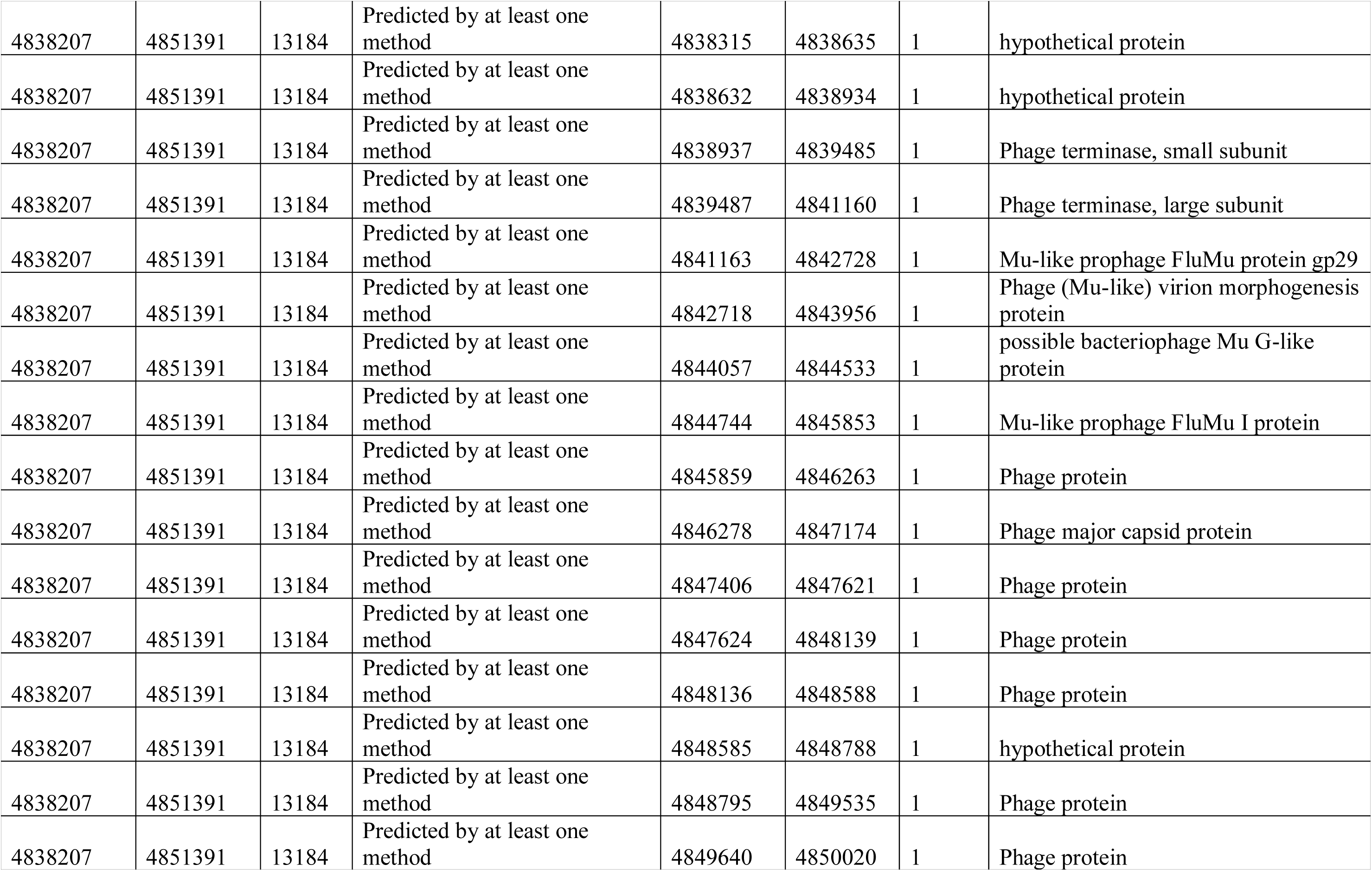
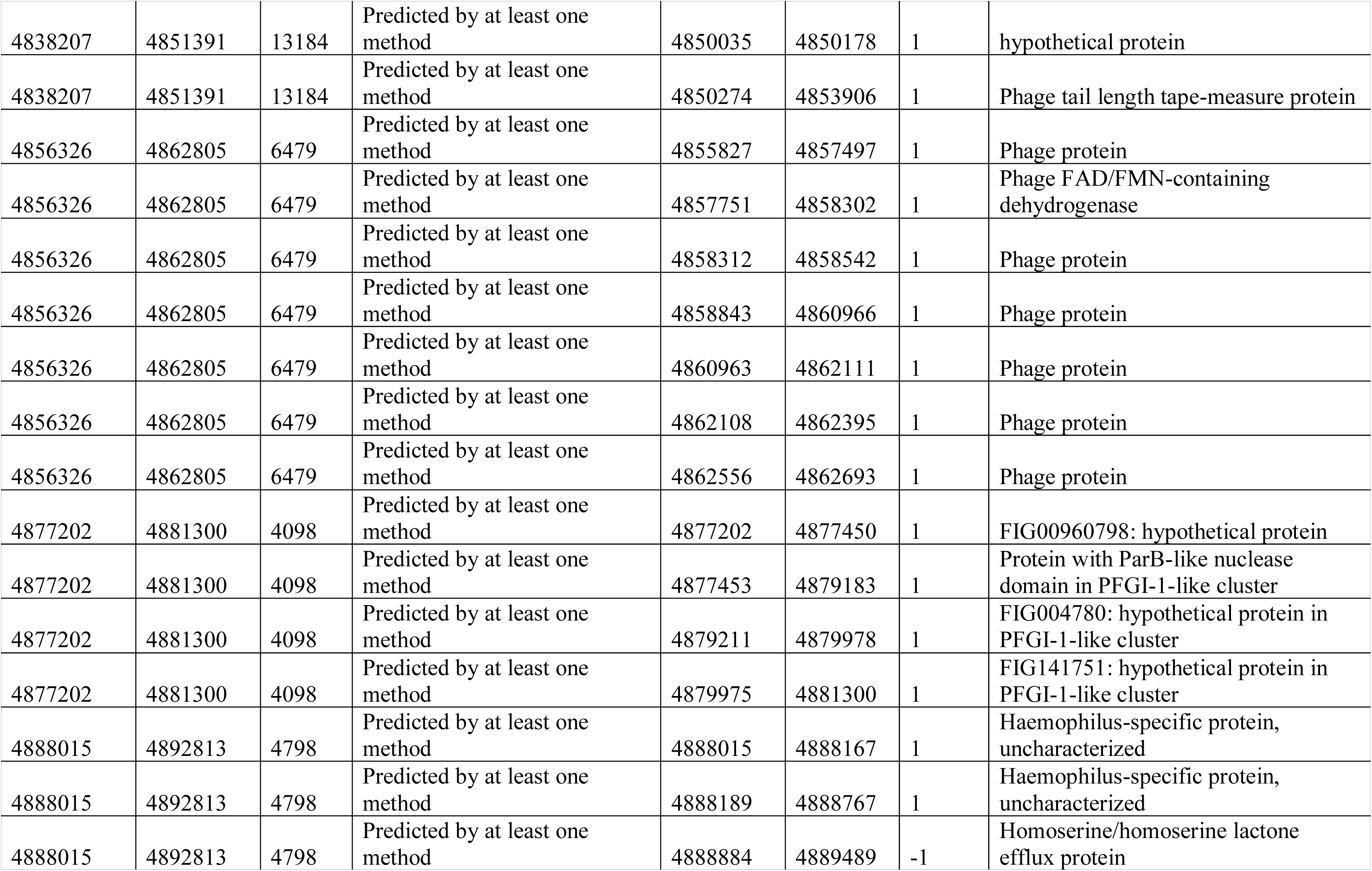
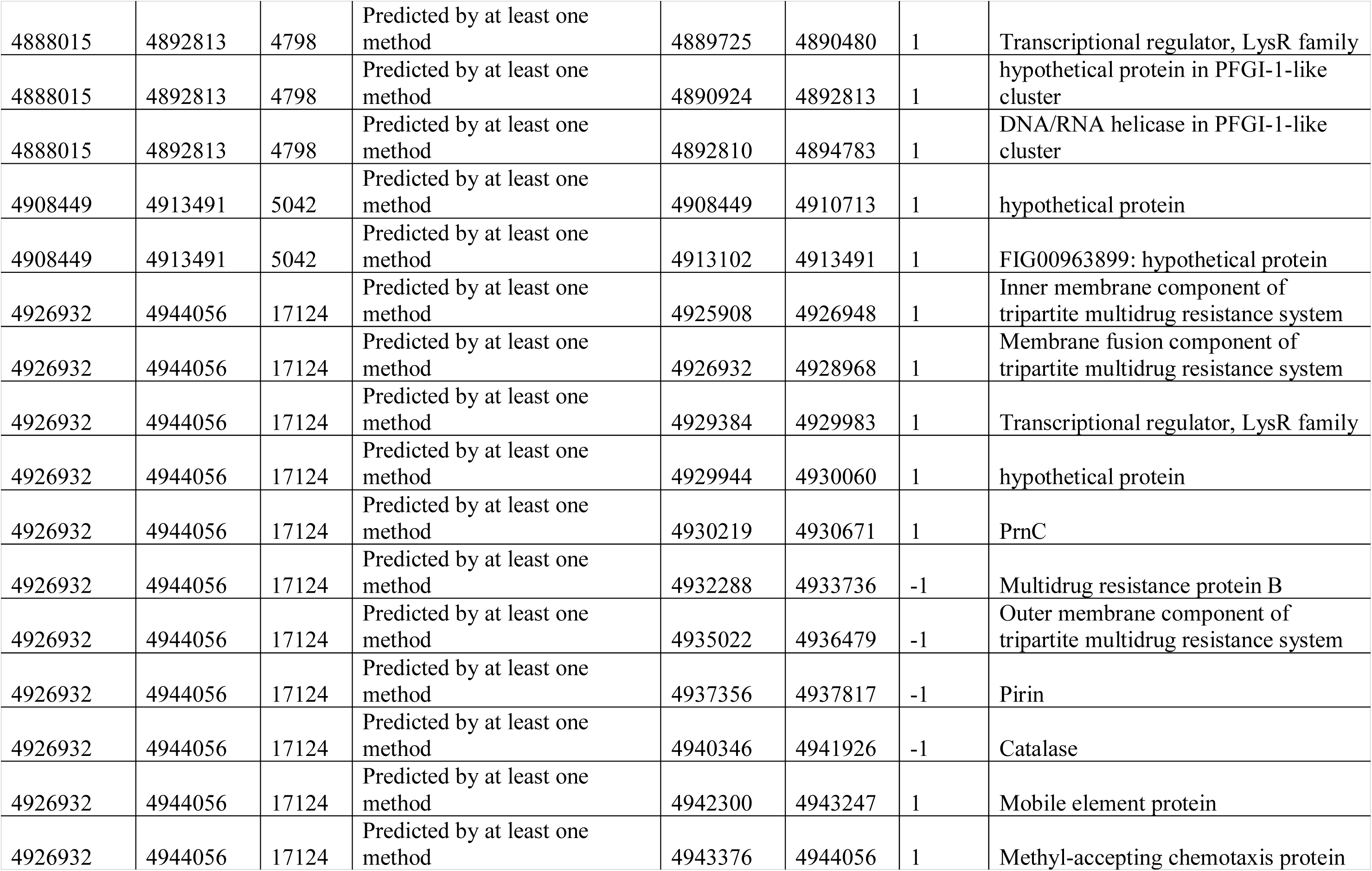
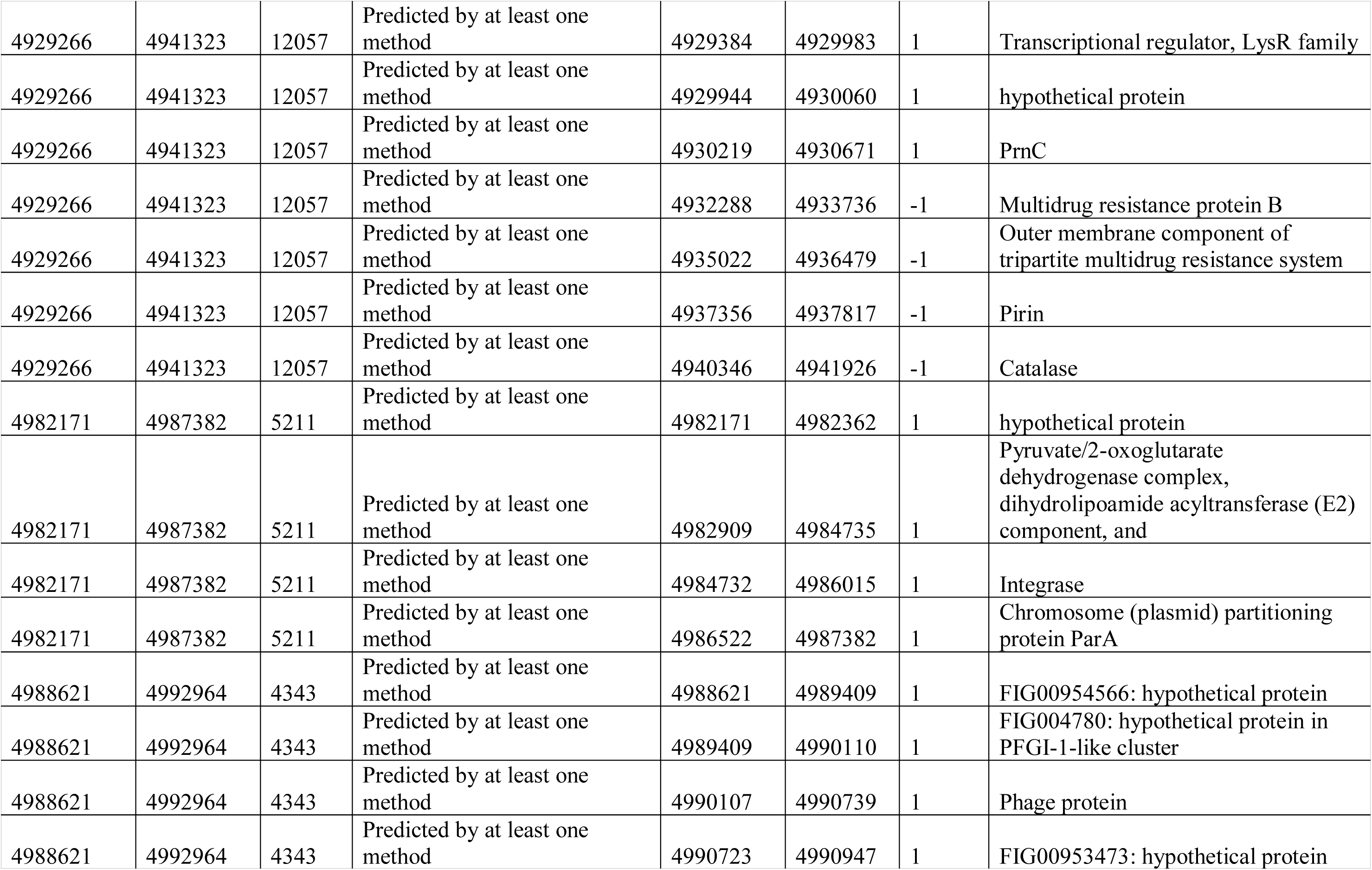
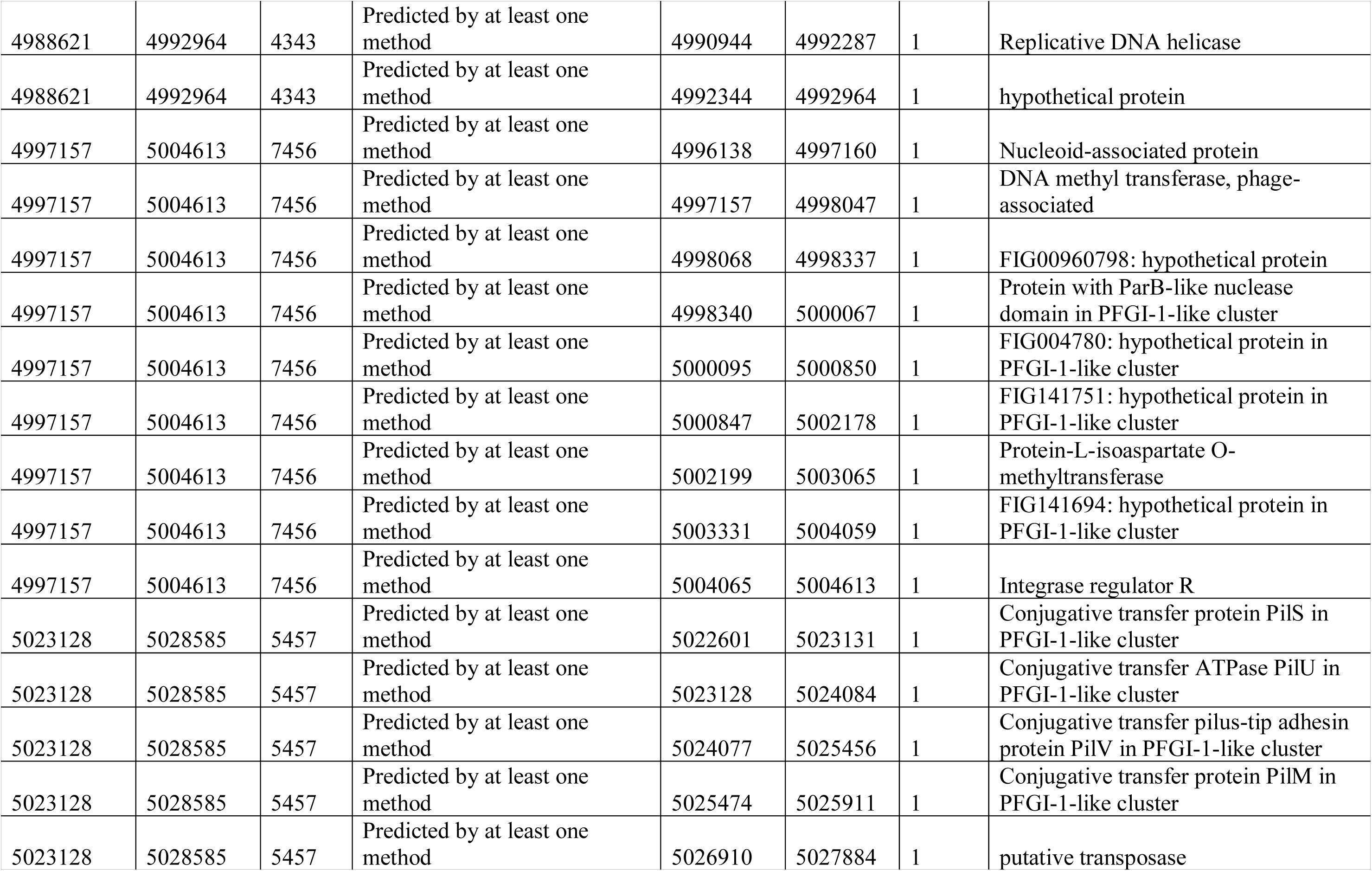
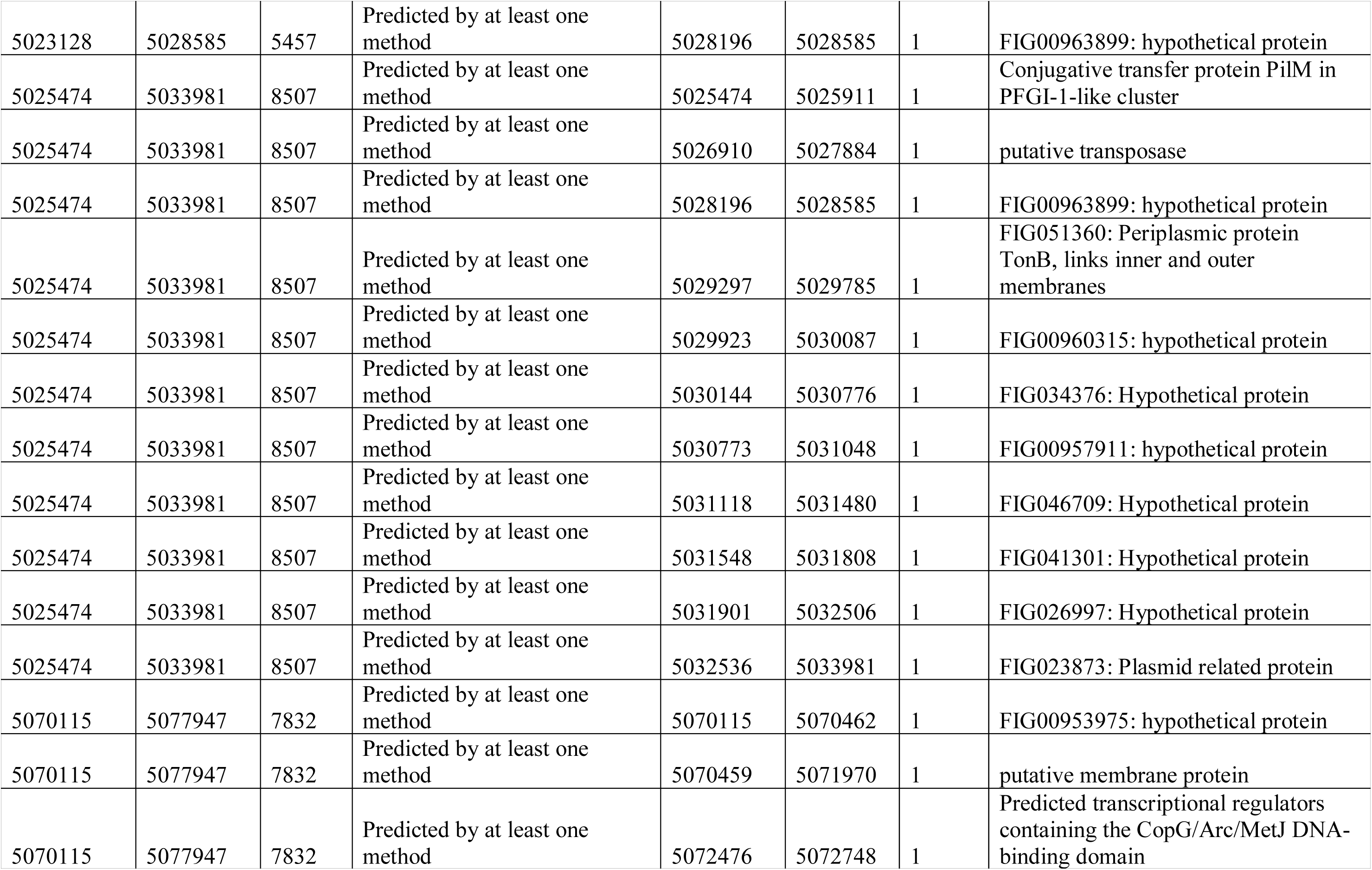
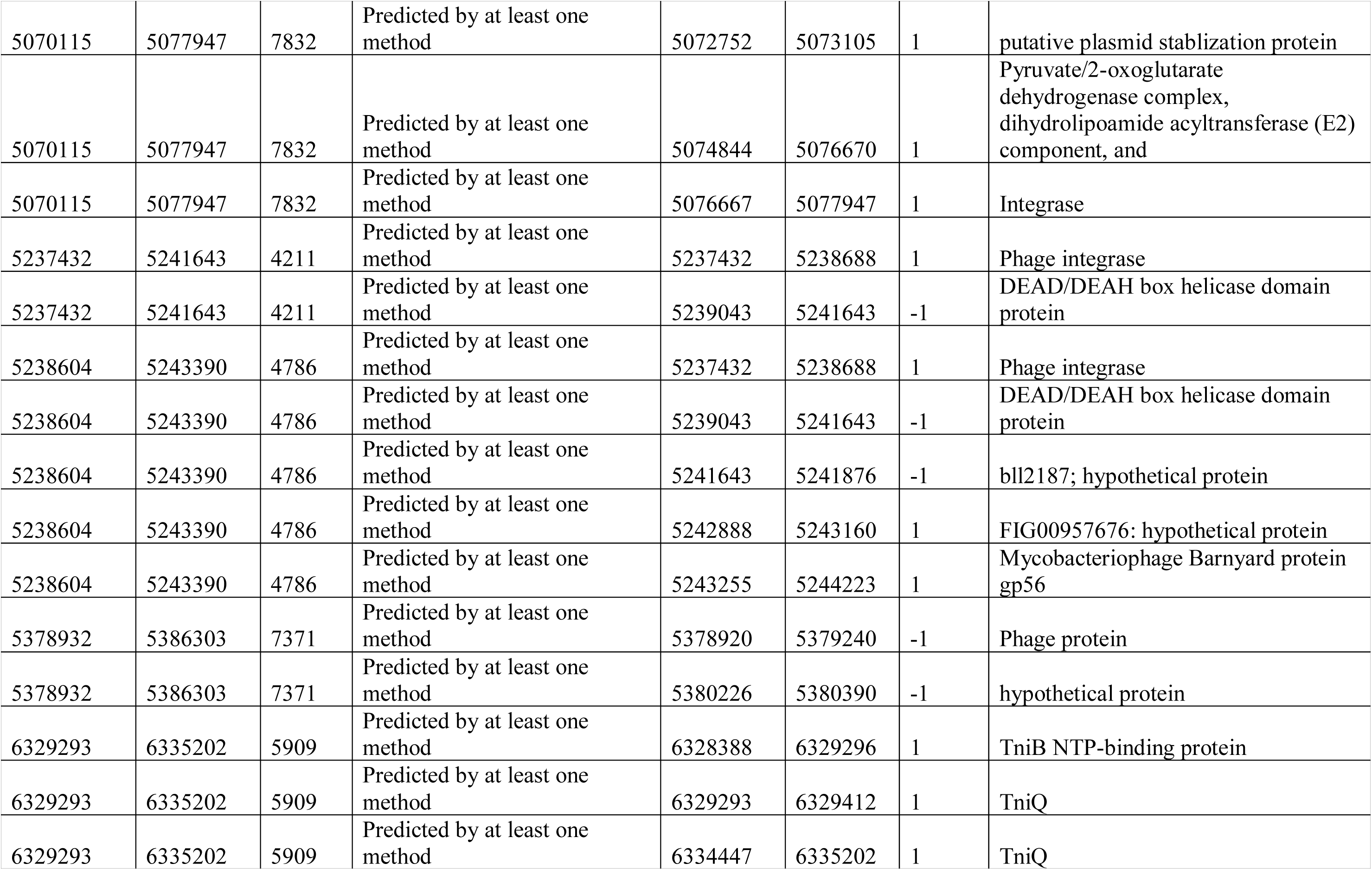
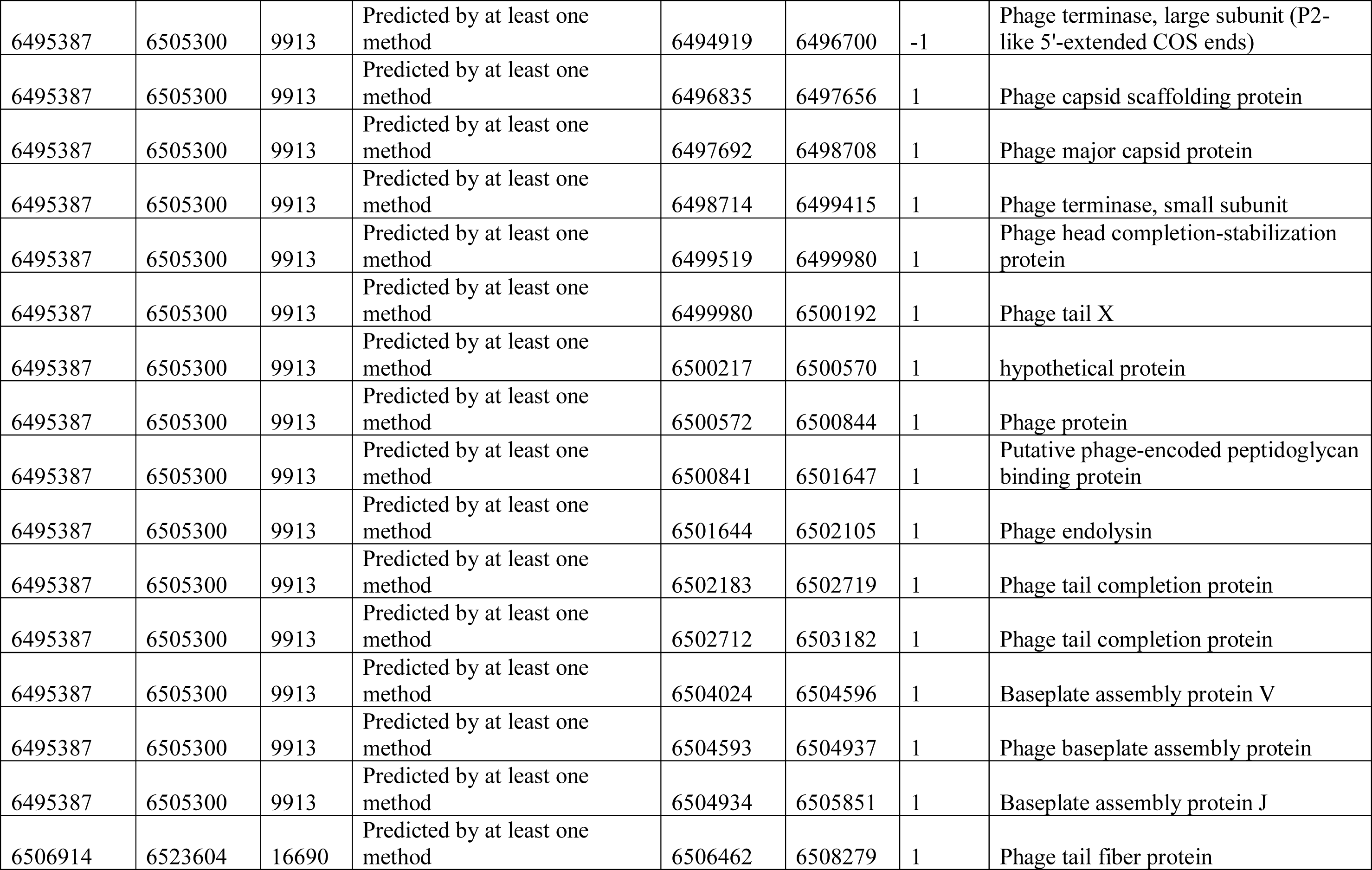
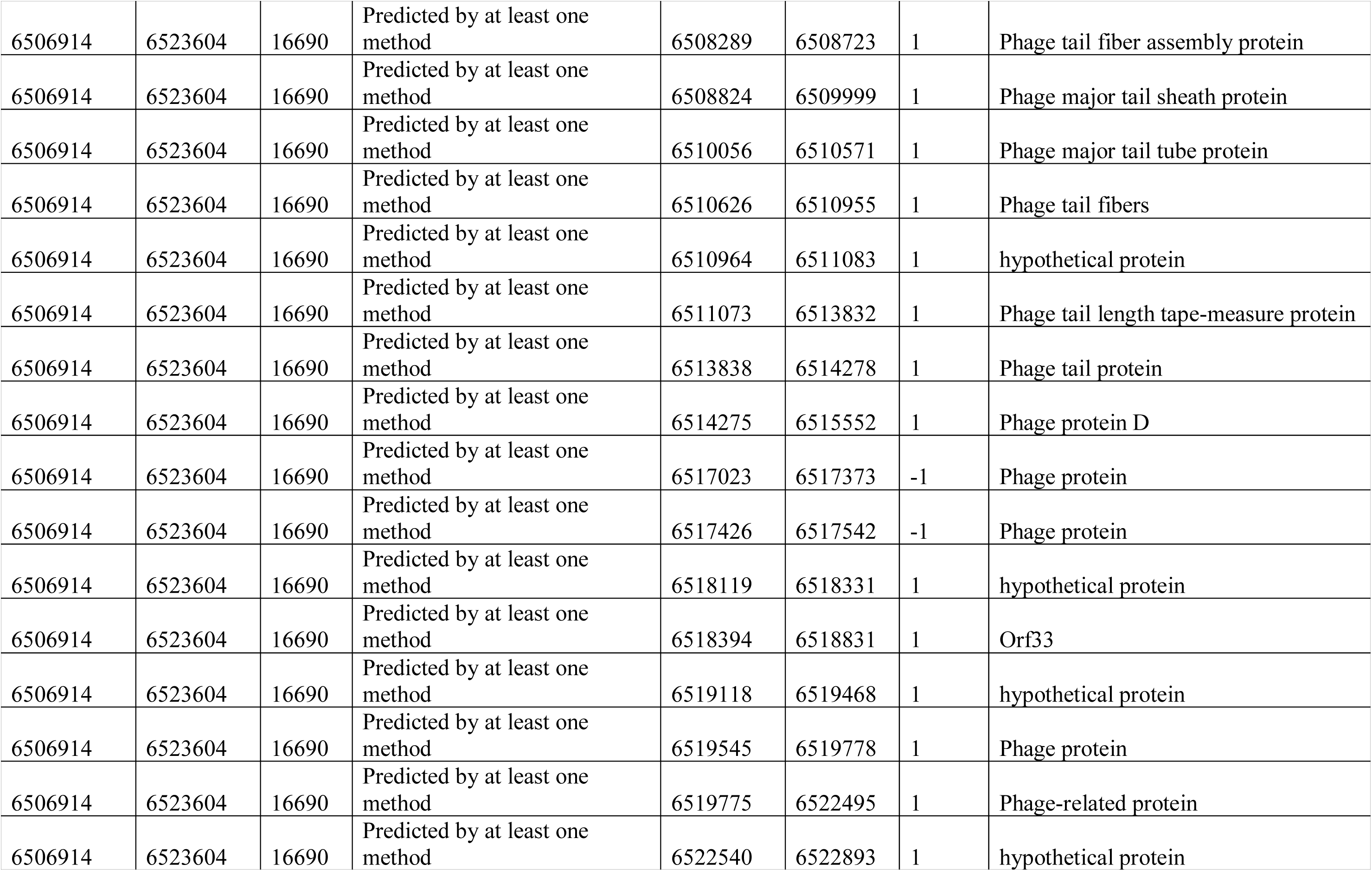
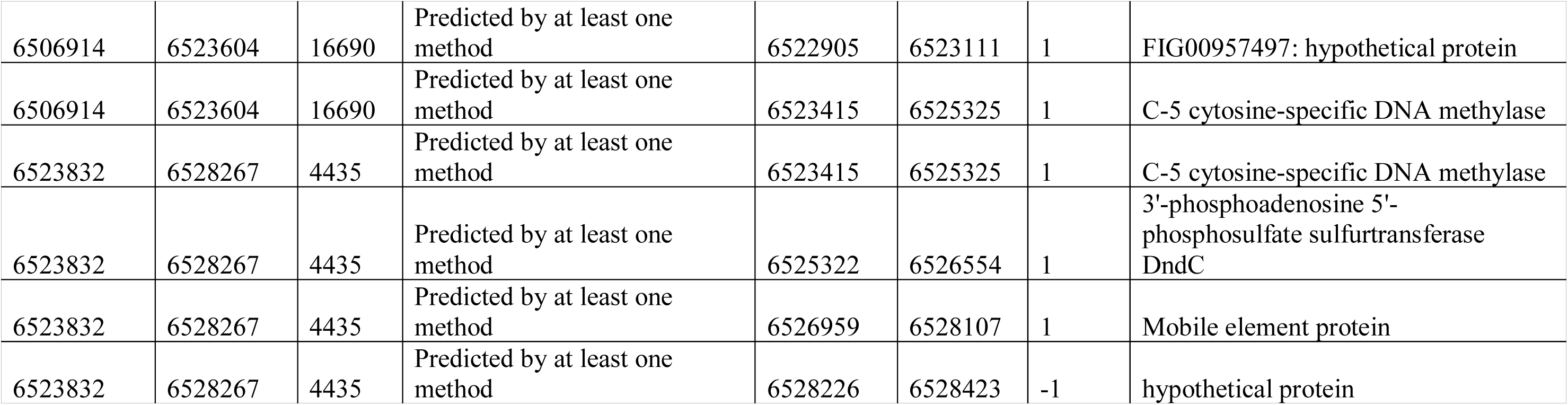
Predicted genomic islands from the full-length genome of PASGNDM345.

**Table S5:**
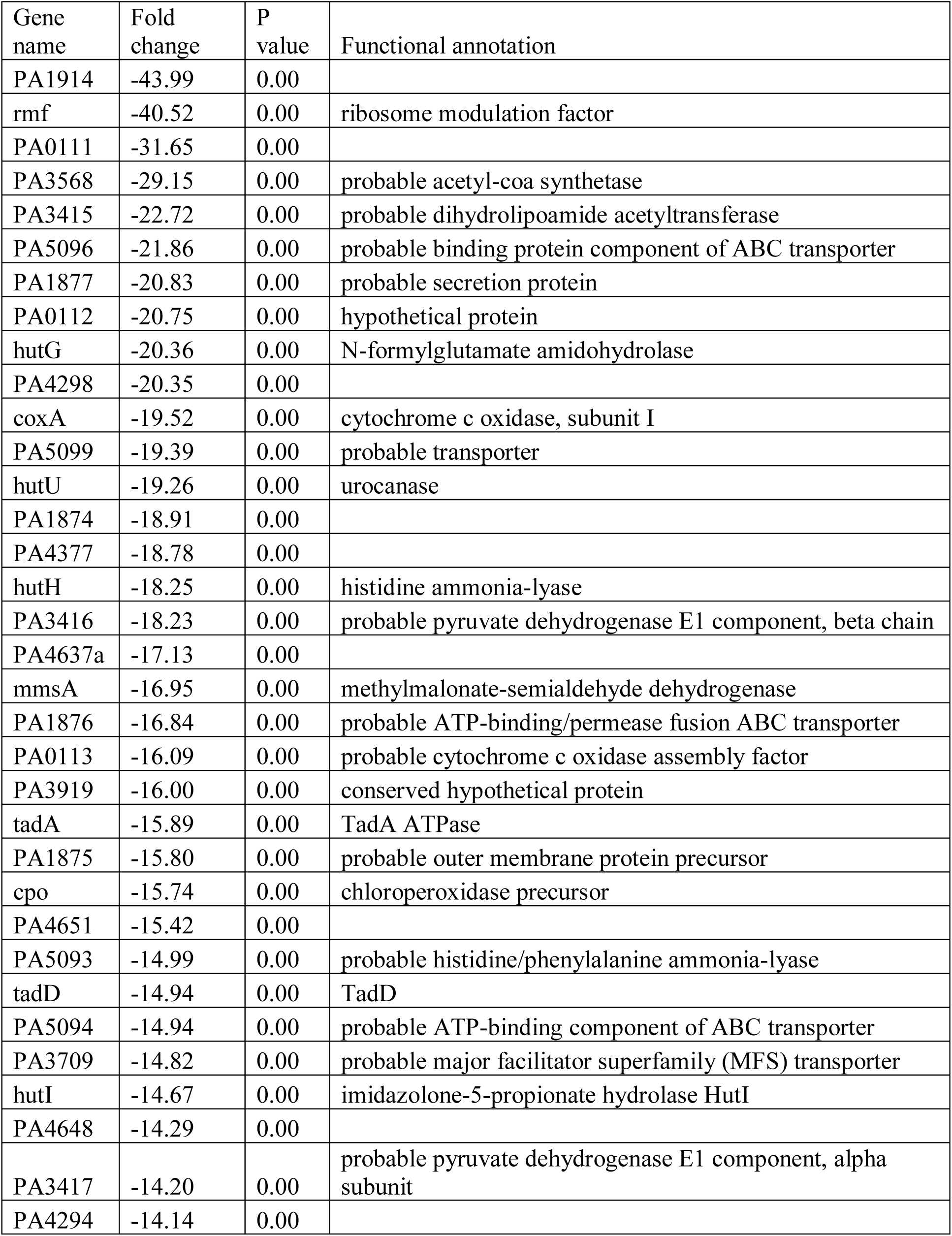
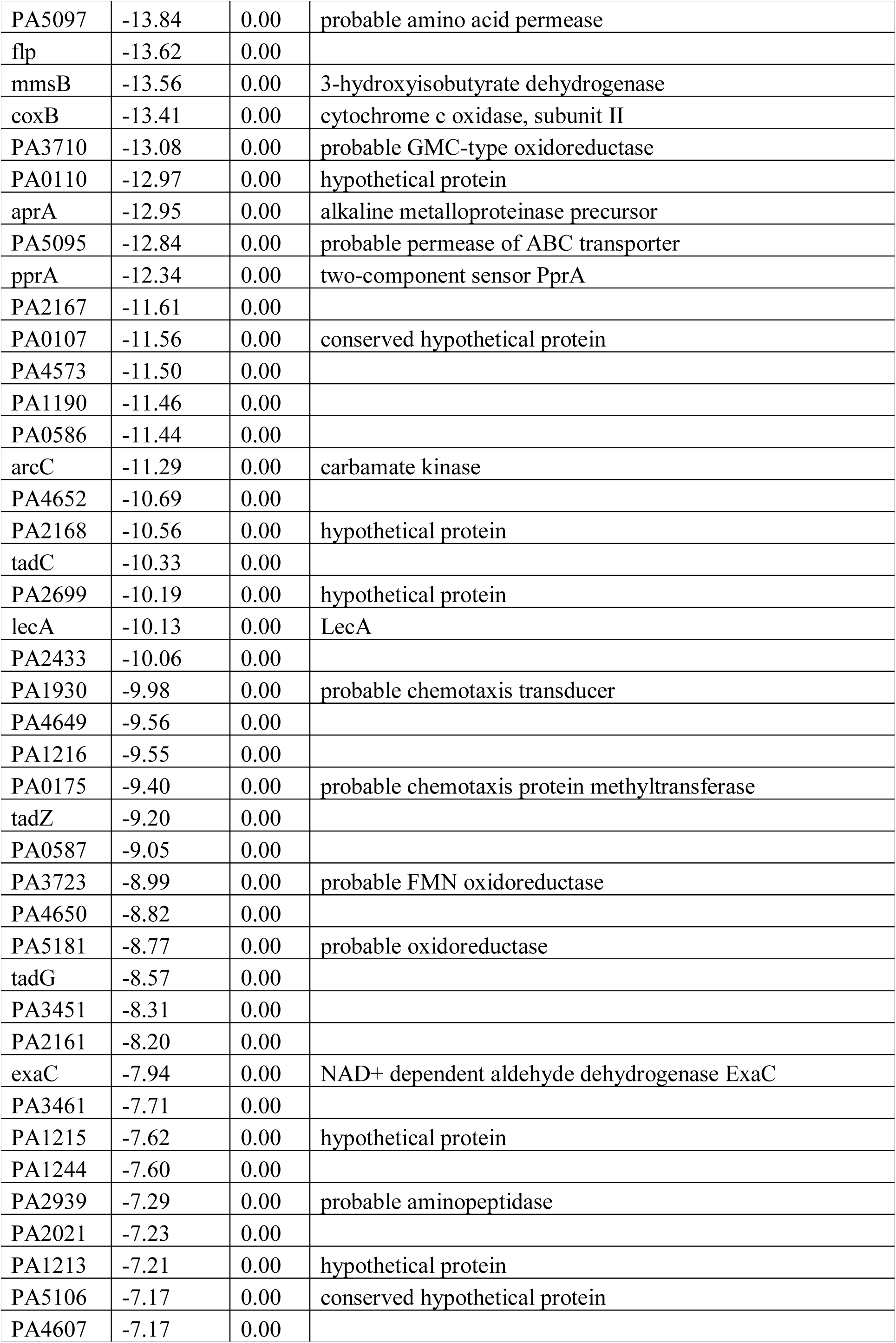
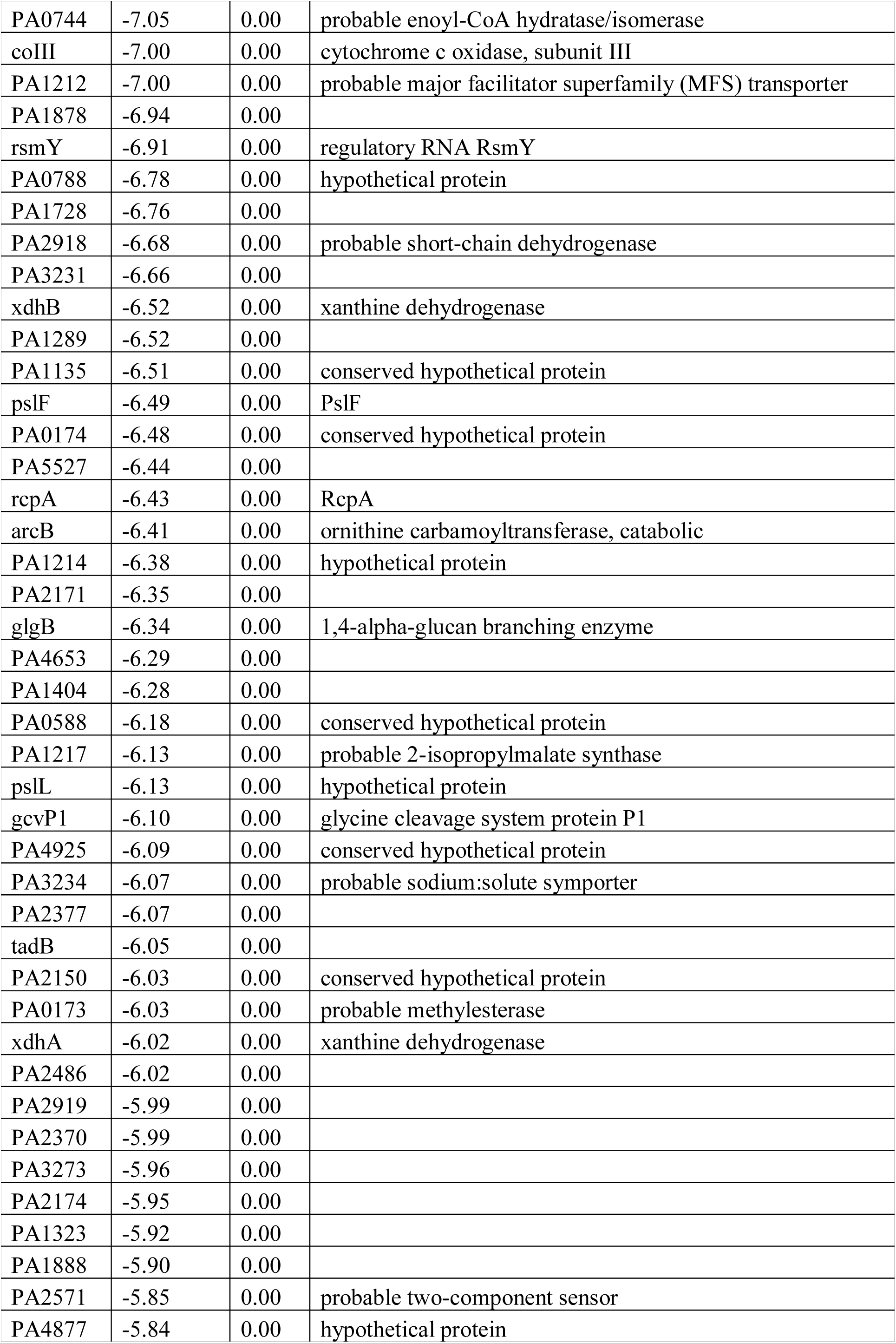
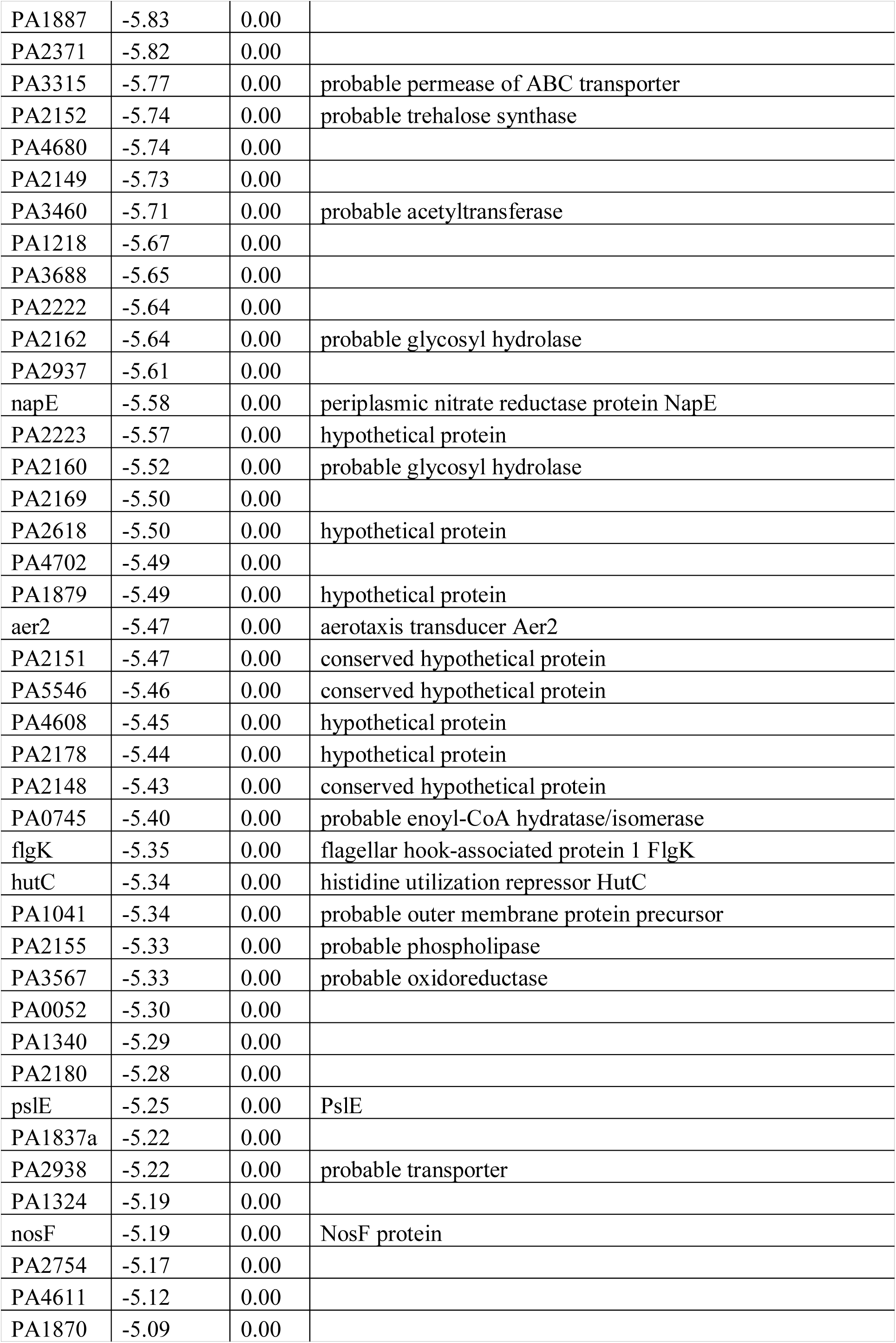
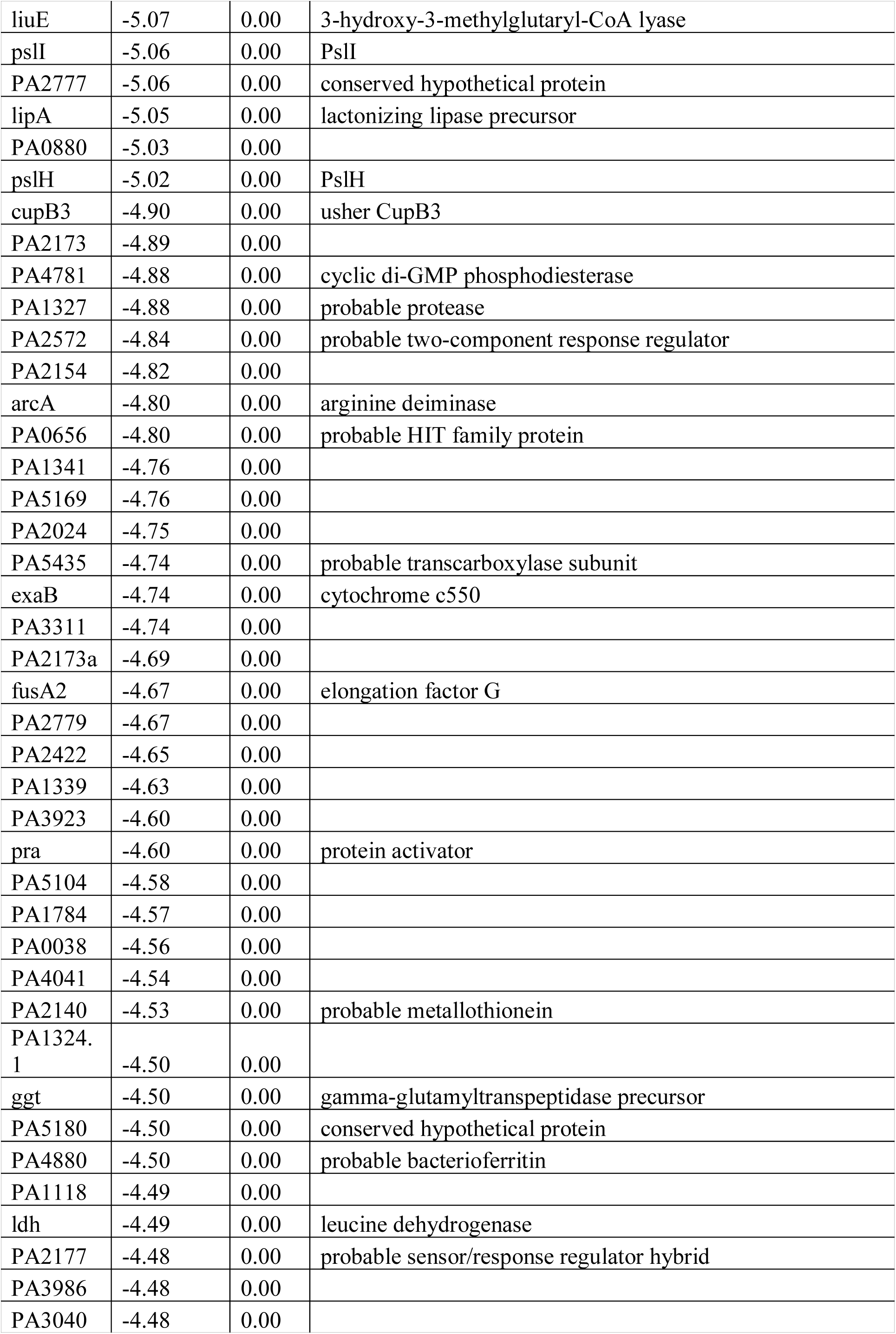
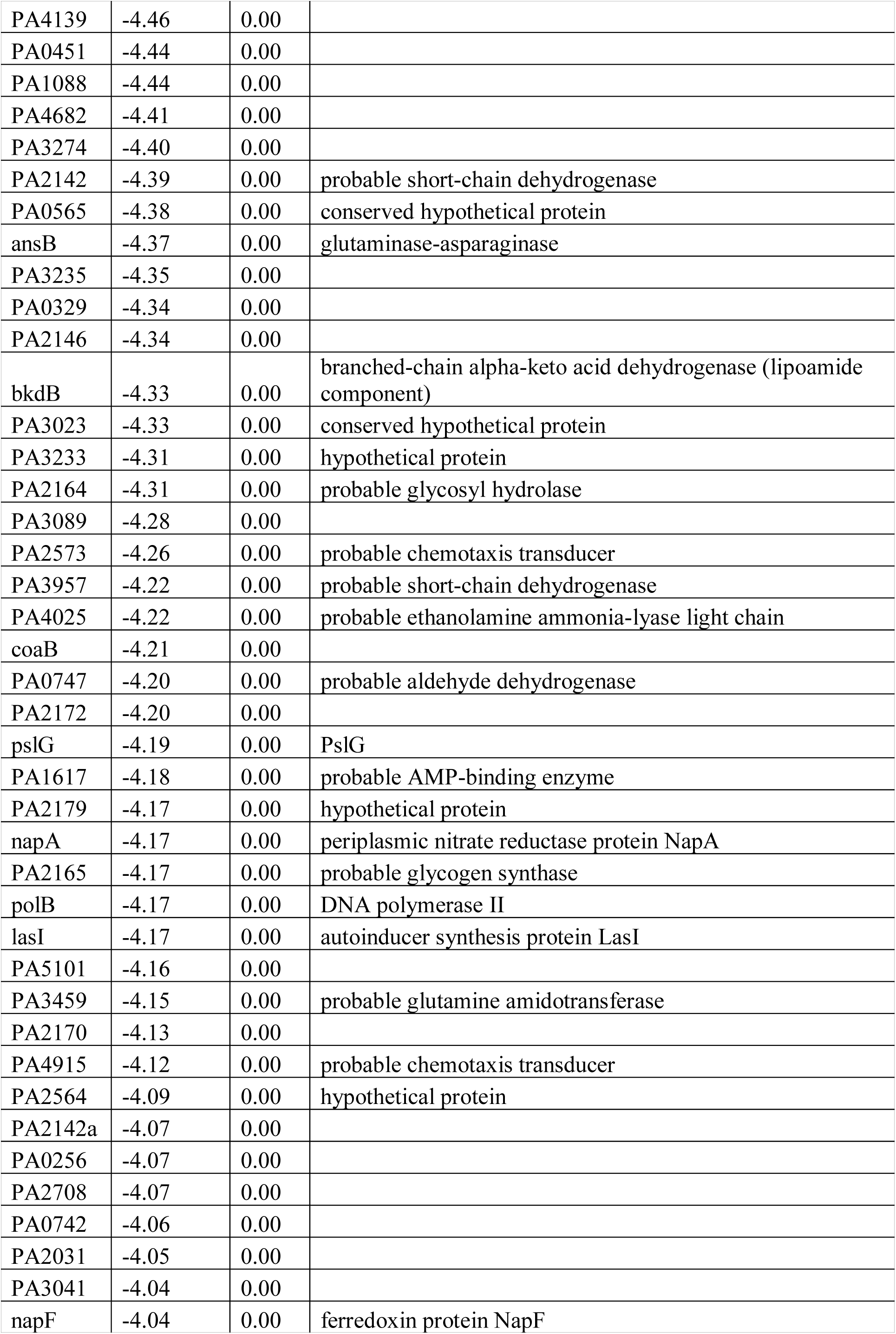
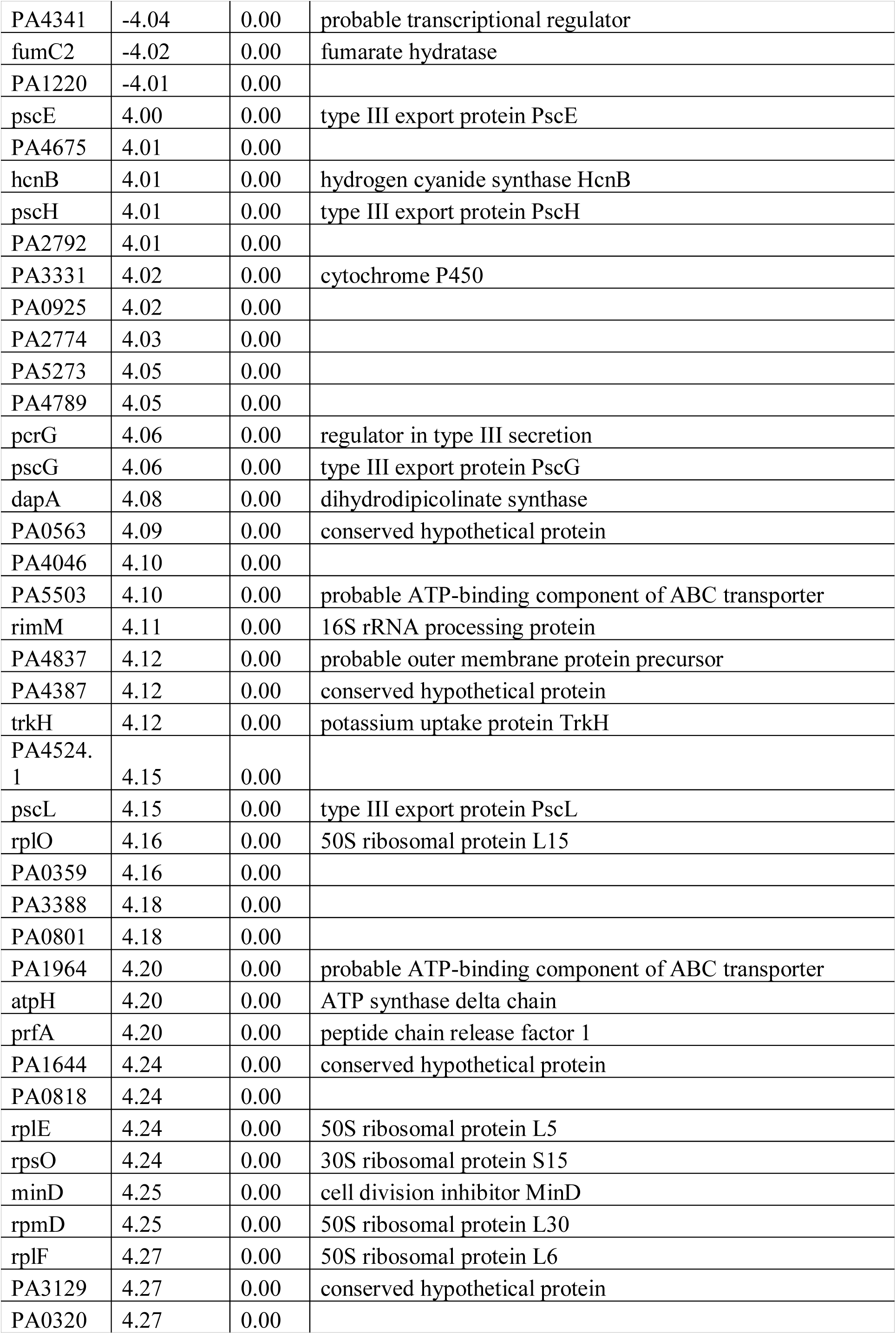
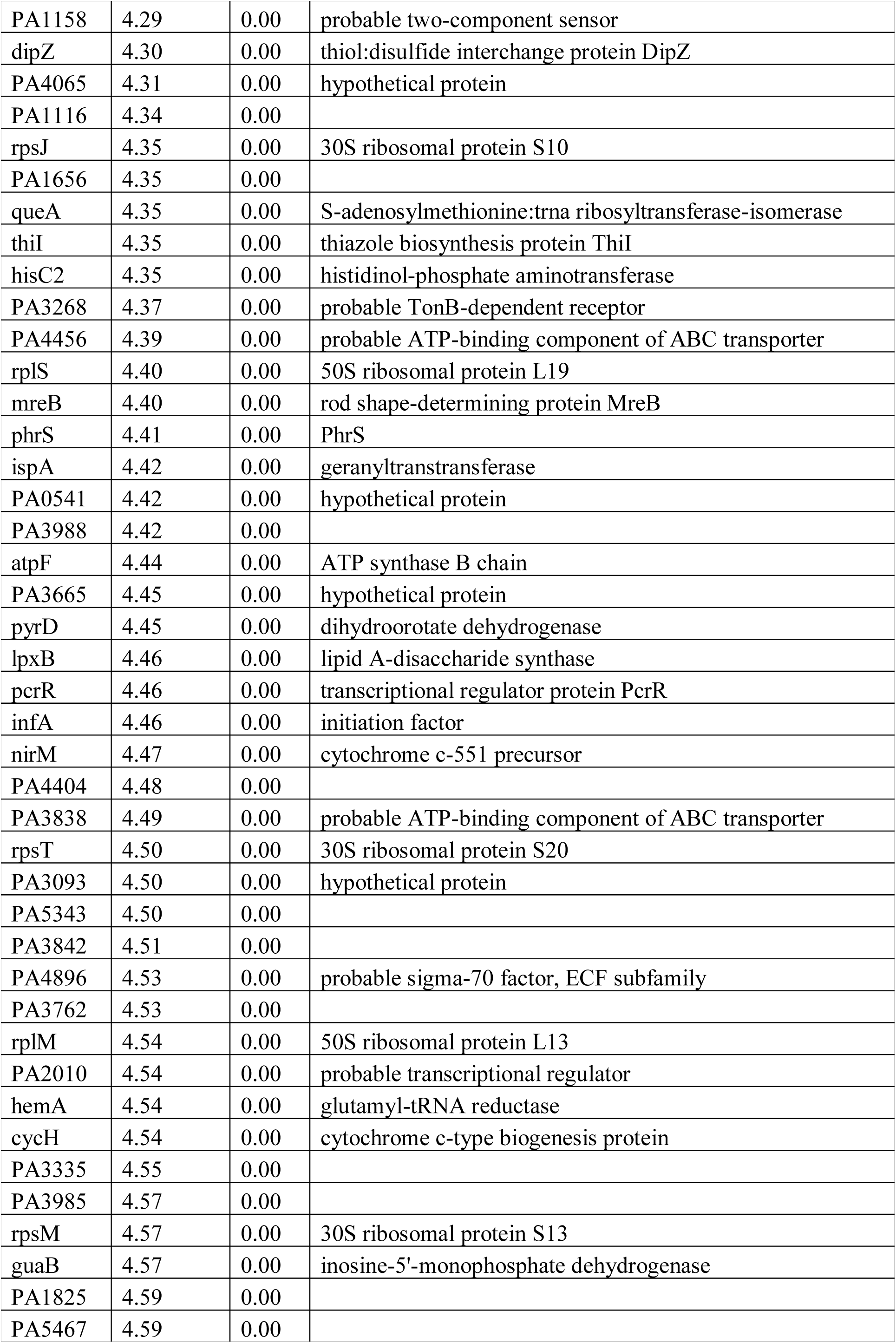
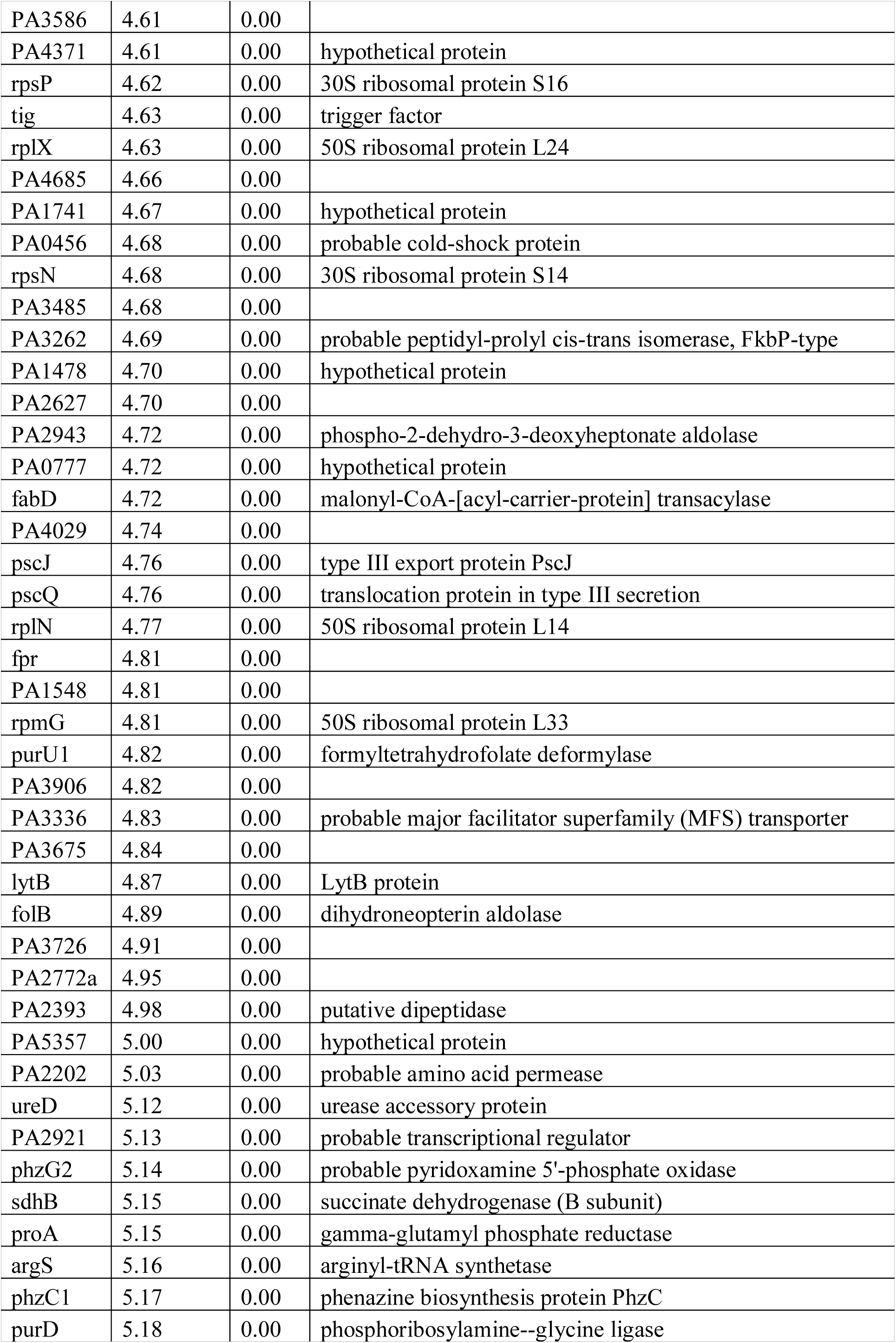
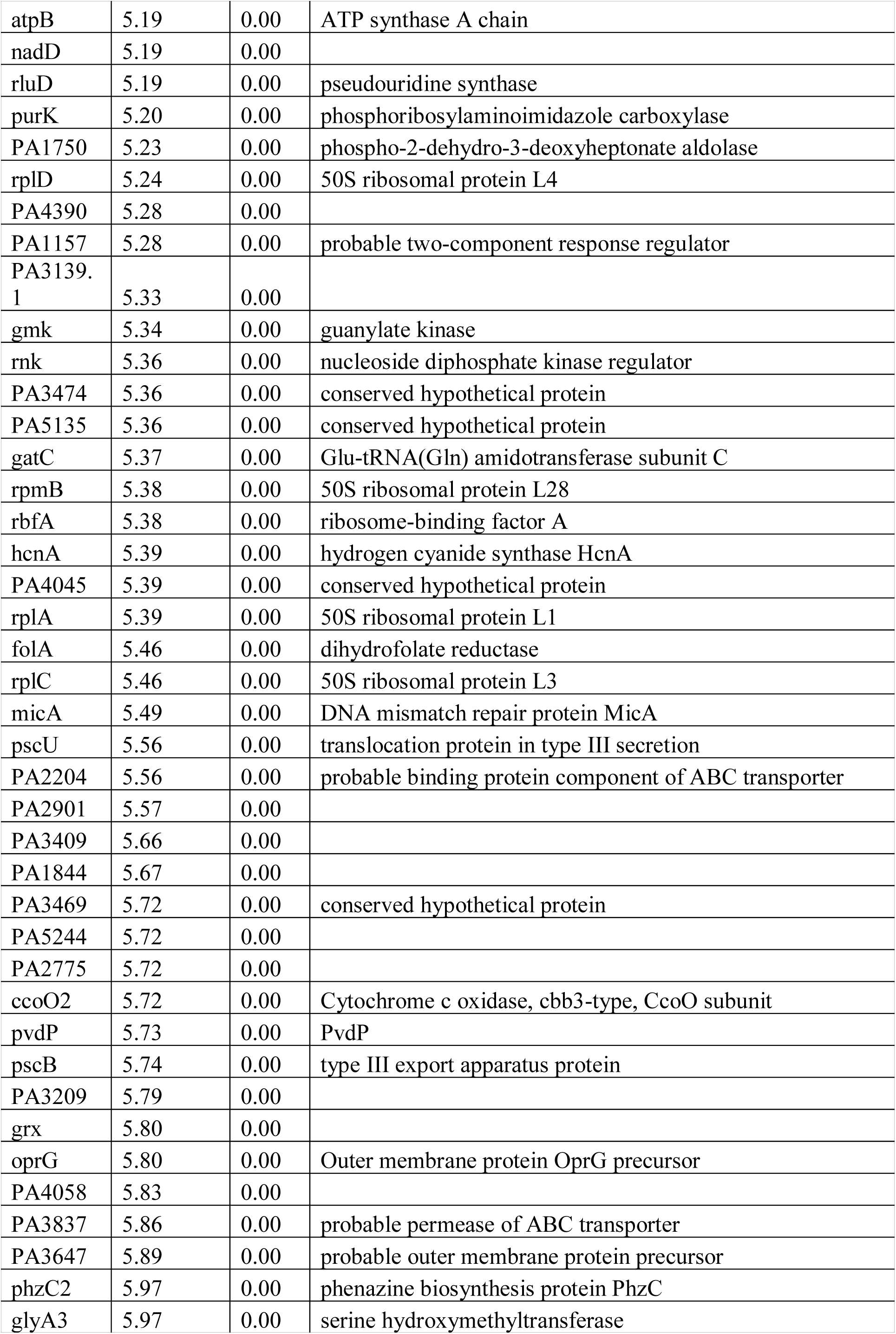
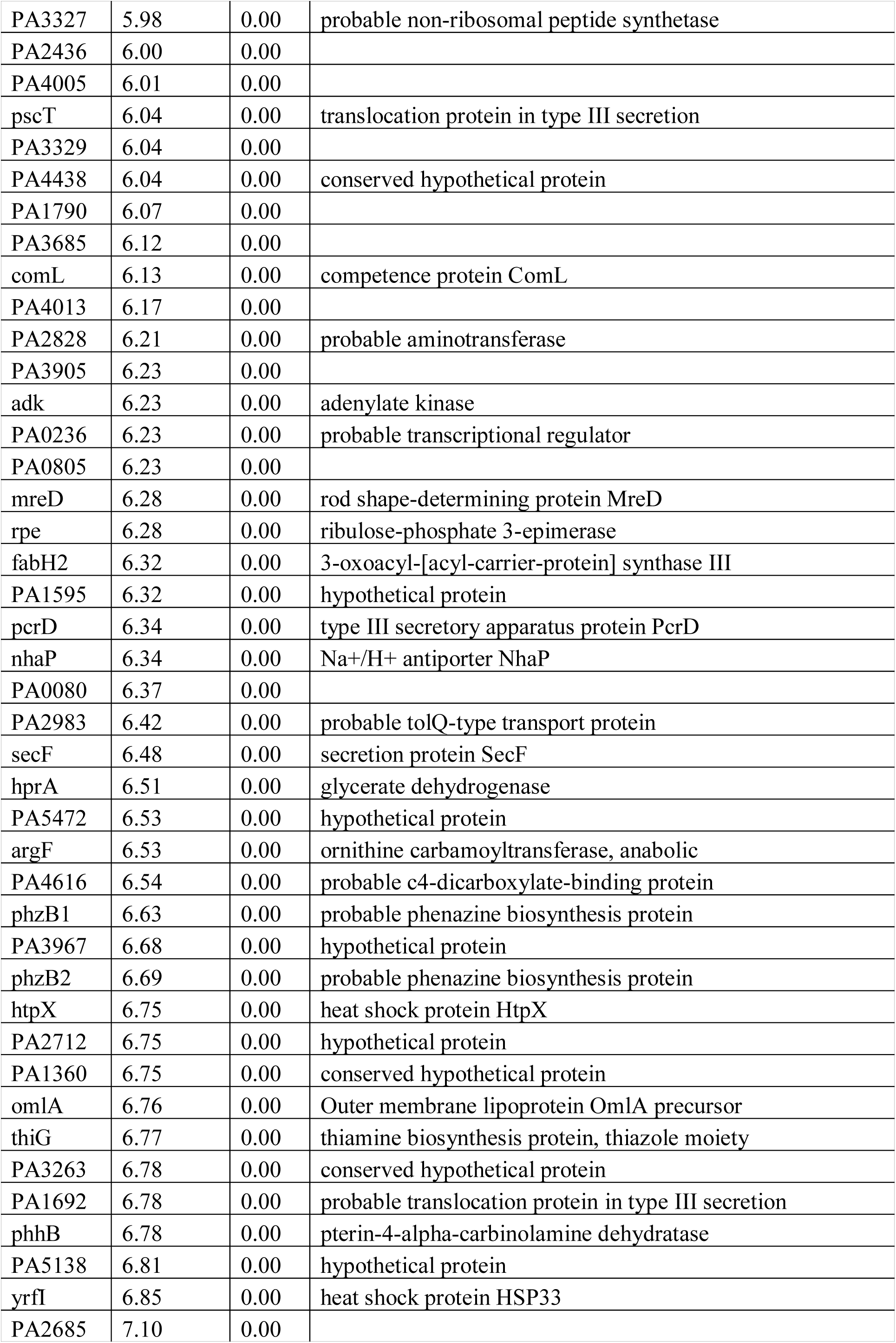
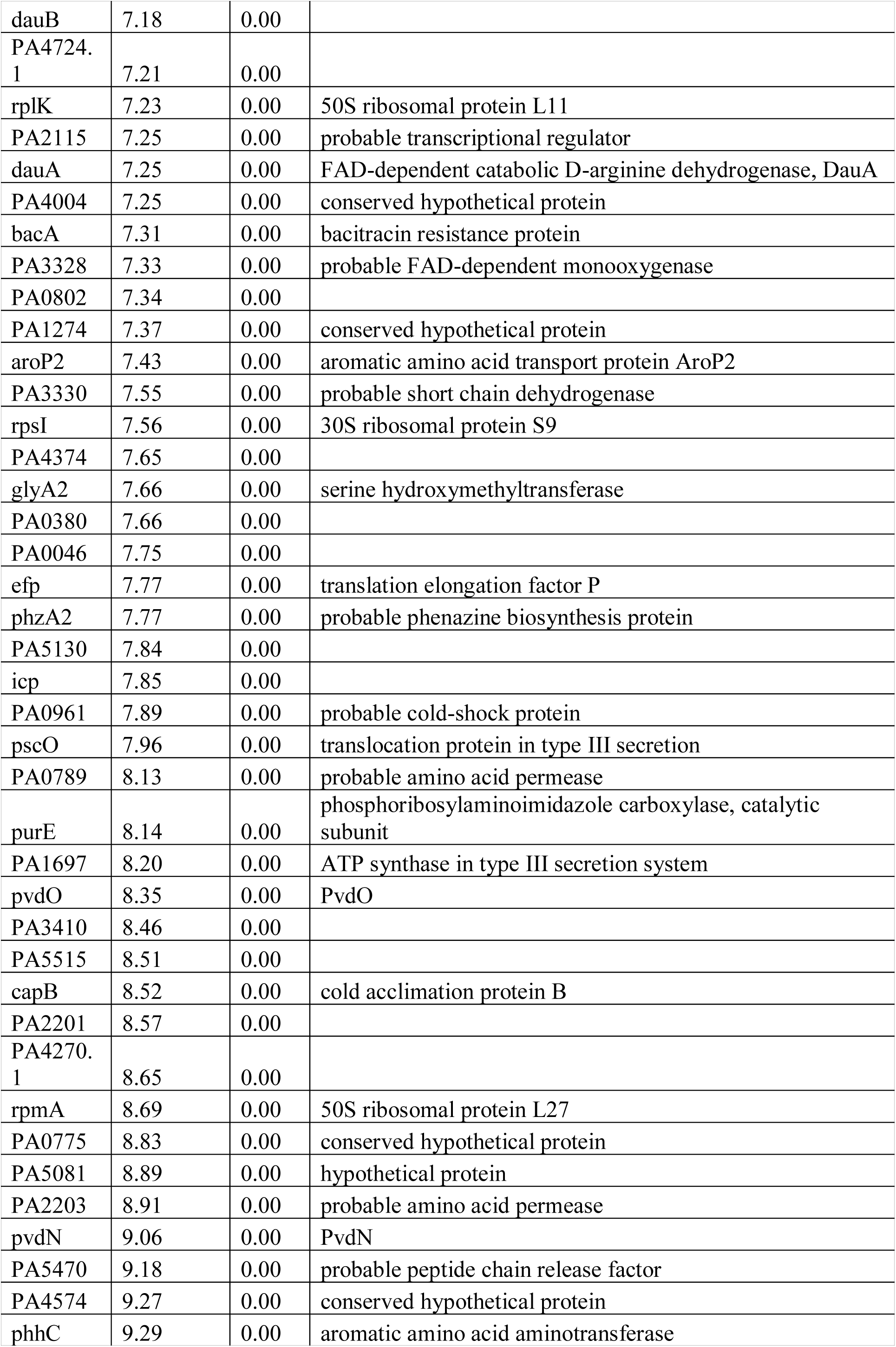
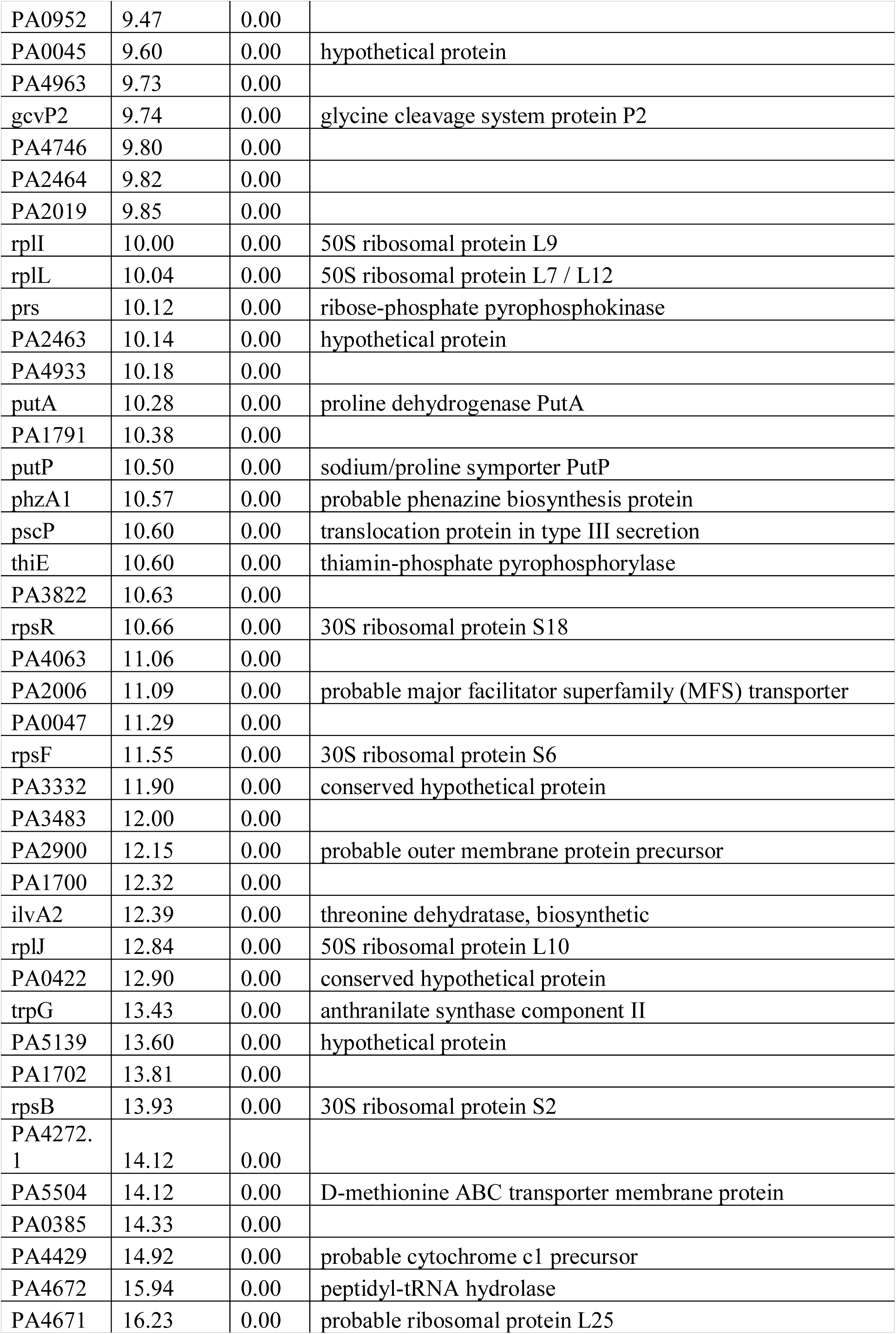
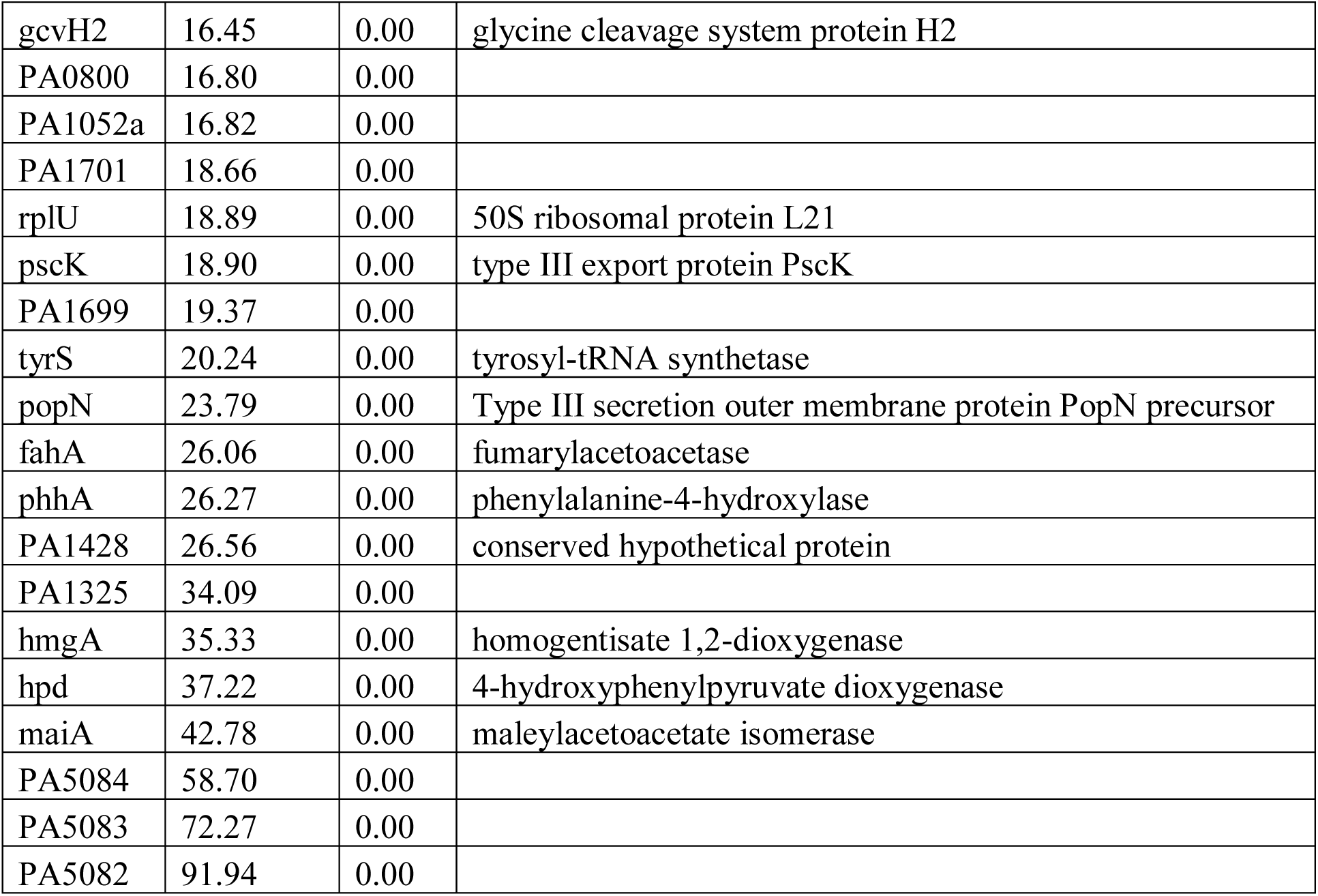
The 550 genes that are differentially expressed between AZM-treated and non-treated PAO1. Gene names, fold change, P value and functional annotations are shown in the table. Cut-off: fold-change > 4, P value < 0.05.

**Table S6:**
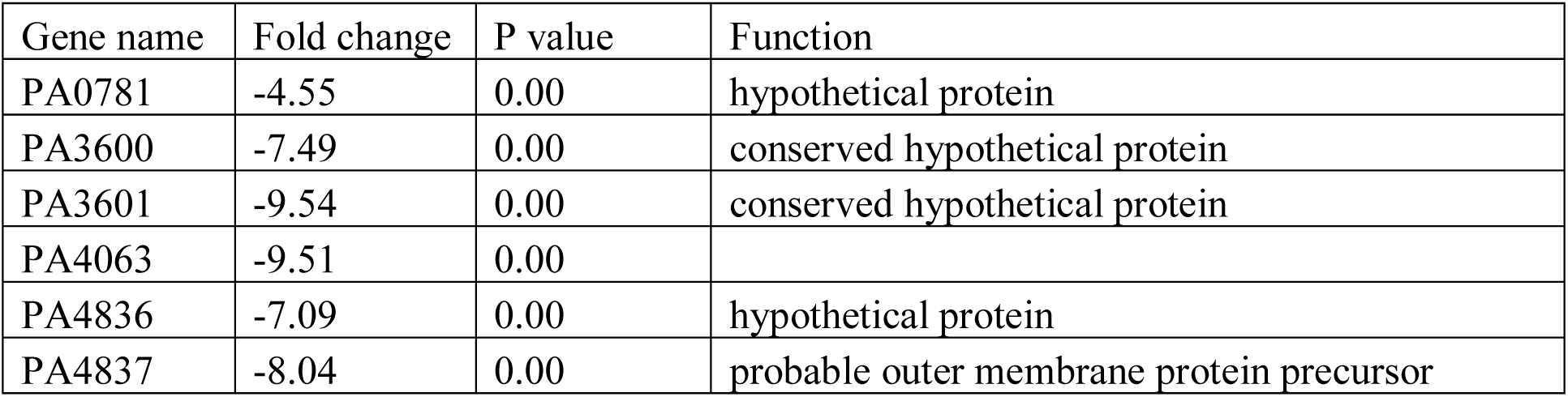
The 6 genes that are differentially expressed between AZM-treated and non-treated PAO1/*pUCP18::msr(E)*. Gene names, fold change, P value and functional annotations are shown in the table. Cut-off: fold-change > 4, P value < 0.05.

**Table S7:** Accession numbers of the genomes used to construct phylogenetic tree in Figure 1.

| **Strain** | **Accession number** |
| --- | --- |
| *P. aeruginosa* DK2 | NC_018080.1 |
| *P. aeruginosa* PA14 | NC_008463.1 |
| *P. aeruginosa* PAO1 | NC_002516.2 |
| *P. aeruginosa* B136-33 | NC_020912.1 |
| *P. aeruginosa* LESB58 | NC_011770.1 |
| *P. aeruginosa* M18 | NC_017548.1 |
| *P. aeruginosa* NCGM2.S1 | NC_017549.1 |
| *P. aeruginosa* RP73 | NC_021577.1 |
| *P. aeruginosa* YL84 | NZ_CP007147.1 |
| *P. aeruginosa* MTB-1 | NC_023019.1 |
| *P. aeruginosa* Carb01-63 | NZ_CP011317.1 |
| *P. aeruginosa* VRFPA04 | NZ_CP008739.1 |
| *P. aeruginosa* PA_D1 | NZ_CP012585.1 |
| *P. aeruginosa* DHS01 | NZ_CP013993.1 |
| *P. aeruginosa* SJTD-1 | NZ_CP015877.1 |
| *P. aeruginosa* AES-1R | NZ_CP013680.1 |
| *P. aeruginosa* ATCC 27853 | NZ_CP015117.1 |
| *P. aeruginosa* USDA-ARS-USMARC-41639 | NZ_CP013989.1 |
| *P. aeruginosa* SCV20265 | NC_023149.1 |
| *P. aeruginosa* PA1 | NC_022808.2 |
| *P. aeruginosa* NCTC10332 | NZ_LN831024.1 |

